# Microbiota- and diet-specific T cells become T_regs_ by default

**DOI:** 10.64898/2026.08.25.747099

**Authors:** Jeffrey J. Bunker, Jamie E. Blum, Xiandong Meng, Eugenell Mae Lopez, Allison M. Weakley, Ashley V. Cabrera, Steven Higginbottom, Ryan Kong, E.A. Schulman, Elizabeth S. Sattely, James J. Moon, Michael A. Fischbach

## Abstract

CD4^+^ T cells recognize antigens from microbiota, diet, and pathogens via T cell receptors (TCRs) and orchestrate immunity by differentiating into tolerogenic regulatory (T_reg_) or pro-inflammatory effector (T_eff_) lineages (e.g. T_H_1 or T_H_17) (*1*). Dysregulation of these responses underlies numerous gastrointestinal inflammatory and infectious diseases (*2–6*). The prevailing paradigm suggests that individual microbes and dietary antigens drive distinct cell fates (e.g., segmented filamentous bacteria [SFB] induce T_H_17 cells (*7*) whereas *Helicobacter hepaticus* (*8*) and diet (*9*) induce T_regs_). However, the generality of this model is uncertain: several key organisms are atypical, and foundational studies often omitted a complex microbiome or a diverse polyclonal TCR repertoire. Here we develop a high-throughput pipeline to screen hundreds of TCRs from mice colonized from birth with a 116-strain human microbiota (hCom2v), demonstrating that TCRs recognizing microbiota or dietary antigens are overwhelmingly enriched in the induced T_reg_ (iT_reg_) lineage. Endogenous CD4^+^ T cells specific for these antigens adopt a uniform iT_reg_ phenotype *in vivo*, both in hCom2v-colonized and conventional mice. This baseline tolerance is robust to acute inflammation but breaks down following a “two-hit” combination of inflammation and genetic susceptibility, allowing T_eff_ to emerge against otherwise T_reg_-restricted antigens. These data support a revised paradigm in which antigen-specific T_reg_ induction is the default response to foreign antigens in the healthy gut, and effector responses are an exception reflecting a perceived threat. Reframing gastrointestinal immunity as a tolerance-first system provides a framework for understanding inflammatory disease pathogenesis and suggests that therapeutic strategies should aim to restore a T_reg_-predominant baseline.

## INTRODUCTION

In the gastrointestinal mucosa, the immune system encounters an enormous diversity of antigens derived from microbiota, food, and pathogens. While tolerance to harmless antigens is crucial to prevent pathologies such as inflammatory bowel disease (*2*), celiac disease (*5*), and food allergies (*4*), failure to mount an appropriate response to a pathogen can be fatal. Discriminating between helpful and harmful antigens requires resolving inherent ambiguities between these categories: commensal bacteria harbor factors such as lipopolysaccharides and peptidoglycan that are also found in pathogens (*10*), food can contain pathogens or their toxins (*11*), and major pathogens can colonize the gut without causing disease (*6*). As a result, typical mechanisms of self-nonself discrimination by the innate immune system may be insufficient to reliably detect genuine threats. Antigen-specific CD4^+^ T cell responses are thought to play a critical role in determining whether the response to a given antigen is tolerogenic or pro-inflammatory (*1*), but the organizing principles that govern these decisions remain poorly understood.

Bacterial species from the gut microbiome, as well as dietary antigens, can elicit potent CD4^+^ T cell responses during homeostasis, in the absence of overt inflammation. The current paradigm suggests that individual microbial species and dietary components elicit specific T_eff_ or T_reg_ responses: for example, SFB induce T_H_17 (*7*), *Klebsiella* spp. induce T_H_1 (*12*), *Akkermansia muciniphila* induce T_FH_ (*13*), and *Clostridium* spp. (*14*), *H. hepaticus* (*8*), and dietary antigens (*9*) induce T_regs_.

Three limitations of prior studies raise questions about whether this paradigm generalizes to a natural gut environment, where an endogenous polyclonal TCR repertoire must recognize and respond to a vast array of microbial and dietary antigens. (*i*) Some studies employed gnotobiotic mice colonized as adults with one or a few organisms. However, a given organism can elicit different responses in this setting than in a complex community (*13*, *15*), and these models bypass the weaning period, a critical window for inducing tolerogenic responses early in life (*16*, *17*). (*ii*) Others relied on model antigens and TCRs (such as the OT-II TCR and its antigen, chicken ovalbumin [OVA] (*18*)), which fail to capture key properties of natural antigens (*19*) as well as the vast diversity and low clonal frequency of the endogenous TCR repertoire (*20*). (*iii*) Several organisms central to this paradigm are atypical: SFB adheres directly to epithelial cells of the terminal ileum, where it stimulates an immune response distinct from any known commensal (*21*); *Klebsiella* spp. were opportunistic pathogens isolated from oral flora rather than the gut (*12*); and *H. hepaticus* is a context-dependent pathogen that elicits chronic colitis, hepatitis, and tumorigenesis in multiple mouse strains (*22*). Although these tools have provided tractable experimental systems, they may not reflect how the immune system responds to more typical commensals. To our knowledge, no study has systematically evaluated CD4^+^ T cell specificity and its relation to lineage fate across the endogenous polyclonal repertoire following natural acquisition of a complex microbiome.

### Modeling physiologic colonization in progeny of hCom2v-colonized mice

To enable large-scale analysis of antigen-specific intestinal CD4^+^ T cell responses in a natural setting, we developed a murine system that recapitulates vertical acquisition of a complex microbiota. We recently developed a complex defined community consisting of the most prevalent strains in the healthy human gut microbiome, termed hCom2 (*23*); here, we utilized a 116-strain variant, hCom2v, which lacks three strains with idiosyncratic growth characteristics. In prior work, we used germ-free mice colonized by variants of this community to explore the specificity of microbiome-specific CD4^+^ T cells in the intestine (*24*). However, this model had two shortcomings: first, since these mice were adults colonized for only two weeks with the defined community, they are not a faithful model of physiologic T cell responses. Second, only about 90 T cell clonotypes were assayed, selected from a variety of cell types across two community variants, making it difficult to draw broad conclusions about how CD4^+^ T cell specificity relates to lineage fate.

We colonized germ-free (GF) adult C57BL/6 (B6) mice with hCom2v, allowed two weeks for equilibration, and then bred them to generate F1 progeny colonized with hCom2v from birth (**Fig. 1a**). Metagenomic sequencing of small intestinal (SI) and colonic contents detected 99 strains in hCom2v F1 mice (**Fig. 1a and Supplementary Fig. 1a**) spanning more than six orders of magnitude in relative abundance. Community composition and strain detection were similar to prior data from short-term hCom2-colonized adult mice (*23*). Although most strains were present in both the SI and colon, the phylum Bacteroidota was biased toward the colon whereas other phyla were typically evenly distributed or SI-biased (**Supplementary Fig. 1b**). hCom2v-colonized mice are fed a standard chow diet consisting of seven protein-containing components, which can be leveraged to identify T cell clones specific for natural dietary antigens (*19*).

**Figure 1:**
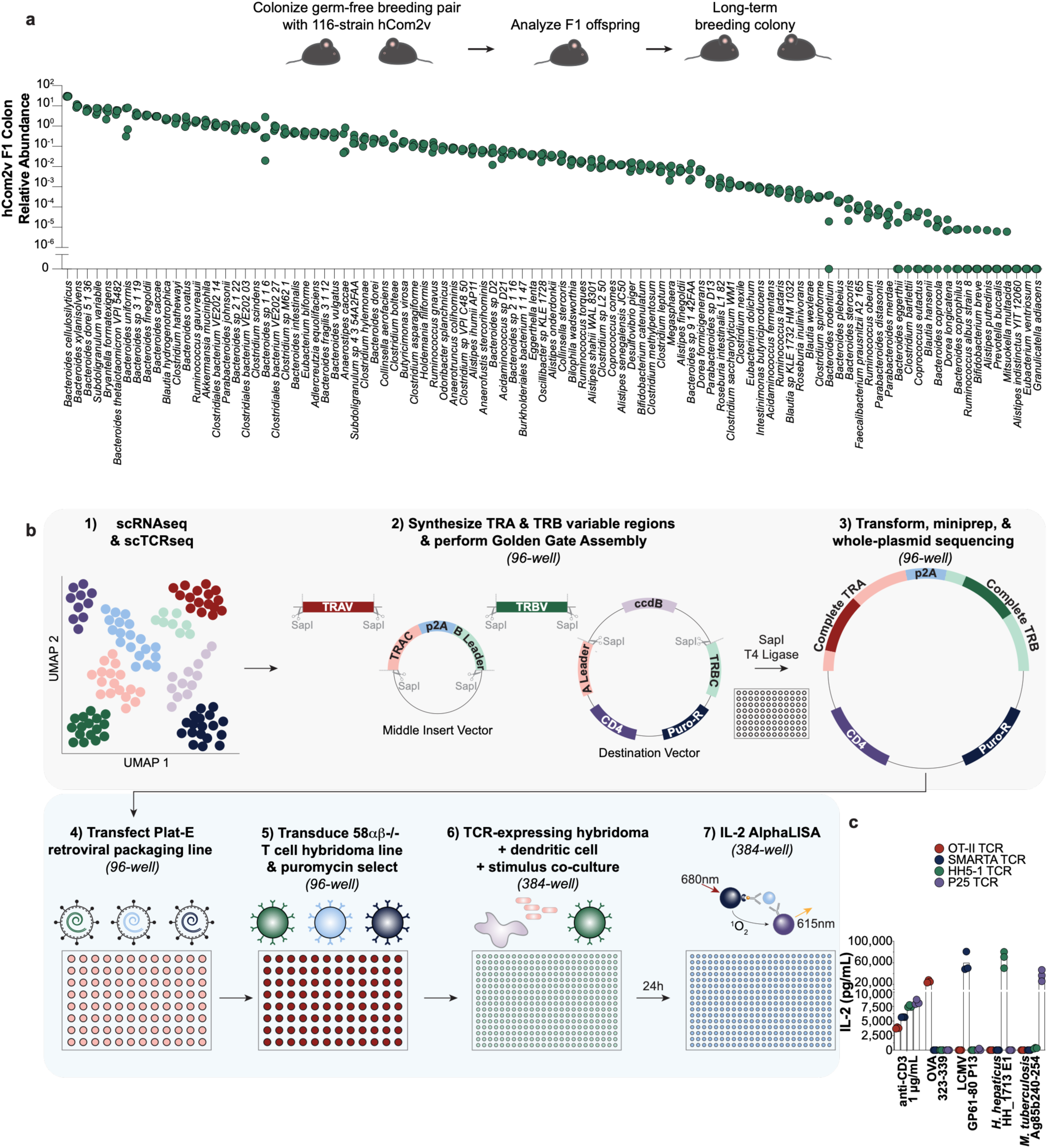
High-throughput mapping of CD4^+^ T cell specificity for microbiota and diet in naturally colonized mice. (**a**) Schematic of the hCom2v model of natural vertical transmission *(top)*. GF adults were gavaged with the 116-strain hCom2v, allowed to equilibrate for 2 weeks, then bred; their F1 offspring, colonized with hCom2v from birth, were analyzed in initial experiments and subsequently used to propagate a gnotobiotic breeding colony. Relative abundance of individual strains in the colon of hCom2vF1 mice by metagenomic sequencing *(bottom)*. Each dot represents an individual mouse (*n* = 4) and only strains detected in the SI or LI of at least one mouse are shown. (**b**) Schematic of our high-throughput pipeline linking T cell fate to TCR specificity. Variable regions from TCRs of interest identified by scTCRseq are synthesized in 96-well format and cloned into retroviral expression vectors using a 96-well Golden Gate assembly strategy. After 96-well transformation into *E. coli*, miniprep, and whole-plasmid sequencing, correctly assembled constructs for each TCR are selected and concentrations normalized. Plasmids are transfected into Plat-E retroviral packaging cells and virus-containing supernatant is used to transduce the TCR-deficient 58αβ^-/-^ T cell hybridoma line in 96-well format; transduced clones are selected with puromycin and cryopreserved. TCR specificity is measured in a 384-well assay in which TCR-expressing hybridomas, primary DCs, and a stimulus are co-cultured for ∼24 hours. IL-2 secretion into the supernatant is measured by AlphaLISA as a readout of TCR activation. (**c**) Four previously characterized TCRs (OT-II, SMARTA, HH5-1, and P25) were cloned and expressed in hybridoma cells and assayed by co-culture against their known cognate peptide, that of the other three TCRs, or soluble anti-CD3ε as a positive control. IL-2 secretion into the culture supernatant was measured by AlphaLISA. Each TCR was assayed against each stimulus in triplicate, with individual dots depicting each replicate and bars indicating the mean.

We subsequently established a breeding colony derived from hCom2v F1 offspring. Metagenomic sequencing of colonic contents or feces from distinct cages, sampled across more than two years, revealed a stable community with 90-115 detectable strains (**Supplementary Fig. 2**). Most strains maintained a consistent relative abundance over time; however, about one year after establishing the colony we detected a sustained increase in abundance of *Blautia wexlerae*, *Bacteroides rodentium*, *B. eggerthii*, and *B. coprocola* with a corresponding decrease in *B. uniformis* and *B. xylanisolvens* (**Supplementary Fig. 2**). Thus, hCom2v-colonized mice recapitulate the defined, complex hCom2 community while enabling natural, physiologic exposure to the immune system and stable propagation over years.

### Characterizing CD4^+^ T cell responses in hCom2v F1 mice

To comprehensively characterize CD4^+^ T cell responses in hCom2v F1 mice, we sorted CD4^+^ T cells isolated from the lamina propria (LP) of the SI and large intestine (LI), Peyer’s patches (PP), and spleen, pooling each tissue across four hCom2v F1 or four GF mice, and performed single-cell RNA sequencing (scRNAseq) and TCR sequencing (scTCRseq). We identified 18 cell clusters across all tissues and groups (**Supplementary Fig. 3a**), annotated them based on marker gene expression (**Supplementary Fig. 3b-c**), and assigned them to 17 cell types (**Supplementary Fig. 4**). These include all expected major populations: naïve T cells, multiple T_eff_ subsets (T_H_1, T_H_2, T_H_17, and T_FH_), natural T_regs_ (nT_regs_) and iT_regs_, and a large effector memory (T_EM_) population which generally lacked lineage-defining transcription factors and cytokines but expressed markers such as *Cxcr6*, *Rgs1*, *Bhlhe40*, and *P2rx7* associated with resident memory T cells (*25*). The spleen contained predominantly naïve T cells, whereas most T_eff_ and T_regs_ were concentrated in the SI and LI LP and T_FH_ were enriched in the PP, as expected.

scTCRseq revealed large clonal expansions among T_eff_, particularly in the SI LP, that were greater in magnitude in hCom2v-colonized than GF mice (**Supplementary Fig. 5a-b**). Small clonal expansions were observed among T_regs_, whereas naïve T cells were predominantly single clones, as expected (**Supplementary Fig. 5a-b**). Clonal overlap between tissues was apparent within both the hCom2v and GF cohorts (**Supplementary Fig. 5c**). Notably, clonal overlap was substantial among T_eff_ lineages, but minimal between T_eff_ and T_reg_ lineages (**Supplementary Fig. 5d**). This segregation of the repertoire into largely non-overlapping T_reg_ and T_eff_ compartments is consistent with studies in specific-pathogen-free (SPF) mice using a fixed transgenic TCRβ chain to limit repertoire diversity (*26*), but contrasts with our prior study in adult mice colonized with hCom variants for two weeks in which extensive clonal sharing between T_reg_ and T_eff_ lineages was observed (*24*), suggesting that developmental timing or duration of microbial exposure may influence clonal partitioning into T_eff_ and T_reg_ fates.

### A high-throughput pipeline to map intestinal CD4^+^ T cell specificity in hCom2v F1 mice

The endogenous CD4^+^ T cell repertoire of an individual mouse contains millions of unique TCRs, each capable of recognizing many individual peptides presented on major histocompatibility complex class II (MHCII) molecules (*27*). Thus, understanding how TCR specificity relates to lineage fate requires scalable approaches to dissect this diversity. With this challenge in mind, we developed a pipeline that would enable us to sample large numbers of TCRs from intestinal T cells, express them in T cell hybridoma lines, and assay their specificity against each of the microbial strains and dietary components present in hCom2v-colonized mice (**Fig. 1b**). It consists of: (*i*) performing scRNAseq and scTCRseq on intestinal CD4^+^ T cells; (*ii*) cloning individual TCRs into retroviral expression vectors using a custom 96-well Golden Gate assembly method; (*iii*) 96-well transfection into retroviral packaging cells; (*iv*) retroviral transduction of a TCR-deficient T cell hybridoma line followed by puromycin-mediated selection of transduced clones; and (*v*) a highly sensitive and specific co-culture assay to determine antigen specificity in 384-well format. This creates a workflow that is compatible with high-throughput screening and, because each TCR is recovered along with its single-cell transcriptome, directly links TCR specificity to lineage fate. It also overcomes a major limitation of prior approaches, in which *ex vivo* expansion of T cell clones can lead to biased selection and distorted clonal frequencies (*28*). Using this pipeline, we screened ∼400 unique TCRs in over 50,000 individual assays against the defined set of microbial strains and dietary components to which hCom2v F1 mice are exposed.

### Cloning and expressing TCRs at high throughput

We developed a custom Golden Gate assembly strategy to clone paired *TRA* and *TRB* variable regions identified by scTCRseq into individual retroviral expression vectors in 96-well format (**Fig. 1b, Supplementary Fig. 6a, and Supplementary Table 1**; see Methods). Assembly was highly efficient, allowing 96 TCRs to be cloned at a >90% success rate in ∼4 half-days of hands-on work (**Supplementary Fig. 6b-d**), and could be performed iteratively to clone hundreds of TCRs. Assembled vectors were transfected into retroviral packaging cells (*29*) and the resulting virus-containing supernatant was used to transduce a TCR-deficient T cell hybridoma line (*30*) in 96-well format, followed by puromycin selection and cryopreservation (**Fig. 1b**). Surface expression of retrovirally encoded TCRβ and CD4 was systematically assessed by flow cytometry just prior to cryopreservation; transduced lines were highly enriched, and surface expression was detected in all but one of >400 TCR-expressing lines (**Supplementary Fig. 6e-f**). We also expressed four control MHCII-restricted TCRs of known specificity widely used in the literature (OT-II (*18*), SMARTA (*31*), HH5-1 (*8*), and P25 (*32*), recognizing peptides from OVA, lymphocytic choriomeningitis virus [LCMV], *H. hepaticus*, and *Mycobacterium tuberculosis*, respectively); these TCRs showed expected staining with Vα and Vβ-specific antibodies for regions with commercially available reagents (**Supplementary Fig. 6g-j**), demonstrating appropriate expression and folding of diverse variable regions. This workflow therefore enables efficient generation of many arrayed cell lines, each expressing a unique TCR for downstream analysis.

### An optimized 384-well co-culture assay for screening TCR specificity

To evaluate TCR specificity at scale, we optimized a 384-well assay in which TCR-expressing hybridomas, primary dendritic cells (DCs), and a test stimulus are co-cultured (**Fig. 1b**; see Methods). As a readout of TCR activation, we measured interleukin 2 (IL-2) secretion by AlphaLISA, a miniaturizable, bead-based proximity immunoassay that requires no wash steps, increasing throughput substantially over traditional ELISA (**Fig. 1b**). We optimized assay conditions—most critically hybridoma and DC cell numbers, alongside timing and DC preparation—using the OT-II TCR side-by-side with a non-reactive control TCR from GF mice (*24*) (see Methods). Against whole OVA, the OT-II TCR produced strong IL-2 signal (several thousand pg/mL) that was titratable and abrogated by an anti-MHCII blocking antibody, whereas the non-reactive control TCR gave minimal signal (**Supplementary Fig. 7a-e**). Both TCRs responded robustly to positive control stimuli and showed minimal reactivity in the absence of stimulus (**Supplementary Fig. 7b and e**).

We next validated the assay using two groups of control TCRs. The first group comprised the OT-II and GF TCRs plus three previously characterized TCRs that recognize hCom2v strains: H1-16 and H1-24, specific for peptides from substrate binding protein (SBP) expressed by Bacillota strains, and H2-11 which recognizes a peptide from a tetratricopeptide repeat lipoprotein (TPRL) expressed by Bacteroidota strains (*24*). Assay of these TCRs against titrated OVA or normalized preparations of each strain demonstrated IL-2 signal substantially above background (thousands to tens of thousands of pg/mL) for each TCR against its cognate stimulus, minimal signal against non-cognate stimuli, and complete blockade by an anti-MHCII antibody (**Supplementary Fig. 7f**). All five TCRs responded appropriately to positive and negative control stimuli (**Supplementary Fig. 7g**). The second group consisted of the OT-II, SMARTA, HH5-1, and P25 TCRs, each assayed against their cognate peptide antigen or that of the other three TCRs. This demonstrated IL-2 signal of ∼20,000-80,000 pg/mL in response to each TCR’s cognate peptide antigen with negligible background against non-cognate peptides (**Fig. 1c**). We conclude that this optimized co-culture assay provides an exceptionally sensitive and specific readout of TCR specificity that is amenable to high-throughput screening.

We developed standardized criteria for selecting and assaying TCRs from each CD4^+^ T cell population (see Methods). Briefly, within each scRNAseq cluster, TCRs were prioritized by clone size, then by clonal overlap with other clusters, and finally at random. We set target sample sizes powered to detect population-level differences in reactivity between experimental and control populations, informed by prior data (*24*). All TCRs selected for this study and their sequences are listed in **Supplementary Table 1**. Each TCR was assayed in a primary screen against 128 stimuli by co-culture: three controls, pooled cecal and colonic contents from hCom2v-colonized mice, each of the 116 strains in hCom2v, and the chow pellet and its seven protein-containing components (**Supplementary Fig. 8a**). TCR-stimulus pairs that reacted above threshold in the primary screen were re-tested in an independent validation assay to establish true positives (**Supplementary Fig. 8b**). Complete reactivity data for all TCRs and stimuli are provided in **Supplementary Table 2**. Applying this framework across all major CD4^+^ T cell populations allowed us to systematically map the relationship between microbial and dietary antigen recognition and lineage fate.

### Induced T_regs_ frequently recognize microbiota and dietary antigens

We first profiled the reactivity of 54 LI iT_reg_ TCRs from hCom2v-colonized mice. We found that 25 TCRs (46%) reacted against either microbiota or dietary components (**Fig. 2a-b**). Fifteen TCRs recognized microbial strains and displayed a variety of reactivity patterns ranging from single-strain to multi-strain recognition. An additional 10 TCRs recognized dietary components, typically either corn or wheat. Some TCRs reacted against cecal and colonic contents from hCom2v-colonized mice in addition to one or more individual strains or dietary components, whereas in other cases reactivity was apparent only at the single-strain or single-component level. The remaining non-reactive clones may recognize self-antigens (*33*, *34*) or antigens not captured in the screen (e.g. due to restricted expression). Notably, clonal expansion in the LI iT_reg_ lineage was modest: the largest clonotype comprised only seven cells and recognized corn (**Fig. 2a**). We also found many microbiota- and diet-reactive clones in a panel of 45 SI iT_reg_ TCRs from hCom2v-colonized mice, although their frequency was somewhat reduced compared to the LI (**Fig. 2b and Supplementary Fig. 9a**). Together, these data indicate that the tolerogenic iT_reg_ compartment constitutes a diverse TCR repertoire targeted against a wide array of microbial and dietary antigens.

**Figure 2:**
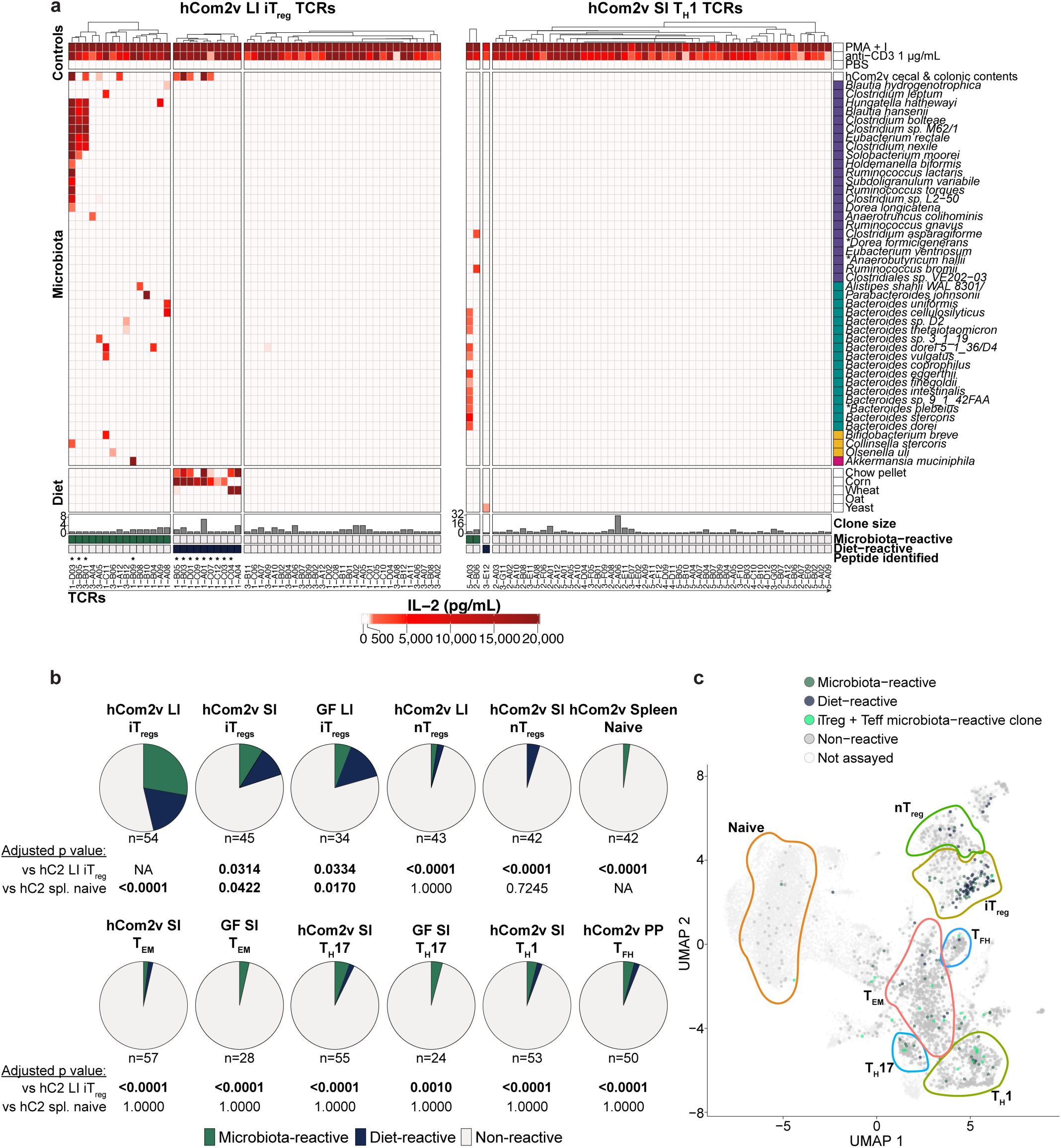
Microbiota- and diet-reactive TCRs predominantly localize to the induced T_reg_ lineage. (**a**) Heatmaps depicting reactivity patterns of 54 LI iT_reg_ or 53 SI T_H_1 TCRs from hCom2v-colonized mice. Each column represents a unique TCR with its identifier listed on the x-axis; each row represents an individual stimulus. Dendrograms depict hierarchical clustering of columns by Spearman correlation distance with Ward’s D2 method. Cell color indicates IL-2 concentration in pg/mL measured by AlphaLISA following co-culture assay. Each TCR was screened against a panel of 128 stimuli, and control stimuli plus those against which at least one TCR had a validated reactivity in any panel are shown. Bottom annotations indicate the clone size for each TCR within the cluster, whether TCRs were scored as microbiota-reactive (green), diet-reactive (blue), or non-reactive (gray), and asterisks indicate TCRs for which a peptide epitope was identified. Rows with an asterisk contain some missing data points. See Supplementary Table 2 for the complete dataset and Methods for the complete screening workflow. (**b**) Summary plots depicting the fraction of TCRs within the indicated populations scored as microbiota-reactive (green), diet-reactive (blue), or non-reactive (gray). The number of TCRs in each panel is indicated below the summary plot. P values compared to the indicated reference populations were calculated by Fisher’s exact test and adjusted for multiple comparisons with the Benjamini-Hochberg method. (**c**) Overlay of TCR reactivity data onto the scRNAseq uniform manifold approximation and projection (UMAP) plot. Approximate outlines of key clusters are indicated with colored lines—all annotated clusters are shown in Supplementary Fig. 4. TCRs scored as microbiota-reactive (green), diet-reactive (blue), or non-reactive (gray) are indicated, as are cells not assayed (faint gray) and a notable microbiota-reactive clone shared between SI iT_regs_ and multiple T_eff_ subsets (sea green).

We next derived panels of TCRs from two key control populations: 34 TCRs from GF LI iT_regs_, which have never been exposed to exogenous microbes, and 42 splenic naïve T cell TCRs from hCom2v-colonized mice, which represent random TCRs that have undergone positive selection in the thymus but no peripheral activation or expansion (**Fig. 2b and Supplementary Fig. 9a**). Just two GF LI iT_reg_ TCRs (6%) and one naïve splenic TCR (2%) recognized strains in hCom2v, substantially lower than the 28% observed among hCom2v-colonized LI iT_regs_. This indicates a baseline 2-6% rate of incidental reactivity against the thousands of microbially-derived antigens present in our screen. Further, 15% of GF LI iT_reg_ TCRs recognized dietary components, comparable to the 19% observed among hCom2v-colonized LI iT_regs_, whereas zero diet-reactive TCRs were observed among splenic naïve T cells. The overall microbiota and dietary reactivity distributions of both populations were significantly different from one another and from hCom2v-colonized LI iT_regs_ (**Fig. 2b**). Thus, LI iT_regs_ in hCom2v-colonized mice are substantially enriched in reactivity toward microbiota compared to the equivalent population from GF mice, and significantly enriched in both microbiota and dietary reactivity compared to splenic naïve CD4^+^ T cells.

Although many T_regs_ reactive against foreign antigens are thought to differentiate from naïve precursors in the periphery (*26*), other studies suggest that the LI LP contains a substantial population of thymically-derived nT_regs_ that recognize antigens from the microbiota (*35*). However, these studies utilized mice with heavily restricted TCR repertoires, where limited clonal diversity can inflate apparent overlap between intestinal T_regs_ and Foxp3^+^ thymocytes relative to the distribution in the endogenous polyclonal repertoire. To examine the reactivity of nT_regs_ toward microbiota and dietary antigens, we derived panels of 43 TCRs from LI nT_regs_ and 42 TCRs from SI nT_regs_ from hCom2v-colonized mice. We found that only ∼5% of TCRs in each population were reactive against either dietary or microbial antigens, which was not significantly different from the background in splenic naïve T cells (**Fig. 2b and Supplementary Fig. 9b**). Our data do not distinguish whether the rare foreign antigen-reactive nT_regs_ we observed are truly of thymic origin, or whether they represent iT_regs_ that overlap transcriptionally with nT_regs_ (*36*). Nonetheless, we conclude that foreign antigen reactivity is uncommon among intestinal nT_regs_ in this setting.

### Microbiota- and diet-reactive TCRs predominantly localize to the induced T_reg_ lineage

Next, we sought to determine patterns of reactivity in the T_eff_ compartment. These lineages generally contained much larger clonal expansions and substantial clonal overlap with one another but only occasional overlap with T_regs_ (**Supplementary Fig. 5a-d**). We derived panels of 57 SI T_EM_ TCRs (which contained the largest clones in our scTCRseq dataset), 53 SI T_H_1 TCRs, 55 SI T_H_17 TCRs, and 50 TCRs from PP T_FH_. Surprisingly, we found little reactivity against either microbiota or dietary antigens among any of these T_eff_ lineages (**Fig. 2a-b and Supplementary Fig. 10a-b**), whereas all TCRs responded to positive control stimuli. The reactivity distributions of the SI T_EM_ and T_H_17 populations were not significantly different from the distributions observed in 28 and 24 TCRs from the equivalent populations from GF mice, respectively (**Fig. 2a-b and Supplementary Fig. 10a-b**). None of the T_eff_ lineages were significantly enriched in reactivity above that observed in splenic naïve CD4^+^ T cells (**Fig. 2b**). The antigens driving T_eff_ clonal expansions in this system remain undefined; these TCRs may recognize self or pseudo-self antigens such as endogenous retroviruses (*37*), or their expansion may reflect antigen-independent bystander activation (*38*). Regardless, microbiota- and diet-reactive TCRs were uncommon amongst all T_eff_ lineages examined, in contrast to their enrichment among iT_regs_.

Examining ∼400 unique TCRs across 12 major CD4^+^ T cell populations, we found that only three populations were significantly enriched in reactivity above the background observed in splenic naïve CD4^+^ T cells: LI and SI iT_regs_ from hCom2v-colonized mice, and LI iT_regs_ from GF mice (which predominantly recognized dietary antigens). By contrast, no T_eff_ population showed significantly enriched reactivity above background. Overlaying our *in vitro* reactivity data onto the scRNAseq UMAP plot reinforces the view that microbiota- and diet-reactive TCRs are found predominantly in the iT_reg_ lineage; a single clone shared between SI iT_regs_ and multiple T_eff_ lineages accounted for many of the microbiota-reactive T_eff_ cells (**Fig. 2c**). These data indicate that the overwhelming majority of microbiota- and diet-reactive TCRs localize to the iT_reg_ lineage and are uncommon among T_eff_. These findings contrast with the prevailing paradigm, which predicts that within a complex community some organisms should elicit T_regs_ whereas others should elicit one or more T_eff_ lineages. Instead, they suggest that induction of antigen-specific T_regs_ may represent the default fate of foreign antigen-reactive TCRs in the healthy gut.

### Microbiota-derived SBP and dietary α-Zein are major iT_reg_ antigens

To identify peptide antigens recognized by individual TCRs, we screened them against previously described epitopes based on each TCR’s reactivity to specific microbial strains or dietary components. This strategy identified peptide antigens recognized by 17 TCRs, all of which were derived from T_regs_ (**Supplementary Fig. 11a-e**). Five TCRs reactive to multiple Bacillota strains recognized peptides from SBP (*24*): two against an N-terminal epitope and three against a C-terminal epitope (**Supplementary Fig. 11a**). Nine corn-reactive TCRs converged on a single C-terminal epitope (*19*) from the seed storage protein α-Zein (**Supplementary Fig. 11b-c**), whereas two corn-reactive TCRs did not recognize this peptide (**Supplementary Fig. 11f-g**). Two wheat-reactive TCRs recognized an epitope from gliadin (*9*, *19*) (**Supplementary Fig. 11d**). Although *Akkermansia muciniphila*-specific responses were previously suggested to induce T_FH_ or T_eff_ differentiation (*13*), other studies have associated *A. muciniphila* with iT_reg_ responses (*39*). We found no T_FH_ or T_eff_ TCRs that recognized *A. muciniphila*; a single LI iT_reg_ TCR specific for this organism was identified (**Fig. 2a**), which recognized a peptide epitope previously associated with T_FH_ responses (**Supplementary Fig. 11e**). Several TCRs that reacted against *Bacteroides* strains did not respond to previously described antigens (*24*, *40*, *41*), indicating specificity for undiscovered epitopes (**Supplementary Fig. 11h-i**). We conclude that iT_regs_ in hCom2v-colonized mice recognize a broad array of peptide antigens, among which microbiota-derived SBP and diet-derived α-Zein appear to be predominant. This likely reflects their high abundance *in vivo*: SBP- and α-Zein-reactive TCRs typically also reacted against pooled cecal and colonic contents from hCom2v-colonized mice (**Fig. 2a, Supplementary Fig. 9a-b**), consistent with SBP’s presence in many Bacillota strains in hCom2v (*24*), and α-Zein’s abundance in the chow (*19*).

### SBP-and α-Zein-specific CD4^+^ T cells are iT_regs_ *in vivo*

To validate our TCR analysis *in vivo*, we developed fluorescently labeled peptide-MHCII tetramers containing the N- and C-terminal epitopes from microbiota-derived SBP or the C-terminal epitope from diet-derived α-Zein and used these to analyze antigen-specific CD4^+^ T cells in hCom2v-colonized mice by flow cytometry (**Fig. 3a**). We detected a small population of SBP tetramer^+^ cells in the LI LP of hCom2v-colonized mice that was absent in GF controls (**Fig. 3b-c**). These cells uniformly expressed the transcription factors Foxp3 and RORγt (**Fig. 3b-c**), consistent with an iT_reg_ phenotype (*42*, *43*). SBP-specific CD4^+^ T cells expressing this same uniform iT_reg_ phenotype were also present in the SI LP of hCom2v-colonized but not GF mice (**Supplementary Fig. 12a-b**). α-Zein-specific CD4^+^ T cells were detectable in the SI and LI LP of both hCom2v-colonized and GF mice and also uniformly expressed an iT_reg_ phenotype (**Supplementary Fig. 12c-f**), as expected and consistent with a recent report (*19*). Although α-Zein-specific cells were largely Foxp3^+^ RORγt^+^ in hCom2v-colonized mice, they were a mixture of Foxp3^+^ RORγt^+^ and Foxp3^+^ RORγt^-^ phenotypes in GF mice (**Supplementary Fig. 12c and e**), suggesting that microbiota can influence T_reg_ RORγt expression independent of antigen specificity (*44*). Metagenomic sequencing confirmed colonization with hCom2v in this cohort (**Supplementary Fig. 12g**).

**Figure 3:**
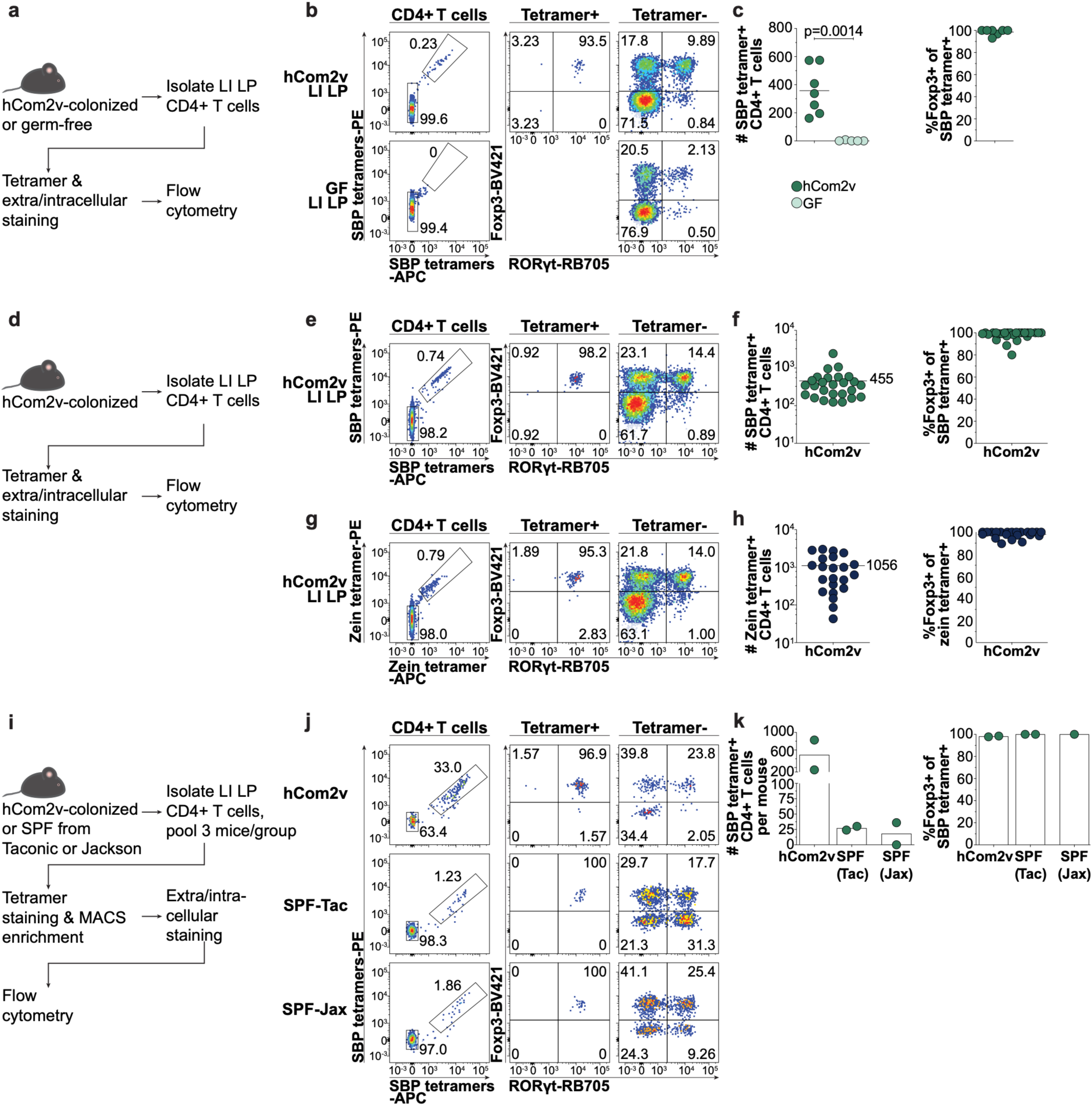
SBP- and α-Zein-specific CD4^+^ T cells are iT_regs_ *in vivo*. (**a**) Schematic depicting LI LP CD4^+^ T cell isolation from hCom2v-colonized or GF mice, tetramer and extra/intracellular staining, and analysis by flow cytometry, relevant to b-c. (**b**) Representative flow cytometry and (**c**) summary plots of SBP tetramer^+^ CD4^+^ T cell number and phenotype in hCom2v-colonized or GF mice. Each dot in the summary plots represents an individual mouse and horizontal lines indicate the mean; data compiled from two independent experiments. P value calculated by unpaired Welch’s t-test. (**d**) Schematic depicting LI LP CD4^+^ T cell isolation from hCom2v-colonized mice, tetramer and extra/intracellular staining, and analysis by flow cytometry, relevant to e-h. (**e**) Representative flow cytometry and (**f**) summary plots of SBP tetramer^+^ CD4^+^ T cell number and phenotype in hCom2v-colonized mice. Each dot in the summary plots represents an individual mouse and horizontal lines indicate the mean; data compiled from nine independent experiments and include some mice also shown in other panels of Figs. 3-4. See Supplementary Fig. 13a for full gating strategy. (**g**) Representative flow cytometry and (**h**) summary plots of α-Zein tetramer^+^ CD4^+^ T cell number and phenotype in hCom2v-colonized mice. Each dot in the summary plots represents an individual mouse and horizontal lines indicate the mean; data compiled from eight independent experiments and include some mice also shown in other panels of Supplementary Figs. 12 and 22. See Supplementary Fig. 13b for full gating strategy. (**i**) Schematic depicting LI LP CD4^+^ T cell isolation, SBP tetramer staining, and enrichment from hCom2v-colonized mice or WT B6 SPF mice purchased from Jax or Taconic (3 per group), relevant to j-k. (**j**) Representative flow cytometry and (**k**) summary plots of SBP tetramer^+^ CD4^+^ T cell number and phenotype from each group following tetramer enrichment. Each dot in the summary plots represents an individual pool of mice and bars indicate the mean; data compiled from two independent experiments—the second appears in Supplementary Fig. 14a.

SBP- and α-Zein-specific iT_regs_ were reproducibly detected across individual hCom2v-colonized mice: aggregating data from 8-9 independent experiments (*n* = 22-26 animals), we recovered a mean of 455 SBP-specific and 1,056 α-Zein-specific CD4^+^ T cells from the LI LP per mouse, which were uniformly iT_regs_ in all cases (**Fig. 3d-h**; see **Supplementary Fig. 13a-b** for full gating strategy). Together, these findings corroborate our TCR analysis and further demonstrate that CD4^+^ T cells specific for major microbiota and dietary antigens differentiate into iT_regs_ *in vivo*.

Because our TCR analysis and *in vivo* validation experiments were conducted in mice colonized with a defined community of human isolates, we next asked whether commensal and diet-specific CD4^+^ T cells adopt a similar phenotype in the context of a complex (native) murine microbiota. We leveraged the fact that SBP is widely conserved among strains of the phylum Bacillota, including those that colonize the murine gut, and that mice from the major commercial vendors Jackson Laboratory (Jax) and Taconic Biosciences (Tac) are fed a similar corn-containing chow diet (*19*). Analyzing intestinal CD4^+^ T cells from wild-type (WT) SPF B6 mice purchased directly from Jax or Tac by tetramer staining and flow cytometry, we readily detected α-Zein-specific CD4^+^ T cells in the SI and LI LP of individual mice that were similar in number and phenotype to those in hCom2v-colonized animals (**Supplementary Fig. 13c-d**), consistent with a recent report (*19*). By contrast, we were unable to reliably detect SBP-specific T cells in individual Jax or Tac mice, suggesting that they were either absent or too rare to distinguish from background without enrichment.

To clarify this point, we performed magnetic enrichment of tetramer^+^ cells from the LI LP of hCom2v-colonized, Jax, or Tac mice (three mice pooled per group; **Fig. 3i**), which we estimated would increase the frequency of tetramer^+^ cells by 50-100-fold (*45*). Following enrichment, we detected approximately 490 SBP tetramer^+^ cells per mouse in hCom2v-colonized animals across two independent experiments, which displayed an iT_reg_ phenotype as expected (**Fig. 3j-k, Supplementary Fig. 14a**). Strikingly, enrichment also revealed a small population of SBP tetramer^+^ cells (about 15-30 per mouse), all exhibiting a uniform iT_reg_ phenotype, in Tac animals in both experiments and Jax in one of two (**Fig. 3j-k**), with the second Jax experiment yielding zero SBP tetramer^+^ cells (**Supplementary Fig. 14a**). Metagenomic sequencing demonstrated that microbiota from hCom2v-colonized, Jax, and Tac animals differed in composition (**Supplementary Fig. 14b**); for example, SFB was detected only in Tac mice (**Supplementary Fig. 14c**), as expected (*7*). Importantly, we identified SBP homologs containing the N- and C-terminal epitopes in assembled metagenomes from all Jax and Tac mice examined—typically in organisms from the families Oscillospiraceae or Lachnospiraceae (**Supplementary Fig. 14c**), consistent with SBP’s known phylogenetic distribution (*24*). Therefore, CD4^+^ T cells specific for SBP or α-Zein adopt a uniform iT_reg_ phenotype *in vivo* even in the context of a conventional murine microbiota.

### Microbiota-specific T_regs_ maintain their identity and expand during acute DSS colitis

Our findings support a model in which most microbiota- and diet-reactive TCRs become iT_regs_ in the healthy gut. Loss of tolerance to microbiota is thought to underlie the pathogenesis of IBD (*46*), although the mechanisms by which this might occur remain elusive. Leveraging our ability to track antigen-specific T_regs_ *in vivo* using peptide-MHCII tetramers, we asked whether antigen-specific tolerance is maintained or breaks down during inflammation. Dextran sodium sulfate (DSS)-induced colitis is a widely used model of IBD in which oral administration of DSS leads to colonocyte toxicity and denudation of the mucus layer, causing acute barrier breach and inflammatory colitis (*47*). Although some studies have noted T_eff_ expansion in the LI LP following DSS treatment (*48*, *49*), others have reported predominant T_reg_ expansion with minimal T_eff_ differentiation (*50*). While T_regs_ are known to suppress colonic inflammation in this setting (*51*), the antigen specificity of these expanding CD4^+^ T cell populations and their phenotypic stability during inflammation are poorly understood.

Treatment of hCom2v-colonized mice with 2% DSS in their drinking water for 7 days followed by 3 days of recovery led to clinical symptoms of diarrhea, bloody stools, and weight loss (**Fig. 4a**) as well as colonic shortening (**Supplementary Fig. 15a**) compared to untreated controls, as expected. While the hCom2v community showed minimal change over the course of the experiment in untreated mice (**Supplementary Fig. 15b**), we observed significant changes in the abundance of 52 strains following DSS treatment compared to pre-treatment fecal samples (false discovery rate (FDR) <0.05; **Supplementary Fig. 15c**). Although most fold changes were small, a few strains shifted more substantially, including notable decreases in *B. wexlerae* and *B. rodentium* and an expansion of *Ruminococcus gnavus*.

**Figure 4:**
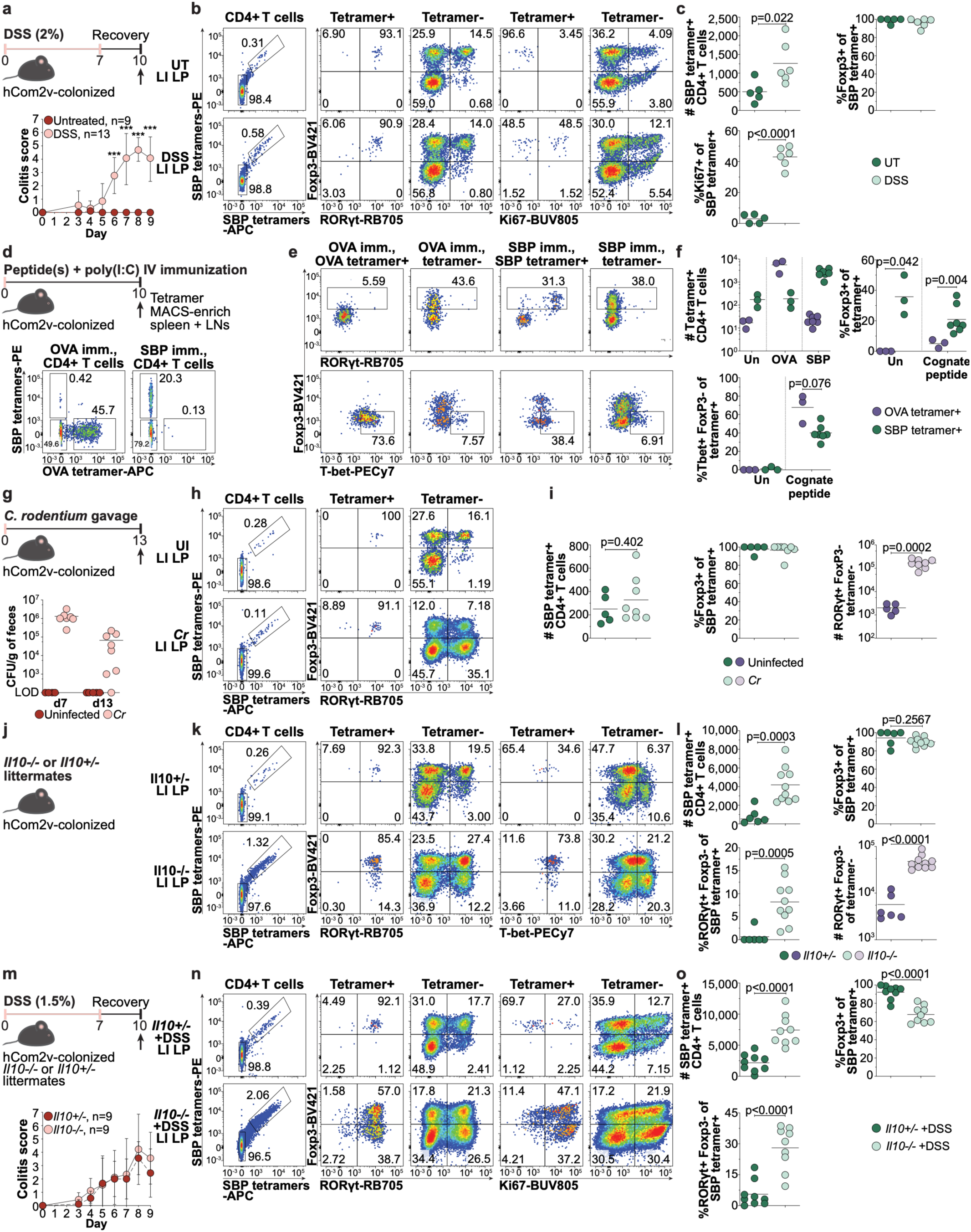
Antigen-specific tolerance is maintained during acute inflammation but breaks down with underlying genetic susceptibility. (**a**) Schematic depicting treatment of hCom2v-colonized mice with 2% DSS for 7 days followed by 3 days of recovery, with analysis on day 10, relevant to b-c (*top*). Colitis composite clinical scores for DSS-treated or untreated mice on the indicated days (*bottom*). Dots indicate mean and error bars standard deviation; data compiled from three independent experiments. P values calculated by Mann-Whitney test, *** indicates *P*<0.001. (**b**) Representative flow cytometry and (**c**) summary plots of SBP tetramer^+^ CD4^+^ T cell number and phenotype in DSS-treated or untreated mice; data compiled from two independent experiments. (**d**) Schematic depicting immunization of hCom2v-colonized mice with peptide(s) plus poly(I:C) adjuvant IV, with SBP and OVA tetramer enrichment from pooled spleen and LNs and analysis on day 10, relevant to e-f (*top*). Representative flow cytometry of SBP and OVA tetramer^+^ populations after enrichment (*bottom*)—see Supplementary Fig. 20a for unimmunized control staining. (**e**) Representative flow cytometry and (**f**) summary plots of SBP and OVA tetramer^+^ CD4^+^ T cell number and phenotype in mice immunized with OVA or SBP peptide(s) or in unimmunized (Un) controls; data compiled from two independent experiments. (**g**) Schematic depicting infection of hCom2v-colonized mice with *C. rodentium* by oral gavage followed by analysis on day 13, relevant to h-i (*top*). *C. rodentium* colony forming units (CFU) per g of feces from infected (*Cr*) or uninfected mice on days 7 and 13 (*bottom*); data compiled from two independent experiments. (**h**) Representative flow cytometry and (**i**) summary plots of SBP tetramer^+^ (green) or tetramer^-^ (purple) CD4^+^ T cell number and phenotype in *C. rodentium*-infected (*Cr*) or uninfected (UI) mice; data compiled from two independent experiments. (**j**) Schematic of hCom2v-colonized *Il10^-/-^* or *Il10^+/-^* littermate controls, relevant to k-l. (**k**) Representative flow cytometry and (**l**) summary plots of SBP tetramer^+^ (green) or tetramer^-^ (purple) CD4^+^ T cell number and phenotype in *Il10^-/-^* or *Il10^+/-^* mice; data compiled from two independent experiments. (**m**) Schematic depicting treatment of hCom2v-colonized *Il10^-/-^* mice or *Il10^+/-^*controls with 1.5% DSS for 7 days followed by 3 days of recovery, with analysis on day 10, relevant to n-o (*top*). Colitis composite clinical scores on the indicated days (*bottom*); dots indicate mean and error bars standard deviation; data compiled from two independent experiments. (**n**) Representative flow cytometry and (**o**) summary plots of SBP tetramer^+^ CD4^+^ T cell number and phenotype in DSS-treated *Il10^-/-^* or *Il10^+/-^* mice; data compiled from two independent experiments. In all summary plots, each dot represents an individual mouse and horizontal lines indicate the mean; P values were calculated by unpaired Welch’s t-test unless otherwise noted.

In one cohort of mice, we sorted total CD4^+^ T cells from the LI LP or mesenteric lymph nodes (mLN) of DSS-treated or untreated mice and performed scRNAseq and scTCRseq. Here, we labeled each mouse with unique oligo-conjugated hashing antibodies to facilitate comparisons between treatment groups using individual animals as replicates. We identified 18 clusters (**Supplementary Fig. 16a**), which we annotated based on marker genes (**Supplementary Fig. 16b-c**) and assigned to 14 cell types (**Supplementary Fig. 17a**). Strikingly, we observed expansion of multiple subsets of T_regs_ in the LI LP following DSS treatment, including both iT_reg_ and nT_reg_ subsets (**Supplementary Fig. 17b**), whereas T_eff_ lineages showed minimal change. We observed negligible change in either T_reg_ or T_eff_ subsets in the mLN, suggesting that T_reg_ accumulation largely occurred locally in the LI (**Supplementary Fig. 17b**). scTCRseq revealed that expanded T_reg_ populations contained small- or medium-sized clones (**Supplementary Fig. 18a-c**). Although large expanded clones were observed in the T_H_2/T_H_17 cluster following DSS treatment, this was apparent in only one of seven DSS-treated animals and was therefore considered an outlier that was unlikely to represent a general phenomenon (**Supplementary Fig. 18b**). We observed clonal overlap between tissues within each treatment group (**Supplementary Fig. 18d**), and within different T_eff_ or T_reg_ lineages but minimal overlap between these two compartments (**Supplementary Fig. 18e**). Thus, expansion of multiple subsets of LI LP T_regs_ appears to be the predominant CD4^+^ T cell response following acute DSS colitis.

DSS-expanded T_regs_ typically expressed immunosuppressive factors such as *Il10*, *Ctla4*, *Pdcd1,* and *Lag3*, and many expressed *Mki67*, suggestive of active proliferation (**Supplementary Fig. 19a**). Beyond their numerical expansion, both iT_regs_ and nT_regs_ expressed increased levels of *Foxp3* following DSS treatment (both *P*<0.0001 by pseudobulk analysis using individual mice as replicates; **Supplementary Fig. 19b**). DSS-expanded T_regs_ also expressed factors associated with tissue repair such as *Areg, Il1rl1, Itgae,* and *Ltb* (**Supplementary Fig. 19c**) (*52*) as well as chemokines and chemokine receptors (e.g. *Ccr2*, *Ccr5*, *Ccr8*, *Gpr18*, and *Cxcl10*) that may direct their localization to sites of inflammation (**Supplementary Fig. 19d**). We also observed upregulation of interferon-stimulated genes across multiple subsets following DSS treatment (**Supplementary Fig. 19e**). These data, together with studies demonstrating exacerbated DSS-induced pathology following T_reg_ depletion (*51*, *53*), suggest that LI LP T_regs_ expand and likely enforce tolerance and promote tissue repair following acute DSS-induced colitis.

We next examined SBP tetramer^+^ CD4^+^ T cells in a separate cohort of hCom2v-colonized mice following DSS treatment. Notably, these cells maintained a uniform iT_reg_ phenotype in both DSS-treated and untreated mice (**Fig. 4b-c**). SBP tetramer^+^ cells increased ∼two-fold in absolute number and contained a significantly higher fraction of Ki67^+^ cells after DSS treatment, indicating active proliferation (**Fig. 4b-c**). Together, these data indicated that microbiota-specific T_regs_ expand and stably maintain their identity, preserving tolerance following acute DSS-induced inflammation.

### Microbiota-specific T_regs_ imprint a tolerogenic bias on pro-inflammatory systemic challenge

The ability of commensal-specific T_regs_ to maintain a tolerogenic phenotype in the face of an inflammatory insult led us to explore how they would behave when confronted with a super-physiologic pro-inflammatory perturbation: systemic immunization with cognate peptide plus adjuvant. We compared hCom2v-colonized mice immunized with SBP peptides or OVA peptide as a control. In this model, mice have been previously exposed to SBP via the microbiota whereas OVA represents a novel antigen to which they are entirely naïve. We tetramer-enriched SBP and OVA-specific cells from pooled spleen and LNs ten days after intravenous immunization and examined them by flow cytometry (**Fig. 4d-f**). In unimmunized controls, about 170 SBP-specific cells were detected per mouse and consisted of a mixture of T_regs_ and naïve T cells, as compared to about 17 OVA-specific cells that showed a naïve phenotype (**Fig. 4f and Supplementary Fig. 20a**), consistent with prior estimates of the OVA-specific pre-immune repertoire (*45*). Immunization with OVA peptide led to substantial expansion of OVA-specific cells, which were largely Foxp3^-^ T-bet^+^ consistent with a T_H_1 phenotype, as expected following immunization with poly(I:C) adjuvant (**Fig. 4d-f**). We also observed expansion of SBP-specific cells following immunization with SBP peptides. However, in contrast to OVA-specific cells, a substantial fraction of SBP tetramer^+^ cells expressed Foxp3 and RORγt consistent with an iT_reg_ phenotype, whereas a smaller fraction showed a T_H_1 phenotype (**Fig. 4d-f**). Given the composition of the SBP-specific repertoire in unimmunized mice (**Supplementary Fig. 20a**), immunization-expanded T_regs_ may arise from the pre-existing T_reg_ pool, whereas the T_H_1 fraction may emerge from naïve precursors. Metagenomic sequencing confirmed colonization with hCom2v in all mice (**Supplementary Fig. 20b**). We conclude that prior microbiota-driven induction of SBP-specific T_regs_ imprints a tolerogenic bias on subsequent pro-inflammatory systemic challenge with their cognate antigen.

### Microbiota-specific T_regs_ maintain their tolerogenic identity during infectious colitis

Having found that non-infectious inflammatory stimuli expand and maintain commensal-specific T_regs_, we next turned to an infectious challenge with the pathogen *Citrobacter rodentium. C. rodentium* is a natural murine pathogen that shares core pathogenicity genes with the human pathogens enteropathogenic *Escherichia coli* (EPEC) and enterohemorrhagic *E. coli* (EHEC) (*54*). During infection, *C. rodentium* attaches directly to distal colonocytes and injects pathogenic effectors via a type III secretion system, leading to characteristic attaching and effacing lesions; disease typically consists of mild diarrhea and is self-limiting. Notably, infection induces a massive T_H_17/T_H_22 T_eff_ response in the LI LP that is required for pathogen clearance (*55*, *56*); many of these cells likely express pathogen-specific TCRs (*57*, *58*). A prior study showed that concurrent introduction of *H. hepaticus* and *C. rodentium* can skew the T_reg_-biased response to *H. hepaticus* toward a T_H_17 phenotype (*8*); however, pathogens typically invade an intestine in which commensal-specific tolerance is already established. We therefore investigated whether pre-existing commensal-specific T_regs_ would maintain or change their fate amidst substantial local infection-induced inflammation. We infected hCom2v-colonized mice by oral gavage with *C. rodentium* and analyzed mice 13 days post-infection, corresponding to the peak of the T cell response (*57*) (**Fig. 4g**). At this timepoint, *C. rodentium* remained culturable from feces, albeit at lower levels than the day 7 peak, whereas uninfected controls showed no growth (**Fig. 4g**). Colon length was modestly but not significantly shorter in infected mice (**Supplementary Fig. 21a**). Infection drove a ∼100-fold expansion of RORγt^+^ Foxp3^-^ T_eff_ cells in the LI LP (**Fig. 4h-i**), consistent with the T_H_17/T_H_22 response characteristic of *C. rodentium* infection. By contrast, SBP tetramer^+^ cells maintained their iT_reg_ phenotype and persisted at numbers comparable to uninfected controls (**Fig. 4h-i**). Metagenomic sequencing confirmed the presence of *C. rodentium* in infected mice and revealed minimal disruption to the hCom2v community (**Supplementary Fig. 21b-c**). These results demonstrate that commensal-specific T_regs_ maintain a tolerogenic phenotype even during a co-localized infection that drives massive T_eff_ differentiation.

### A minor T_eff_ population emerges against T_reg_-restricted antigens in the absence of IL-10

The preservation of antigen-specific T_reg_ responses during acute infectious or inflammatory challenges likely explains how the adaptive immune system restores homeostasis and avoids IBD during routine insults. However, IBD is linked to hundreds of genetic susceptibility loci (*2*), raising the possibility that genetic alterations might predispose to disease by altering the outcome of immune responses that would otherwise be tolerogenic. We focused on interleukin 10 (IL-10), a central anti-inflammatory regulator of gastrointestinal immune homeostasis that is produced prominently by iT_regs_ (*53*) and has pleiotropic functions including suppression of pro-inflammatory gene expression (*59*) and maintenance of T_reg_ identity (*60*). Rare human individuals with monogenic deficiency in IL-10 or its receptor develop very-early-onset IBD (*61*), polymorphisms in this pathway are associated with adult-onset IBD (*62*), and neutralizing autoantibodies against IL-10 were recently described in ∼3.5% of patients with IBD (*63*). Mice deficient in IL-10 develop spontaneous colitis whose penetrance depends on mouse strain background and microbiota composition (*64*, *65*)—particularly colonization with *H. hepaticus* (*66*), where *H. hepaticus*-specific T_regs_ are diverted to a T_H_1/T_H_17 T_eff_ phenotype in the absence of IL-10 (*8*). However, *H. hepaticus* behaves as a pathobiont in this scenario, driving an unusually strong CD4^+^ T cell response that may not represent the response to typical commensals.

To examine the phenotype of microbiota- or diet-specific iT_regs_ in the absence of IL-10, we bred germ-free *Il10^-/-^* mice into our hCom2v colony to generate *Il10^-/-^* or *Il10^+/-^* littermate controls colonized with hCom2v from birth (**Fig. 4j**). hCom2v-colonized *Il10^-/-^* animals appeared grossly healthy and exhibited no overt clinical symptoms of colitis; colon lengths were comparable to *Il10^+/-^* controls (**Supplementary Fig. 22a**). hCom2v community structure was similar between knockout and control animals (**Supplementary Fig. 22b**). We found a significant increase in the absolute number of SBP and α-Zein tetramer^+^ cells in *Il10^-/-^* mice compared to *Il10^+/-^* controls (**Fig. 4k-l and Supplementary Fig. 22c-d**). Strikingly, while most of these cells maintained their iT_reg_ phenotype in the absence of IL-10, a small population of tetramer^+^ Foxp3^-^ T_eff_ cells expressing RORγt and T-bet emerged that was absent in heterozygous controls (**Fig. 4k-l and Supplementary Fig. 22c-d**); a similar T_eff_ population was also expanded among bulk tetramer^-^ cells (**Fig. 4k-l**). We could not determine whether these antigen-specific T_eff_ represent ex-T_regs_ that formerly expressed Foxp3 (*60*) or reflect a failure to suppress *de novo* T_eff_ differentiation from naïve precursors, as the requisite fate-mapping models (*67*) are not readily available on a germ-free background and would likely be difficult to interpret for a population of this size. Regardless, we conclude that IL-10 deficiency permits the emergence of a minor T_eff_ population against typically T_reg_-restricted antigens.

### Combined IL-10 deficiency and acute inflammation lead to substantial loss of tolerance

While the emergence of T_eff_ against normally T_reg_-restricted antigens in the absence of IL-10 was striking, this population was small and the animals appeared grossly healthy, suggesting a largely subclinical phenomenon. We therefore hypothesized that a “second hit” with acute DSS-induced inflammation might precipitate a more substantial loss of antigen-specific tolerance (*68*). We treated hCom2v-colonized *Il10^-/-^* or *Il10^+/-^* littermates with 1.5% DSS in the drinking water for 7 days followed by 3 days of recovery (**Fig. 4m**). This low dose—selected to minimize excess mortality in the *Il10^-/-^* cohort (*69*)— resulted in similar clinical symptoms and colon lengths between genotypes (**Fig. 4m and Supplementary Fig. 23a**). Consistent with our prior DSS studies (**Supplementary Fig. 15**), treatment led to significant but low-magnitude changes in the abundance of many strains in hCom2v that did not segregate by genotype (**Supplementary Fig. 23b-c**), including the larger decreases in *B. wexlerae* and *B. rodentium* and expansion of *R. gnavus* seen previously. In *Il10^+/-^* +DSS controls, both SBP- and α-Zein-specific T cells maintained their iT_reg_ phenotype and showed evidence of proliferation based on Ki67 staining, as expected (**Fig. 4n-o and Supplementary Fig. 23d-e**). Remarkably, we observed substantially expanded populations of SBP- and α-Zein-specific cells in *Il10^-/-^*+DSS mice, characterized by the outgrowth of a significant, highly proliferative (Ki67^hi^) population of Foxp3^-^ RORγt^+^ T_eff_ together with iT_regs_ (**Fig. 4n-o and Supplementary Fig. 23d-e**). Thus, acute inflammation in a genetically susceptible background led to substantial loss of tolerance to normally T_reg_-restricted antigens.

## DISCUSSION

Our data support a simple organizing principle for gastrointestinal immunity: CD4^+^ T cells specific for microbiota or dietary antigens become iT_regs_ by default. Although this initially appears to contrast with the prevailing paradigm, in which distinct microbial taxa drive various T_eff_ or T_reg_ fates, the two can be reconciled once T_eff_ responses are understood as exceptions that arise when an organism is recognized as a potential threat or when experimental conditions lead the organism to elicit a non-physiologic response.

Our systematic screen of ∼400 TCRs across 12 major populations revealed that the vast majority of microbiota- or diet-specific TCRs in hCom2v-colonized mice localized to the induced T_reg_ lineage. T_reg_-derived TCRs recognized at least 35 individual strains spanning four phyla in hCom2v and multiple dietary components. This phenomenon was not an idiosyncrasy of the hCom2v community: our peptide-MHCII tetramer analyses demonstrated that CD4^+^ T cells specific for the predominant microbiota and dietary antigens in the hCom2v system became iT_regs_ *in vivo* in conventional SPF mice. Our findings align with prior studies identifying a wide range of organisms recognized by antigen-specific T_regs_, including *Parabacteroides* and *Bacteroides* spp. (*26*, *35*), *Clostridium* spp. (*14*, *35*, *70*), and *H. hepaticus* (*8*). Over a decade ago, the observation of bulk T_reg_ expansion following colonization with an 8-member community and with DSS treatment prompted early hypotheses of a tolerogenic default (*50*). However, no study has systematically measured specificity across CD4^+^ T cell lineages in the context of natural colonization with a complex microbiome and diet; here, this revealed the unique enrichment of microbiota- and diet-reactive TCRs within the iT_reg_ lineage. Notably, microbiota- and diet-specific TCRs were typically present at small clone sizes, suggestive of a broad, low-magnitude tolerogenic response that marks benign foreign antigens across the repertoire during homeostasis. T_reg_ induction therefore emerges as the predominant outcome of microbiota and dietary antigen recognition in the healthy gut.

Given this tolerogenic default, how can the well-documented examples of microbially-driven T_eff_ induction be reconciled? The clearest case is SFB, which is known to induce T_H_17 responses and has been widely studied as a model for host-microbiome interactions (*7*). However, it is highly atypical in ways that may explain this activity: most notably, this organism adheres directly to the ileal epithelium where it stimulates local pro-inflammatory circuits (*71*) and triggers an unusual endocytic sampling mechanism in epithelial cells that is distinct from any known organism (*21*). Indeed, direct epithelial adhesion is atypical and other organisms that attach to epithelial cells (e.g. *Candida albicans, Bifidobacterium adolescentis*, or a collection of epithelial-adherent strains from humans) similarly induce T_H_17 differentiation (*72*, *73*). Epithelial adhesion is likely perceived as a potential threat given its significance in the pathogenesis of common diarrheal infections such as *C. rodentium*, EPEC, and EHEC, which attach to and efface the epithelium (*54*). Although T_H_17 cells elicited by *C. rodentium* and SFB differ in some respects (*74*), both are likely triggered by similar underlying cues related to adhesion. Notably, the principal functional effects of IL-17A—modulation of epithelial junctions, production of antimicrobial peptides, and neutrophil recruitment (*75*)—all act to counteract adherent extracellular pathogens. Thus, T_H_17-inducing organisms likely reflect immune recognition of a potential threat, with epithelial adhesion serving as a potential unifying mechanism irrespective of the organism’s true pathogenic capabilities. Consistent with this, few T_H_17 cells in hCom2v-colonized mice recognized microbiota, indicating that hCom2v lacks organisms that provoke these responses and that T_H_17 responses do not generally arise against typical commensals.

Only a handful of organisms have been reported to elicit T_H_1 responses, and each stands out as a potential pathogen or pathobiont rather than a benign commensal. For example, *Klebsiella pneumoniae* and *Klebsiella aeromobilis* (*K. aerogenes* (*76*)) stimulate T_H_1 responses in monocolonized mice (*12*), yet both are opportunistic human pathogens known to colonize the gut and cause infections such as pneumonia and urinary tract infections (*77*); indeed, the *K. pneumoniae* strain used in this study caused severe pneumonia upon inoculation into the lung (*12*). Moreover, these strains were isolated from human saliva and studied following ectopic gut mono- or oligo-colonization of germ-free mice; whether they would also induce T_H_1 responses in a natural colonization setting is unknown. Similarly, a pathogenic adherent-invasive *E. coli* strain induces a T_H_1 response (*78*). The pathobionts *H. hepaticus* and *Bilophila wadsworthia* can induce T_H_1 responses and exacerbate disease in the setting of IL-10 deficiency (with a high milk fat diet additionally required for *B. wadsworthia*) (*8*, *79*). Thus, few if any benign commensals are known to induce T_H_1 responses in the healthy gut, and the commonly cited T_H_1 inducers are more reminiscent of *bona fide* gastrointestinal pathogens such as *Listeria monocytogenes* (*80*) and *Salmonella enterica* (*81*). Furthermore, the intestinal T-bet^+^ CD4^+^ T cells that differentiate in large numbers during homeostasis may not be T_eff_ at all: recent work indicates that many are regulatory T_R_1 cells that arise independently of microbiota and are modulated by dietary composition (*82*). Consistent with this, we found few if any T_H_1 cells reactive against microbiota or dietary antigens in hCom2v-colonized mice, indicating that typical commensals do not elicit these responses.

T_FH_ are predominant in PP and mLN, where they function to promote T cell-dependent IgA responses (*83*). As with the T_H_17 and T_H_1 lineages, only a small number of organisms—namely, SFB, *H. hepaticus*, and *A. muciniphila—*are known to induce antigen-specific T_FH_ responses (*8*, *13*). Indeed, sequencing of IgA-bound commensals in T cell-deficient mice established that T-independent responses are sufficient to coat most commensals with IgA, with SFB representing a rare T-dependent exception (*84*, *85*). In support of these findings, we found very few T_FH_ TCRs reactive against microbiota or diet in hCom2v-colonized mice. While SFB and *H. hepaticus* are atypical for reasons discussed elsewhere, *A. muciniphila* warrants special consideration as a nuanced case. Prior work showed that *A. muciniphila* elicited predominant T_FH_ responses in a minimal 8-member community but other fates—including T_regs_— when introduced with a complex, undefined microbiota (*13*). Other studies have likewise associated *A. muciniphila* with T_reg_ induction (*39*). The *A. muciniphila* strain in hCom2v is a type strain that may differ meaningfully from the primary isolate studied previously (*13*). Nonetheless, we found no PP T_FH_ reactive against *A. muciniphila* in hCom2v-colonized mice; only a single TCR specific for this organism was identified, which was derived from an iT_reg_ and recognized the same epitope previously associated with T_FH_ differentiation (*13*).

The paucity of microbiota- and diet-specific TCRs among T_eff_ lineages indicates that their homeostatic expansion must be driven by alternative cues in the absence of pathogens or pathobionts. Our co-culture assay showed exceptional sensitivity and identified dozens of reactivities among T_reg_-derived TCRs, indicating that the low T_eff_ reactivity was not merely a failure of detection. While the precise targets of these homeostatic T_eff_ populations remain to be defined, our data establish that reactivity against typical commensals and dietary antigens is uncommon within these lineages; reactivity did not exceed that observed in random TCRs from splenic naïve T cells. Notably, most CD4^+^ T cell subsets are present in GF and antigen-free mice (*44*), albeit at reduced magnitude, as well as in the human fetal intestine (*86*). These baseline T_eff_ may recognize self or pseudo-self antigens such as endogenous retroviruses or may arise through antigen-independent bystander mechanisms, among other possibilities. We hypothesize that compartmentalization of T_reg_ versus T_eff_ fates is enforced by the nature of the antigen-presenting cell (APC) that stimulates a given response: whereas T_reg_ induction during homeostasis is driven by specialized RORγt^+^ APCs, organisms exhibiting features that mark them as potential threats—such as epithelial adhesion or invasion—may instead be sampled by alternative APCs, such as classical DCs, that subsequently instruct T_eff_ differentiation (*87*).

Our findings highlight the importance of capturing microbiota complexity and natural vertical transmission in gnotobiotic models. Much of our understanding of host-microbiome interactions derives from mono- or oligo-colonized mice, an approach that has yielded fundamental mechanistic insights. Yet, colonization at non-physiologic abundances, in the absence of competing taxa, or outside the developmental window in which tolerance is typically established can elicit immune responses that would not occur in a natural context. Consistent with a key role for timing and duration of microbial exposure, SBP-specific TCRs recovered from mice colonized as adults for two weeks with related defined communities were distributed across both T_eff_ and T_reg_ lineages (*24*), whereas the SBP-specific TCRs we recovered following colonization from birth were restricted to the T_reg_ lineage—a phenotype we confirmed *in vivo* via tetramer staining. RORγt^+^ APCs and goblet cell-associated antigen passages are known to peak in gut tissues in early life, and may contribute to the enhanced propensity for tolerogenic responses observed in this window (*16*, *17*, *88*). These considerations may help reconcile our findings with prior studies reporting T_eff_ induction by individual organisms, many of which were characterized in reduced-complexity or adult-colonization settings. Ultimately, a key frontier will be to apply systematic TCR screening approaches to hosts with diverse MHC alleles and undefined microbiota, including humans.

The commensal- and diet-specific iT_reg_ response was remarkably resilient during acute colitis, illuminating how this default response is beneficial during inflammation: pre-existing iT_regs_ specific for benign commensal or dietary antigens maintain their phenotype and enforce tolerance, having already marked these antigens as likely non-harmful. These findings may explain how the immune system avoids IBD despite the myriad inflammatory insults encountered over a typical human lifetime. We observed that two hits—combined IL-10 deficiency plus acute inflammation—were required for substantial loss of antigen-specific tolerance. Given the recent discovery that 3.5% of IBD patients have neutralizing autoantibodies against IL-10 (*63*), as well as the very-early-onset IBD seen in rare individuals with monogenic deficiencies in this pathway (*61*), our findings offer a plausible mechanistic explanation for the pathological loss of tolerance that leads to disease in this subset of patients. Furthermore, this discovery may serve as a paradigm for understanding additional genetic susceptibility loci in IBD (*2*), wherein pleiotropic genetic abnormalities lead to subclinical defects in tolerogenic pathways that become clinically significant only with subsequent inflammatory exposures.

More generally, our findings provide a framework for understanding pathogenic CD4^+^ T cell abnormalities in gastrointestinal disease: T_eff_ responses against a microbial or dietary antigen indicate either that it has been recognized as a potential threat, or that an immunological defect has led the default T_reg_ response to malfunction. Accordingly, they suggest two targeted therapeutic strategies that depend on the nature of the abnormality: first, where T_eff_ responses target organisms that genuinely drive pathology, removing those organisms may be therapeutic (e.g. via antibiotics or microbial transplantation); second, when T_eff_ target antigens that are otherwise benign, as in IL-10-deficient mice following inflammatory challenge, the defect lies primarily with the host and may favor immunological interventions to restore a T_reg_-predominant baseline.

The logic of immune responses to foreign antigens in the gut is becoming clear. By defaulting to tolerogenic responses in the healthy gut, the immune system marks a broad array of benign foreign antigens with antigen-specific T_regs_. This assignment is durable: during inflammation, pre-existing T_regs_ persist and tolerance to microbiota and diet is preserved, even as T_eff_ mobilize against active threats. This architecture is robust—tolerance withstands transient inflammation—but not infallible: when a genetic defect in a tolerogenic pathway coincides with inflammation, tolerance breaks down and benign antigens are targeted by effector responses. Recognizing this logic, and further elaborating the specific ways it can fail, should inform how we understand and treat immune-mediated gastrointestinal diseases.

## ACKNOWLEDGMENTS

We thank members of the Fischbach laboratory and Stanford Microbiome Therapies Initiative for discussions and support, particularly Xianfeng Zeng, Amina Jbara, and Mikhail Levia for assistance with anaerobic microbiology and Pranav Lalgudi, Duy Nguyen, and Felicia Garcia for assistance with gnotobiotic experiments. Most cell sorting and flow cytometry data collection for this project was performed on instruments in the Stanford Shared FACS Facility (RRID: SCR_017788), including instruments obtained using NIH S10 Shared Instrument Grants S10RR027431-01 and 1S10OD026831-01. scRNAseq and scTCRseq library preparation, sequencing, and initial Cell Ranger analysis were performed at the Stanford Genomics Service Center including on instruments obtained with NIH S10 Shared Instrument grants S10OD025212 and 1S10OD021763. AlphaLISA data were collected on an instrument in the Sarafan ChEM-H High-Throughput Screening Knowledge Center (RRID: SCR_023235). OVA peptide-MHCII tetramers were obtained through the NIH Tetramer Core Facility. This work was supported by the Leona M. and Harry B. Helmsley Charitable Trust (M.A.F.), the Stanford Microbiome Therapies Initiative, NIH grants 5T32AI007502 and K99AI201798 (J.J.B.), 5R01AI175642, 5R01AI152484, and 5R01DK101674 (M.A.F.), R01DK126910 (J.J.M.), NSF grant 2125383 (M.A.F.), the Howard Hughes Medical Institute (E.S.S.), the Life Sciences Research Foundation (J.E.B.), an NSF GRFP award DGE-2146755 (R.K.), the DoW Vannevar Bush Faculty Fellowship (M.A.F.), and the Chan Zuckerberg Biohub (M.A.F.).

## AUTHOR CONTRIBUTIONS

J.J.B. and M.A.F. conceived the study. J.J.B. designed and performed most experiments and analyzed the data, supervised by M.A.F. J.E.B. contributed to the dietary antigen screening strategy and performed a subset of the peptide epitope identification assays together with R.K. and E.A.S., supervised by E.S.S. X.M., A.M.W., and A.V.C. prepared samples for 16S and metagenomic sequencing and contributed to analysis of the associated data. X.M. and J.J.B. identified SBP homologs in assembled metagenomes. E.M.L. and S.H. assisted with gnotobiotic mouse experiments. J.J.M. designed and produced peptide-MHCII tetramers and provided technical advice. J.J.B. and M.A.F. wrote the paper; all authors reviewed and approved the manuscript prior to submission.

## SUPPLEMENTARY MATERIALS

Materials and Methods

Supplementary Figures 1-23

Supplementary Tables 1-3

Supplementary References

