## Supplementary Materials for "Microbiota- and diet-specific T cells become T_regs_ by default"

**This PDF file includes:**

Materials and Methods
Supplementary Figures 1-23
Legends for Supplementary Tables 1-3
Supplementary References

#### MATERIALS AND METHODS

##### Mice

C57BL/6J mice (purchased from Jackson Laboratory) and C57BL/6NTac mice (purchased from Taconic Biosciences) were maintained in a specific-pathogen-free (SPF) facility at Stanford University. Germ-free (GF) C57BL/6 mice were from a long-term breeding colony maintained in sterile isolators at Stanford University. GF *Il10*$^{-/-}$ mice (C57BL/6 background) were obtained from the National Gnotobiotic Rodent Resource Center at The University of North Carolina at Chapel Hill and subsequently maintained in sterile isolators at Stanford University. To generate hCom2v-colonized F1 mice, the 116 strains in hCom2v were grown individually (see "hCom2v bacterial strains" below), pooled, and cryopreserved as a glycerol stock. Adult GF mice were orally gavaged with ~200 µL of the thawed stock, allowed to equilibrate for two weeks, and then males and females were housed in pairs for breeding. Initial analyses used the F1 progeny of these animals; a breeding colony was subsequently established by breeding the offspring, and the colony was propagated by establishing new breeding cages as needed. hCom2v-colonized mice were housed in individually ventilated sterile gnotobiotic cages and provided autoclaved chow (LabDiet 5K67) and water *ad libitum*. To generate hCom2v-colonized *Il10*$^{-/-}$ and *Il10*$^{+/-}$ littermate controls, GF *Il10*$^{-/-}$ mice were bred to hCom2v-colonized C57BL/6 mice to generate hCom2v-colonized *Il10*$^{+/-}$ offspring, which were then bred to GF *Il10*$^{-/-}$ mice to generate litters of hCom2v-colonized *Il10*$^{-/-}$ and *Il10*$^{+/-}$ mice. Mice were 8-10 weeks of age for most experiments and were age- and sex-matched between groups to the extent possible; DSS experiments used only male animals to minimize known sex differences in disease severity (*1*); *Il10*$^{-/-}$ experiments used animals ~12 weeks of age given previous association of this time point with a relatively high penetrance of enterocolitis (*2*). All animal procedures were approved by the Stanford University Institutional Animal Care & Use Committee (IACUC).

##### TCR screen design

Here, we describe the parameters used in the selection and screening of TCRs and their rationale. For detailed experimental methods see the corresponding section headings below.

*Sample size calculation:* Prior to the study, we performed power calculations to set reasonable target ranges for the number of TCRs to screen per population. Our prior study, which examined ~90 TCR clones spanning multiple lineages from hCom1d- or hCom2d-colonized mice two weeks after colonization of adult GF mice (*3*), observed an overall ~30% rate of reactivity against microbiota or the chow pellet. We estimated a background reactivity of 5-10% in control populations. Target sample sizes were calculated for a comparison of two independent study groups and a dichotomous primary endpoint using the ClinCalc.com Sample Size Calculator, assuming 30% reactivity in experimental populations versus 5-10% in control, $\alpha = 0.05$, power = 0.8, and a 2:1 experimental:control allocation ratio. This yielded goal sample sizes of 56-96 experimental and 28-48 control TCRs per population to guide screening, with the range reflecting the assumed 5% versus 10% background; actual sample sizes achieved per population are reported in Fig. 2b.

*TCR selection criteria:* A spreadsheet was generated tabulating TCRs from our scRNAseq/scTCRseq dataset. Within each $CD4^+$ T cell population, unique TCRs were sorted in descending order of clonal abundance, then—among TCRs of equal abundance—by descending number of populations in which the TCR appeared; remaining ties were broken randomly. Our selection strategy thus favored more highly represented clonotypes over random singletons. TCRs were selected in this order with a goal of reaching the target sample size ranges defined above. Identical TCRs that met selection criteria for multiple populations were expressed and assayed on the first occurrence and tallied for subsequent populations. All TCRs selected for this study are listed in Supplementary Table 1.

*TCR quality control (QC) criteria:* TCRs were subjected to QC at multiple stages of our pipeline with the goal of eliminating TCRs that did not clone or express properly or were otherwise non-functional. All TCR-containing plasmids were evaluated by whole-plasmid sequencing and those for which no properly assembled plasmid was obtained were excluded, unless additional colonies were screened to identify a correct clone. Following transduction into T cell hybridoma cells and puromycin selection, surface CD4 and TCRβ were analyzed by flow cytometry and TCRs that did not express both markers were excluded (the untransduced, unselected parental line served as a negative control). Finally, TCR-expressing lines were required to respond to anti-CD3ε positive control antibody stimulation (see 'Primary screen'); lines that did not produce IL-2 ≥ 500 pg/mL were excluded. A small number of cell lines that did not grow adequately in culture were excluded. For TCRs meeting selection criteria in multiple populations, the first occurrence was cloned and assayed and its QC outcome (pass or fail) applied to all instances. The QC outcomes and inclusion or exclusion of all TCRs selected for this study are listed in Supplementary Table 1.

*Primary screen*: Each TCR was screened against a panel of 128 stimuli (see Supplementary Fig. 8a for plate layout). These included three controls: phorbol 12-myristate 13-acetate (PMA) and ionomycin (Thermo 00-4970-93; 1X final concentration), soluble anti-CD3ε (145-2C11; BioLegend 100359; 1 µg/mL final concentration), and PBS-only. 116 wells contained individual strains in hCom2v that were normalized in PBS and autoclaved, and 1 well contained an autoclaved and normalized preparation of cecal and colonic contents from hCom2v-colonized mice. 8 wells contained the chow pellet or a preparation of each individual protein-containing dietary component normalized based on dry weight in PBS. Details regarding preparation of each stimulus and assay methods are described below under their corresponding headings. Stimuli in the primary screen were considered positive if they produced IL-2 ≥ 500 pg/mL above background (values were background-subtracted using the PBS-only control; negative values were set to 0). Although most wells showed minimal signal (near zero), occasional wells—particularly those near the positive-control wells—produced spurious signal likely reflecting technical spillover rather than true positives. This signal was usually low-level (up to a few hundred pg/mL) but occasionally exceeded 500 pg/mL. The ≥ 500 pg/mL threshold was set to exclude most such artifacts while retaining sensitivity to detect genuine positives. All primary screen positives and a small number of borderline values were subsequently re-tested in a secondary hit validation assay, verifying true positives and excluding false positives (see "Secondary hit validation").

*Secondary hit validation:* Positive hits from the primary screen were re-tested in an independent secondary hit validation assay, using fresh aliquots of TCR-expressing cells and stimuli and run on a different day. Anti-CD3ε was included as a positive control, and stimuli were assayed in triplicate—the plate layout was designed to prevent spillover between wells (see Supplementary Fig. 8b for example layouts). Stimuli that gave IL-2 ≥ 250 pg/mL in ≥ 2 replicates were considered true positives, and those that did not were considered false positives. Because stimuli were spatially isolated from positive-control wells, spillover artifacts observed in the primary screen that exceeded the 500 pg/mL threshold did not recur, allowing genuine reactivity to be confirmed. The confirmatory threshold was chosen because a true positive near the 500 pg/mL primary cutoff would be expected to exceed 250 pg/mL in >95% of individual replicate measurements (assuming a SD of ~150 pg/mL, as observed for values in this range); this cutoff therefore minimizes exclusion of genuine low positives while requiring reproducibility to confirm their reactivity. Values reported in the screen reflect the confirmed result for each stimulus: primary screen values were retained for true positives, and values for stimuli determined to be false positives were replaced with their secondary validation measurement (mean of triplicates).

**TCR cloning into retroviral expression vectors**

The following sections describe 96-well cloning of TCRs into retroviral expression vectors, which was used for cloning most TCRs into plasmids; in some cases, similar methods were used on a smaller number of TCRs but with scaled-down methods (e.g. single tubes, PCR strip tubes, or partial plates).

*Preparing Golden Gate components:* Our Golden Gate assembly strategy consists of four components (see Supplementary Fig. 6a and Supplementary Table 1): the synthesized *TRA* and *TRB* for a given TCR each flanked by cut sites for the Type IIS restriction enzyme SapI, and two universal plasmids—a Destination Vector and a Middle Insert Vector, each containing two SapI cut sites that allow seamless integration of the complete TCR into a single expression vector. SapI was selected because its 7-base recognition sequence is expected to occur less frequently in TCR sequences than other commonly used Type IIS restriction enzymes with a 6-base recognition sequence.

Paired *TRA* and *TRB* variable regions were identified by scTCRseq. These consisted of the four framework (FR) and three complementarity-determining (CDR) regions (FR1-CDR1-FR2-CDR2-FR3-CDR3-FR4). 5' and 3' attachments containing SapI cut sites were added to each variable region—attachment sequences are listed in Supplementary Table 1. Variable regions were screened for endogenous SapI cut sites and, where present, silent mutations were manually introduced to disrupt the cut site. The final Golden Gate-ready sequences were synthesized as two plates of IDT eBlocks in 96-well format resuspended in nuclease-free water at 10 ng/µL—one plate contained the *TRA* sequences and the other contained the *TRB* sequences with a given well corresponding to a given TCR pair. Plates were stored at -20°C until use. All Golden Gate sequences used in this study are listed in Supplementary Table 1.

The Destination Vector and Middle Insert Vector sequences are listed in Supplementary Table 1 and will be available from Addgene upon publication. The Destination Vector contains the vector backbone and comprises universal TCR elements (*TRA* leader and *TRB* constant region), a puromycin resistance cassette, murine CD4, retroviral elements (5' and 3' long terminal repeats [LTRs]), a *ccdB* cassette to select against colonies transformed with undigested Destination Vector, and ampicillin (amp) and chloramphenicol (chlor) resistance cassettes. This vector was derived from a previously described TCR expression vector (*3*) by serial restriction digestion and insertion of synthesized DNA fragments (Twist) with complementary restriction overhangs to 1) remove two SapI cut sites, one in CD4 (by silent mutation) and another in the vector backbone, and 2) insert the *TRA* leader/*ccdB*/chlor-R/*TRBC* fragments. The resulting Destination Vector was propagated in One Shot *ccdB* Survival 2 $T1^R$ Competent Cells (Thermo A10460) in Luria Broth (LB) with 100 µg/mL ampicillin (ApexBio K2801) and 25 µg/mL chloramphenicol (Zymo A1002-5). The Middle Insert Vector contains universal TCR elements that are inserted between the *TRA* and *TRB* variable regions in the final product (*TRA* constant region [*TRAC*], a self-cleaving P2A linker, and the *TRB* leader) as well as a kanamycin resistance cassette. This vector was generated by digesting the pUK21 backbone (Addgene #49788) with PciI and BamHI followed by ligation of a digested synthesized fragment (Twist) containing the *TRAC*-P2A-*TRB* leader insert and complementary restriction overhangs. The resulting Middle Insert Vector was propagated in One Shot TOP10 Competent Cells (Thermo C404003) in LB with 50 µg/mL kanamycin (Zymo A1003-5). For each vector, a single colony verified by ZeroPrep whole-plasmid sequencing (Plasmidsaurus) was expanded, purified as a single large batch by pooling midipreps (Qiagen Plasmid Plus 12945), quantified by NanoDrop, re-verified by whole-plasmid sequencing of the final purified prep (Plasmidsaurus), then stored as aliquots at -20°C.

Upon thermal cycling in the presence of SapI and T4 ligase, and transformation into standard competent *E. coli* cells followed by amp selection, the final product containing the complete TCR is highly enriched because unrecombined Destination Vector is negatively selected due to the presence of *ccdB* and unrecombined Middle Insert Vector is negatively selected due to the lack of an amp-R cassette.

*Golden Gate assembly reaction*: Two 96-well plates containing *TRA* and *TRB* Golden Gate-ready sequences (see above) were combined 1:1 and mixed using a 12-channel pipette to give an equimolar *TRA:TRB* mixture in a single plate. Each Golden Gate reaction was performed in 10 µL total volume and contained each of the four components at 3 nM (for the Middle Insert Vector, this refers to the insert region, not the entire plasmid), 7.5 U SapI (NEB R0569L), 250 U T4 DNA Ligase (NEB M0202M), 1 µL 10X T4 DNA Ligase buffer (NEB B0202), and nuclease-free water to 10 µL (NEB B1500L). In practice, 1.5 µL/well of the 1:1 *TRA*:*TRB* eBlock mixture was added to a 96-well PCR plate using a 12-channel pipette (Bio-Rad HSL9641), then 8.5 µL/well of a master mix containing the remaining components was added on top (192 ng/well Destination Vector, 63 ng/well Middle Insert Vector, 0.75 µL/well SapI [10 U/µL], 0.125 µL/well T4 DNA ligase [2000 U/µL], 1 µL/well 10X T4 DNA Ligase buffer, and nuclease-free water to 10 µL/well total volume). 96-well plates were sealed, mixed, and briefly centrifuged. The Golden Gate assembly reaction was performed in a thermal cycler with the following conditions: 60 cycles of 37°C for 5 min and 16°C for 5 min, followed by 60°C for 5 min, then a hold at 12°C once completed.

*Transformation:* A 96-well MultiShot FlexPlate of TOP10 Competent Cells (Thermo C4081201) was removed from the freezer and placed in a chilled 96-well metal block on ice for ~30 s. The foil seal was removed and 1 µL/well of the Golden Gate assembly mixture (above) was added. The plate was covered with a Qiagen AirPore seal (19571) and incubated on ice for 20 min. The plate was heat-shocked in a 42°C water bath for 30 s, then transferred back to the chilled metal block for ~1 min. The seal was removed, 90 µL of SOC medium was added to each well, and the plate was incubated at 37°C (static) for 1 h. During this time, 8 square 6x6 gridded plates containing LB 100 µg/mL carbenicillin (Simport Scientific D210-16) were pre-warmed at 37°C for ~30 min then allowed to dry with the lid removed in a laminar flow hood for ~30 min. The bacterial suspension was diluted 1:20 and 1:200 in LB, and 10 µL of the undiluted stock and each dilution were spot plated onto the 6x6 grid using an Integra adjustable spacing electronic multi-channel pipette to switch between 96-well and 6-point grid spacing. Six adjacent wells from two adjacent rows in the 96-well plate were plated in this fashion (e.g. agar plate 1 contained wells A01-A06 and B01-B06, three dilutions each), allowing the entire 96 wells to be plated on the 8 agar plates. Plates were incubated overnight at 37°C.

*Miniprep and sequencing:* Three single colonies per TCR were inoculated—one colony per plate, using sterile inoculating picks (LevGo 18250-SP)—into the corresponding well of three 96-well S-block plates (Qiagen 19585), with each well containing 1.3 mL LB 100 µg/mL amp. Plates were covered with a Qiagen AirPore seal and incubated overnight in a 37°C shaking incubator. The next day minipreps were purified using the QIAprep 96 Plus Miniprep Kit (Qiagen 27291) and QIAvac 96 (19504) according to the manufacturer's instructions, except the final elution step was performed by centrifuging 5 min at maximum speed instead of by vacuum. A 6 µL aliquot of each miniprep was added to a 96-well semi-skirted plate (Bio-Rad HSL9641), dehydrated by heating at 65°C for ~45 min, sealed, and submitted for whole-plasmid sequencing (Plasmidsaurus); the remaining plasmid prep was stored at -20°C. Sequences were analyzed by aligning each plasmid to the expected *TRA*, *TRB*, and vector sequences using Geneious Prime, and the cloning outcome (perfect assembly, assembly error, or DNA synthesis error) was manually tallied (see Supplementary Fig. 6b-d); if multiple correct clones were obtained the first occurrence was typically selected. Apparent length variation at a poly-G homopolymer between the puromycin cassette and IRES-CD4 regions was treated as a sequencing artifact rather than true variation, as homopolymer miscalls are a known limitation of Nanopore sequencing (*4*).

*Plasmid concentration measurement and normalization:* Correctly assembled plasmids were transferred to the corresponding well of a new 96-well microtube rack (Qiagen 19588). The concentration of each plasmid was measured in duplicate in 96-well format using the Quant-iT dsDNA Assay Kit, broad range (Thermo Q33130 and M33089) according to the manufacturer's instructions. Fluorescence was measured using an Agilent BioTek Synergy H1 plate reader with an excitation of 485 nm and emission of 528 nm. Standards were fit to a linear standard curve in Microsoft Excel and used to interpolate the concentrations of each plasmid (from the mean of the two replicates). Plasmids were normalized to 50 ng/µL by adding TE buffer (VWR 786-151), then frozen at -20°C; rare preps that were below this concentration were used undiluted.

**TCR expression in cell lines**

*Transfection into Plat-E retroviral packaging cells:* The Plat-E retroviral packaging cell line (*5*) (Cell Biolabs RV-101) was cultured in sterile-filtered DMEM 10% FCS 1% penicillin/streptomycin (pen/strep; Thermo 15140122) with 1 µg/mL puromycin (Thermo A1113803) and 10 µg/mL blasticidin (Thermo A1113903). The day prior to transfection, Plat-E cells were harvested using trypsin-EDTA (Sigma T4049-100ML), and centrifuged (5 min, 500*g*); the supernatant was aspirated and the pellet resuspended in DMEM 10% FCS 1% pen/strep (without puromycin or blasticidin). Cells were counted in duplicate using a Countess II cell counter, and concentration was adjusted to 2.75 x $10^5$ cells/mL. ~4.0 x $10^4$ cells/well (~150 µL/well) were seeded into a 96-well flat-bottom tissue culture plate using a 12-channel pipette and cultured overnight in a cell culture incubator (37°C with 5% $CO_2$). The next day, a 96-well plate of plasmids was thawed. Transfection was performed using the Lipofectamine 3000 transfection kit (Thermo L3000001). Two master mixes were prepared in Opti-MEM (Thermo 31985062): a P3000 master mix (0.2 µL/well P3000 reagent and 2.8 µL/well Opti-MEM) and a Lipofectamine master mix (0.3 µL/well Lipofectamine and 4.7 µL/well Opti-MEM). The P3000 master mix was aliquoted into a new 96-well PCR plate using a 12-channel pipette. Next, 2 µL of each plasmid (normalized to 50 ng/µL as above) was added to the corresponding well and mixed by pipetting. The Lipofectamine master mix was then added, mixed by pipetting, briefly centrifuged, sealed, and incubated ~15 min at room temperature. The 10 µL transfection mixture was added to the 96-well plate of Plat-E cells prepared the day prior, gently mixed, and incubated 48 h in a tissue culture incubator.

*Transduction of 58αβ$^{-/-}$ hybridoma cells:* The 58αβ$^{-/-}$ TCR-deficient T cell hybridoma cell line (*6*) was cultured in DMEM 10% FCS 1% pen/strep and inoculated several days ahead of the planned transduction. ~48 h after transfection of Plat-E retroviral packaging cells (see above), the 96-well plate containing the Plat-E cells was centrifuged (5 min, 1,000*g*) and 100 µL/well of the supernatant was transferred to a new 96-well U-bottom tissue culture plate. Protamine sulfate (1.5 µL/well of a 1 mg/mL solution, sterile-filtered in MilliQ water; MP Biomedicals 0219472905) was added to each well of retrovirus-containing supernatant using a 12-channel pipette and mixed by pipetting. $58\alpha\beta^{-/-}$ cells were counted in duplicate using a Countess II cell counter, adjusted to $5.0 \times 10^5$ cells/mL in DMEM 10% FCS 1% pen/strep, and $\sim 2.5 \times 10^4$ cells/well (~50 µL/well) were added to the retroviral supernatant/protamine sulfate mixture and mixed gently by pipetting. Cells were transduced by centrifugation at 1,000*g* for 120 min at 32°C. The cell pellet was resuspended by pipetting, and the plate was incubated overnight in a tissue culture incubator.

*Puromycin selection of transduced cells:* The day following retroviral transduction, eight 12-well tissue culture plates containing 2 mL/well DMEM 10% FCS 1% pen/strep plus 1 µg/mL puromycin were prepared. Transduced cells were mixed by pipetting, then transferred to the corresponding wells in the 12-well plate using an Integra adjustable spacing 4-channel electronic pipette, and gently mixed; plates were then incubated in a tissue culture incubator. Cell growth was monitored daily by microscopy, and untransduced controls were included in each experiment.

*Cryopreservation and flow cytometry analysis of surface TCR and CD4 expression:* After ~4 days of puromycin selection, transduced wells typically showed robust growth with orange-yellow medium, whereas untransduced wells showed shrunken cells with minimal growth and red medium. Next, 2 mL of each TCR-expressing hybridoma culture was transferred to the corresponding well of a sterile Qiagen 96-well S block plate using an Integra adjustable spacing 4-channel electronic pipette. A 50 µL aliquot of each well was transferred to a new 96-well U-bottom plate containing 50 µL/well 2X antibody master mix in flow buffer containing the following antibody-fluorophore conjugates: TCRβ-APC (H57-597; BioLegend 109212; 1:200), CD4-PE (GK1.5; BioLegend 100408; 1:200), and TruStain FcX PLUS (anti-mouse CD16/32 S17011E; BioLegend 156604; 1:400). An untransduced and unselected sample of $58\alpha\beta^{-/-}$ cells was stained and analyzed side-by-side. Cells were gently mixed, incubated on ice for 20 min, washed with excess flow buffer, centrifuged (5 min, 500*g*), resuspended in ~200 µL flow buffer, and analyzed on a BD CytoFLEX flow cytometer; surface CD4 and TCRβ expression for all TCRs is shown in Supplementary Fig. 6e-f. A subset of wells from the 96-well S block was counted and viability measured using a Countess II cell counter. The plate was sealed with a Qiagen AirPore seal and centrifuged (10 min, 500*g*). The plate was unsealed in the tissue culture hood, and supernatant was aspirated using an 8-channel vacuum adapter. Each well was resuspended in ~1 mL CELLBANKER cryopreservation medium (amsbio 11910), a subset of wells were again counted and viability measured using a Countess II cell counter (targeting $\sim 5 \times 10^5$-$5 \times 10^6$ cells/mL), and 3 x ~325 µL aliquots of each cell line were added to the corresponding well of 3 x 96-well racks of 0.5 mL Matrix tubes (Thermo 3745-BR) and then capped

(Thermo 4477RED) using an 8-channel electronic capper/decapper (Thermo 4105MAT). The racks were placed in an insulated foam box at -80°C overnight, then transferred to freezer racks for long-term storage.

*Flow cytometry analysis of control TCRs with Vα and Vβ-specific antibodies:* Staining of control TCRs (OT-II, SMARTA, HH5-1, or P25) with Vα and Vβ-specific antibodies (each stained with the panel appropriate for its TCR; see Supplementary Fig. 6g-j) was performed as described above, except that the cells (or untransduced and unselected 58αβ$^{-/-}$ cells) were stained with the appropriate subset of the following antibodies and analyzed on a BD LSRII flow cytometer: CD3ε-APC (145-2C11; BioLegend 100312; 1:100), Vα2-PE (B20.1; BioLegend 127807; 1:100), Vβ5.1/5.2-FITC (MR9-4; BioLegend 139513; 1:100), CD4-BV421 (GK1.5; BioLegend 100443; 1:200), Vβ8.3-FITC (1B3.3; BioLegend 156305; 1:100), Vβ8.1/8.2-PE (MR5-2; BioLegend 140103; 1:100), and Vβ11-PE (KT11; BioLegend 125907; 1:100).

**DC expansion and isolation**

A B16 melanoma cell line overexpressing Flt3L (*7*) to induce dendritic cell (DC) hyperplasia was grown in sterile-filtered DMEM 10% FCS 1% pen/strep. Adult SPF C57BL/6 mice were injected with ~5 x $10^6$ cells intraperitoneally. 10-14 days later, spleens were harvested and placed in sterile PBS. In a tissue culture hood, 1-2 spleens were chopped into ~2-4 pieces, placed into a gentleMACS C tube (Miltenyi 130-093-237), and processed using reagents from the StraightFrom Spleen CD11c MicroBead Kit, mouse (Miltenyi 130-130-146), prepared per the manufacturer's instructions. For each spleen, 820 µL buffer S, 20 µL Enzyme A, 60 µL Enzyme D, and 100 µL anti-CD11c beads were added. The C-tube was closed tightly, attached to a Miltenyi gentleMACS Octo Dissociator with heater, and the program 37C_m_SDK_1 was run. Following a brief centrifugation, the cell suspension was filtered through a sterile 70 µm cell strainer (Corning) and remaining tissue was mashed with a plunger from a sterile 1 mL syringe. The strainer was washed with 1-2 mL sterile-filtered MACS separation buffer (PBS 0.5% BSA 2 mM EDTA—prepared by diluting MACS BSA Stock Solution [130-091-376] 1:20 with autoMACS Rinsing Solution [130-091-222] per the manufacturer's instructions). Cells were centrifuged (5 min, 500*g*) and the supernatant was aspirated. The pellet was resuspended in ~2 mL MACS separation buffer and filtered through a new 70 µm cell strainer. One sterile LS column (Miltenyi 130-042-401) per C-tube was placed in a Miltenyi QuadroMACS separator stand and prepared by applying 3 mL MACS separation buffer. The cell suspension was applied to the column and flow-through containing unlabeled cells was collected. The column was washed three times with 3 mL MACS separation buffer—flow-through was collected and combined with the unlabeled cells. The column was removed from the magnetic stand and positioned over a new sterile 15 mL conical tube. CD11c$^+$ cells were eluted by applying 5 mL MACS separation buffer and immediately firmly pushing the plunger into the column. If multiple columns were processed in parallel, the positive fractions were combined and centrifuged (5 min, 500*g*). Supernatant was aspirated and the cell pellet was resuspended in sterile-filtered DMEM 10% FCS 1% pen/strep. Cells were counted in duplicate using a Countess II cell counter and then normalized to 1 x $10^7$ cells per mL in DMEM 10% FCS 1% pen/strep. Analysis of pre- and post-enrichment fractions by flow cytometry using antibodies against CD11c (N418-PE; BioLegend 117308; 1:100) and MHCII (M5/114.15.2-APC; BioLegend 107614; 1:100) demonstrated that the enriched $CD11c^+$ fraction was ~96-97% $CD11c^+$ $MHCII^+$ cells. Typically, about 50-75 x $10^6$ DCs were obtained per B16-Flt3L-injected mouse spleen.

**Growth of hCom2v bacterial strains and preparation for TCR reactivity screening**

Individual bacterial strains in hCom2v were revived from frozen glycerol stocks and grown individually in liquid and/or solid media in an anaerobic chamber (10% $CO_2$, 5% $H_2$, 85% $N_2$). A complete list of strains and media used is provided in Supplementary Table 3. Solid media were purchased: YCFAC (Anaerobe Systems AS-675), ETSA (Anaerobe Systems AS-548), RCA (Anaerobe Systems AS-6061), and Columbia (Hardy Diagnostics A16BX). Some liquid media were also purchased: chopped meat (Hardy Diagnostics AG20H or BD 297307), mGAM (HiMedia M1801-500G), and Anaerobe Systems YCFAC broth (AS-680); however, most liquid media were made in-house from individual components (see Supplementary Table 3). For colonization of mice with hCom2v, strains were grown individually then pooled and stored as a glycerol stock as described previously (*8*). For preparation of individual strains for T cell reactivity screening, strains were grown individually, pelleted by centrifugation (10 min, 4,000*g*), resuspended in sterile PBS, and adjusted to an $OD_{600}$ of ~1.0. Suspensions were then autoclaved, aliquoted, and frozen at -80°C in 96-well cluster tubes (Corning 4411 and 4418). Strain preparations underwent QC by culture on agar plates, MALDI-TOF, and 16S rRNA gene sequencing. Strains that either could not be grown to a sufficient extent for large-scale screening or that contained < 1% of the correct strain on 16S sequencing analysis were denoted as missing values (NA) in TCR screening analyses. A small number of strain preparations contained the intended strain plus a contaminant; in these cases, negative results were retained while positive results were retained only if confirmed with a pure strain preparation or if the TCR's cognate antigen was identified and shown to be expressed by the intended strain. Because strains were optimized for growth and purity over the course of the study, some strains became available for screening after the analysis of a subset of TCRs; thus, results for these strains are present for some TCRs and NA for others. For the primary screen against 128 stimuli, strain preparations were aliquoted into 384-well master mix plates using an Integra 8-channel adjustable spacing electronic pipette, sealed, and frozen at -80°C.

**Preparation of hCom2v cecal and colonic contents for TCR reactivity screening**

Here, 3-4 hCom2v-colonized mice were processed in parallel. The cecum and colon were dissected from each mouse, contents were transferred to a sterile tube, 1 mL sterile PBS was added, and the sample was homogenized by taping horizontally on a vortex and vortexing vigorously for 5 min. Samples were centrifuged (5 min, 400*g*) to pellet large debris and the supernatant was transferred to a new tube through a 70 μm cell strainer; samples from multiple mice were pooled at this stage. The suspension was centrifuged (5 min, 8,000*g*) and the supernatant was aspirated. The pellet was resuspended in sterile PBS and normalized to an $OD_{600}$ of ~1.0. An aliquot was frozen and community composition verified by metagenomic sequencing. The remaining sample was autoclaved, aliquoted, and stored at -80°C. For co-culture assays, 4 μL/well of this prep was used.

**Preparation of dietary components for TCR reactivity screening**

Chow pellets or individual proteinaceous components of the 5K67 (LabDiet) mouse chow (wheat, corn, oat, fish meal, soybean, alfalfa, and yeast; commercially sourced) were resuspended at 60 mg/mL in sterile PBS. Four stainless steel beads (Qiagen 69989) were added per preparation, and the sample was homogenized by vortexing (taped horizontally on a vortex for 5 min). Samples were heated at 70°C for ~10 min, then aliquoted and frozen at -80°C. For co-culture assays, 4 μL/well of each prep was used.

**Co-culture assay and IL-2 measurements**

The co-culture assay consists of a TCR-expressing hybridoma line generated as above (see "TCR expression in cell lines"), primary DCs (see "DC expansion and isolation"), and a test stimulus (typically prepared in advance and frozen). As a starting point for all assays, cryopreserved TCR-expressing lines were inoculated into culture in DMEM 10% FCS 1% pen/strep 1 μg/mL puromycin 3-7 days in advance; we found that inoculating 7 days in advance and then counting cells and passaging after 3 days led to the most consistent cell numbers for large-scale screening. We observed that cryopreservation substantially reduced DC stimulatory capacity (see Supplementary Fig. 7c), so DCs were freshly prepared for all assays and mice were injected with B16-Flt3L tumors 10-14 days before a planned co-culture assay.

*Optimization:* We optimized the co-culture assay in 384-well format using hybridomas expressing the OT-II TCR or a non-reactive negative control TCR from GF mice (Supplementary Fig. 7). By co-culturing $2 \times 10^4$ TCR-expressing cells with increasing numbers of DCs against titrated whole OVA (Hooke DS-0142) in a 40 μL culture volume, we obtained peak IL-2 AlphaLISA signal at $1 \times 10^5$ DCs/well; signal did not differ substantially when cells were co-cultured for 24 vs 48 h and was abrogated by an anti-MHCII blocking antibody (Y-3P; Bio X Cell BE0178; 10 μg/mL final concentration; see Supplementary Fig. 7a). Both TCRs responded to a variety of positive control stimuli including PMA/ionomycin, soluble anti-CD3ε, and anti-CD3/CD28 Dynabeads (Thermo 11452D), with minimal signal to medium alone (Supplementary Fig. 7b). Titrating TCR-expressing cell numbers in the presence of 1 x $10^5$ DCs/well, we found that 2 x $10^4$ TCR-expressing cells gave optimal signal (Supplementary Fig. 7d-e). We therefore concluded that 2 x $10^4$ TCR-expressing cells and 1 x $10^5$ DCs per well cultured in 40 µL for 24 h gave optimal signal, with minimal background from non-reactive TCRs or under MHCII blockade.

*Primary screen:* TCR-expressing hybridoma lines were grown in DMEM 10% FCS 1% pen/strep 1 µg/mL puromycin in 6-well tissue culture plates. Typically, 12-14 TCRs were assayed in parallel in each experiment. Cells were transferred to a sterile 15 mL conical tube and centrifuged (5 min, 500*g*); the supernatant was aspirated, and the cell pellet was resuspended in 2 mL DMEM 10% FCS 1% pen/strep (without puromycin). Cells were counted in duplicate using a Countess II cell counter and the concentration was adjusted to 2 x $10^6$ cells/mL in DMEM 10% FCS 1% pen/strep. Primary DCs were isolated as described under "DC expansion and isolation"; after elution from the magnetic column, the cell pellet was resuspended in 10 mL DMEM 10% FCS 1% pen/strep. Cells were counted in duplicate using a Countess II cell counter and the concentration was adjusted to 1 x $10^7$ cells/mL. A pre-aliquoted 384-well master mix plate (Corning 3342) containing defined volumes of the 128 stimuli was thawed, and PMA/ionomycin and anti-CD3ε were added fresh to the appropriate wells. Stimuli in the plate were diluted with DMEM 10% FCS 1% pen/strep, mixed by pipetting, and then dispensed into the appropriate wells of a 384-well tissue culture plate (Thermo 164688; see Supplementary Fig. 8a for plate layout) using the repeat dispense mode on an Integra 16-channel electronic pipette. Dispensed volumes ranged from 4-7.5 µL per well depending on the dilution, in all cases delivering the equivalent of 4 µL of undiluted stimulus. Using an Integra 16-channel electronic pipette, medium (added to bring the final well volume to 40 µL), 1 x $10^5$ DCs (10 µL), and 2 x $10^4$ TCR-expressing cells (10 µL) were aspirated, separated by air gaps; the combined volume was dispensed into the stimulus-containing wells and mixed by pipetting. The plate was incubated in a tissue culture incubator for ~24 h. The next day, the plate was centrifuged (5 min, 1,000*g*), and ~35 µL supernatant/well was transferred to a new 384-well plate, sealed, and frozen at -20°C until analysis by AlphaLISA as described below (typically within 1-2 weeks).

*Secondary hit validation:* Here, 4 µL of undiluted stimulus was added to individual wells of a 384-well tissue culture plate (see example plate layout in Supplementary Fig. 8b). Medium, 1 x $10^5$ DCs, and 2 x $10^4$ TCR-expressing cells (grown from a fresh aliquot) were added and co-cultured as described above under 'Primary screen', and IL-2 was measured by AlphaLISA.

Criteria for interpreting primary screen and secondary hit validation results are described above under "TCR screen design." Heatmaps were generated using the ComplexHeatmap package (*9*) in R (v4.4.1).

*Peptide epitope mapping:* Here, 4 µL of a 0.1 mg/mL solution of peptide in PBS was added to individual wells of a 384-well tissue culture plate (giving a 10 µg/mL final concentration in 40 µL).

Medium, 1 x $10^5$ DCs, and 2 x $10^4$ TCR-expressing cells were added and co-cultured as described above under 'Primary screen', and IL-2 was measured by AlphaLISA.

A subset of assays for zein and gliadin epitope identification was conducted in 96-well format with IL-2 measured by ELISA (Supplementary Fig. 11c-d). Here, TCR-expressing cell lines found to be diet-reactive were screened against a panel of mouse chow ingredients or peptide epitopes. In the initial screening, 2 x $10^4$ TCR-expressing hybridoma cells and 5 x $10^4$ dendritic cells were used per assay. Lines giving weak or inconclusive signal were re-tested with 1 x $10^5$ hybridoma cells and 5 x $10^5$ dendritic cells to increase signal. After 24 h of co-culture, IL-2 in the culture supernatant was measured by ELISA (BioLegend, 431001) as described (*10*). Standards were fit to a four-parameter logistic curve and sample IL-2 values interpolated using GraphPad Prism. Samples that could not be interpolated were flagged as below ('< Curve') or above ('> Curve') the standard curve; these were assigned values of 0 and 5,775 pg/mL, respectively, corresponding to the lowest (blank) and highest standards included in the curve.

The following peptides were used in this study (all synthesized by GenScript at ≥95% purity): $SBP_{76\text{-}89}$ YYVGFDANQGAELQ (*3*), $SBP_{405\text{-}419}$ YDAFAINMVKTDNAA (*3*), $OVA_{323\text{-}339}$ ISQAVHAAHAEINEAGR (*11*), $TPRL_{29\text{-}53}$ SDYFTVTPQVLEAVGGKVPATINGK (*3*), α-$Zein_{223\text{-}233}$ FYQQPIIGGAL (*10*), α-Zein FYLHAMPNA, $Gliadin_{273\text{-}285}$ CNVYIPPYCTIAP (*12*), Am3735 TLYIGSGAILSGN (*13*), Am3740 LIFESSNALGLGR (*13*), Am2C.1 ISAASYVVPPLPNKR (*14*), Am2C.9 NKRSEANPANPPSFA (*14*), Am6H.3 HPSRLHAPAITQNAL (*14*), Am8H.1 LPGYGLPLCAGPGAS (*14*), $BT4295_{541\text{-}554}$ EEFNLPTTNGGHAT (*15*), $bHex_{568\text{-}581}$ KWDYKGSRVWLNDQ (*16*), LCMV $GP_{61\text{-}80}$ P13 GLNGPDIYKGVYQFKSVEFD (*17*), HH_1713 E1 GNAYISVLAHYGKNG (*18*), and MTB $Ag85b_{240\text{-}254}$ FQDAYNAAGGHNAVF (*19*).

*IL-2 AlphaLISA:* IL-2 was measured in the culture supernatant using the AlphaLISA Mouse IL-2 Detection Kit (Revvity AL585C or AL585F). Frozen supernatant from co-culture assays above was brought to room temperature, mixed in a plate mixer, and briefly centrifuged. A standard curve of mouse IL-2 diluted in DMEM 10% FCS 1% pen/strep was prepared fresh according to the manufacturer's instructions and was run on every plate. Using an Integra 16-channel electronic pipette, 8 µL of a 2.5X master mix of anti-mIL2 acceptor beads and biotinylated anti-mIL2 antibody (in 1X AlphaLISA immunoassay buffer) was aspirated, followed by an air gap, then 2 µL of each sample or standard. The combined volume was dispensed into the corresponding well of a 384-well white OptiPlate (Revvity 6007290), mixed by pipetting, and then sealed (Revvity 6050185) and incubated for 60 min at room temperature. A 2X streptavidin donor bead master mix (in 1X AlphaLISA immunoassay buffer) was prepared; 10 µL was added to each well and mixed by pipetting, and the plate was resealed and incubated for 30 min at room temperature. The streptavidin donor bead addition step was performed in the dark and plates were subsequently kept protected from light. Plates were centrifuged briefly and then data were collected on a BMG CLARIOstar Plus instrument with the following filters and settings:

excitation 680-40; dichroic filter SP AS; emission 620-10; 384-well aperture. Standards were fit to a four-parameter logistic curve and sample IL-2 values interpolated using MyAssays. Samples that MyAssays could not interpolate were flagged as below ('< Curve') or above ('> Curve') the standard curve; these were assigned values of 0 and 100,000 pg/mL, respectively, corresponding to the lowest (blank) and highest standards included in the curve.

**SBP and α-Zein tetramer production**

The generation of peptide-MHCII tetramers has been described in detail (*20*). In brief, soluble I-$A^b$ molecules heterodimerized through Fos-Jun leucine zipper (LZ) motifs and covalently linked to N-terminal SBP epitope $SBP_{74-85}$ (DTYYVGFDANQG), C-terminal SBP epitope $SBP_{403-414}$ (SQYDAFAINMVK), or $Zein_{223-232}$ (FYQQPIIGGA) peptide epitopes were expressed and biotinylated in stably transfected *Drosophila* S2 cells. Following immunoaffinity purification with anti-I-A antibody (clone Y-3P), these biotinylated peptide-MHCII complexes were titrated and tetramerized to streptavidin pre-conjugated to R-phycoerythrin (PE) or allophycocyanin (APC) fluorochromes (Prozyme). $Zein_{222-233}$ (VFYQQPIIGGAL) tetramers were constructed with a human IgG1Fc knob-in-hole (KIH) heterodimer backbone (*21*) instead of leucine zipper, and affinity purified using protein G. Although the LZ and KIH Zein tetramers gave similar results, the KIH version showed superior staining and was used in most experiments; some data points in Fig. 3h and one of two experiments shown in Supplementary Fig. 12c-f used the LZ tetramer.

**Intestinal lamina propria (LP) lymphocyte isolation**

The small intestine (SI) or large intestine (LI; cecum plus colon) was dissected from each mouse, and fat, Peyer's patches (PPs), and luminal contents were removed. Intestinal tissues were opened longitudinally, washed by swirling in HBSS 2% FCS for 5-10 s, cut into ~0.5 cm pieces, and placed into a 50 mL tube with 10 mL pre-warmed RPMI 1% FCS 2 mM EDTA (Boston BioProducts). Each SI was split into two halves that were processed in parallel and combined at the Percoll step below. Tissue pieces were incubated at 37°C with shaking for 15 min, collected on a 100 μm cell strainer (Corning), and placed in 10 mL fresh RPMI 1% FCS 2 mM EDTA for another 15 min with shaking at 37°C. Tissue pieces were collected again on a fresh 100 μm cell strainer and flow-through (containing predominantly epithelial cells and intraepithelial lymphocytes) was discarded. Tissue pieces were transferred to a new 50 mL tube containing 10 mL RPMI 20% FCS with 1 mg/mL collagenase A (Roche), 0.1 mg/mL DNase I (Roche), and 0.1 U/mL Dispase II (Sigma) and incubated at 37°C with shaking for 60 min. This suspension was filtered through a fresh 100 μm cell strainer (Miltenyi). Any remaining tissue was mashed with a 1 mL syringe plunger, and the cell strainer was washed with ~30 mL cold RPMI. The resulting suspension was vortexed, centrifuged (5 min at 550*g* with low brake), and the supernatant was aspirated. Lymphocytes were enriched by centrifugation in 10 mL 35% Percoll (Sigma) (v/v in PBS; 20 min at 3000 rpm)—the two SI halves for a given mouse were combined at this step. Percoll was removed by aspiration and the cell pellet containing lamina propria lymphocytes was processed for downstream applications.

**Cell isolation from secondary lymphoid organs**

Spleen, PPs, or lymph nodes (LNs) were dissected from each mouse and fat was removed. Tissues were placed in flow buffer on ice, mashed through a 70 μm cell strainer (Corning) with a 1 mL syringe plunger, and the strainer was washed with 1-2 mL of flow buffer. The resulting cell suspension was centrifuged (5 min at 500*g*), supernatant was aspirated, and the cell pellet was processed for downstream applications.

**Peptide-MHCII tetramer staining and extra/intracellular staining for flow cytometry**

Cell suspensions from mouse tissues were obtained as described above. Sterile-filtered HBSS 2% FCS was used as flow buffer. Staining was performed as follows:

*Live/dead staining:* Cells were resuspended in PBS (protein-free) with 1 μL/mL fixable near-IR live/dead stain (Thermo L10119) for 15 min on ice, washed with excess flow buffer, incubated 2-3 min on ice, centrifuged (5 min, 550*g*, low brake), and supernatant was removed by aspiration.

*Cell counting:* 3 μL per tissue from a suspension (resuspended in a defined volume) was added to 282 μL flow buffer and 15 μL counting beads (Spherotech ACBP-100-10). 2,000 bead-gated events were recorded on the flow cytometer and used to calculate the absolute cell number based on the known bead concentration and volume of the cell suspension.

*Peptide-MHCII tetramer staining:* Peptide-MHCII tetramers containing SBP N- and C-terminal epitopes and the C-terminal α-Zein epitope fluorescently conjugated to PE or APC were produced in-house (see "SBP and α-Zein tetramer production" for details). For most intestinal LP tetramer staining experiments, the cell suspension from one mouse's SI or LI was resuspended in ~45 μL per mouse per tissue pre-warmed RPMI 10% FCS 50 nM dasatinib (Sigma SML2589-50MG) plus TruStain FcX PLUS (anti-mouse CD16/32 S17011E; BioLegend 156604; 1:400), incubated for 5 min, and then split into two 25 μL aliquots per tissue. 25 μL of pre-mixed 2X tetramer (both SBP tetramers or the α-Zein tetramer in both PE and APC) in pre-warmed RPMI 10% FCS 50 nM dasatinib was added to give 25 nM final concentration of each tetramer. Cells were stained in a 37°C water bath for 1 h, gently mixing by tapping the side of the tube every 20 min.

*Extracellular staining:* The tetramer-cell suspension above was moved to ice and 2X extracellular master mix in RPMI 10% FCS 50 nM dasatinib was added on top and incubated 20 min on ice. Cells were washed with excess flow buffer, centrifuged (5 min, 550*g*, low brake), and supernatant was aspirated. The extracellular master mix typically contained the following antibody-fluorophore conjugates (clone, vendor, and final dilution are provided for each): B220-APC/Cy7 (RA3-6B2; BioLegend 103224; 1:200), CD11b-APC/Cy7 (M1/70; BioLegend 101226; 1:200), CD11c-APC/Cy7 (N418; BioLegend 117324; 1:200), F4/80-APC/Cy7 (BM8; BioLegend 123118; 1:200), NK1.1-APC/Cy7 (PK136; BioLegend 108724; 1:200), TER-119-APC/Cy7 (TER-119; BioLegend 116223; 1:200), CD45.2-FITC (104; BioLegend 109806; 1:200), and CD44-BUV395 (IM7; BD 568507; 1:800).

*Intracellular staining:* Foxp3 Fixation/Permeabilization working solution and 1X Permeabilization Buffer were prepared using the eBioscience Foxp3/Transcription Factor Staining Buffer Set according to the manufacturer's instructions (Thermo 00-5523-00). All intracellular staining steps were performed in 1.5 mL Eppendorf tubes. Cell pellets were resuspended in 200 µL of Foxp3 Fixation/Permeabilization working solution, mixed by pipetting up and down, and incubated ~30 min at room temperature protected from light. Cells were washed with 1 mL 1X Permeabilization Buffer, and centrifuged multiple times in a protocol intended to minimize cell loss due to the loose pelleting tendency of gut samples (6 min at 600*g* with low brake, then 1 min 30 s at 400*g* with low brake; tubes were removed from the centrifuge, gently rotated side-to-side to slide cell pellets on the side of the tube to the bottom, incubated vertically for 1 min, and then centrifuged 1 min at 300*g* with low brake); the supernatant was removed by aspiration. The pellet was resuspended in 200 µL 1X intracellular staining master mix and stained overnight at 4°C protected from light. The next morning, the suspension was washed with 1 mL 1X Permeabilization Buffer, centrifuged via the same multi-step protocol described above, the supernatant was aspirated, and cells were resuspended in ~500 µL HBSS 2% FCS prior to analysis on a flow cytometer. The intracellular master mix typically contained the following antibody-fluorophore conjugates in 1X Permeabilization Buffer 10% Brilliant Stain Buffer Plus (BD 566385) (clone, vendor, and final dilution are provided for each): CD4-BV785 (GK1.5; BioLegend 100453; 1:400), T-bet-PE/Cy7 (eBio4B10; Thermo 25-5825-82; 1:400), RORγt-RB705 (Q31-378; BD 570259; 1:800), TCRβ-BV605 (H57-597; BioLegend 109241; 1:400), Foxp3-BV421 (FJK-16s; Thermo 404-5773-82; 1:100), and TruStain FcX PLUS (anti-mouse CD16/32 S17011E; BioLegend 156604; 1:400). Some experiments also included Ki67-BUV805 (B56; BD 569635; 1:100).

*Analysis:* Samples were analyzed on 5-laser BD Symphony, LSRII, and Fortessa flow cytometers at the Stanford Shared FACS facility. Compensation was calculated using splenocytes stained with anti-CD4 (GK1.5; 1:200 separately in each color for 20 min on ice), fixed using the same procedure described above, washed with 1 mL 1X Permeabilization Buffer, resuspended and incubated overnight in 1 mL 1X Permeabilization Buffer at 4°C, and then centrifuged and resuspended in ~500 µL HBSS 2% FCS prior to analysis similar to above. Samples were filtered through a 35 µm cell strainer (Corning) just prior to flow cytometry analysis. For tetramer staining analyses, the entire sample was typically recorded.

**SBP tetramer enrichment from pooled hCom2v or SPF mice**

Cell suspensions from mouse LI LP were obtained as described under "Intestinal lamina propria (LP) lymphocyte isolation"; three mice were pooled per group (hCom2v, SPF-Tac, or SPF-Jax) following the Percoll step. Sterile-filtered HBSS 2% FCS was used as flow buffer. Staining was performed as follows—see "Peptide-MHCII tetramer staining and extra/intracellular staining for flow cytometry" for detailed reagent information, which was the same as described below except as noted:

*Peptide-MHCII tetramer staining:* The pooled LI LP cell pellet was resuspended in ~135 µL per group per tissue pre-warmed RPMI 10% FCS 50 nM dasatinib plus TruStain FcX PLUS, and incubated for 5 min. 150 µL of pre-mixed 2X SBP tetramers in pre-warmed RPMI 10% FCS 50 nM dasatinib was added to give 25 nM final concentration of each tetramer. Cells were stained in a 37°C water bath for 1 h, gently mixing by tapping the side of the tube every 20 min. Cells were washed with excess flow buffer, centrifuged (5 min, 550*g*, low brake), and the supernatant was aspirated.

*MACS bead staining and MACS enrichment:* Cell pellets were resuspended in 200 µL MACS bead master mix (100 µL RPMI 10% FCS + 50 µL anti-APC MACS beads [Miltenyi 130-090-855] + 50 µL anti-PE MACS beads [Miltenyi 130-048-801]) and incubated for 20 min on ice. Cells were washed with excess flow buffer, centrifuged (5 min, 550*g*, low brake), and supernatant was aspirated. While centrifuging, a Miltenyi LS column (130-042-401) was prepared by attaching it to a magnetic stand and adding 3 mL MACS separation buffer (see "DC expansion and isolation" for composition). Cell pellets were resuspended in 3 mL MACS separation buffer and filtered through a 70 µm pre-separation filter (Miltenyi 130-095-823) while applying the sample to the column. The sample tube and strainer were rinsed with 3 mL MACS separation buffer, and this was applied to the column; the column was rinsed two additional times with 3 mL MACS separation buffer. The column was removed from the magnetic stand and placed on a 15 mL conical tube. 5 mL MACS separation buffer was added, and bound cells were eluted by rapidly plunging. The suspension was centrifuged (5 min, 550*g*, low brake), and supernatant was aspirated.

*Live/dead* staining and *cell counting* were performed as described above under "Peptide-MHCII tetramer staining and extra/intracellular staining for flow cytometry."

*Extracellular staining:* Cells were resuspended in 150 µL flow buffer plus TruStain FcX PLUS (1:400). 150 µL of 2X surface master mix in flow buffer was added on top and incubated 20 min on ice (containing the same antibodies and concentrations listed above under "Peptide-MHCII tetramer staining and extra/intracellular staining for flow cytometry"). Cells were washed with excess flow buffer, centrifuged (5 min, 550*g*, low brake), and the supernatant was aspirated.

*Intracellular staining* and *analysis* were performed according to the same methods described above under "Peptide-MHCII tetramer staining and extra/intracellular staining for flow cytometry" except a 300 µL volume was used for both the fixation and staining steps.

**Peptide immunization and tetramer enrichment**

hCom2v-colonized mice were immunized with 50 µg poly(I:C) (Invivogen vac-pic) plus 100 µg OVA peptide (or 100 µg each of N- and C-terminal SBP peptides) IV via retro-orbital injection under ketamine/xylazine anesthesia in a biosafety cabinet using sterile technique and reagents. Ten days later, pooled spleen + LNs (inguinal, axillary, brachial, cervical, mesenteric, and peri-aortic) from each mouse were harvested and cell suspensions were generated as described above under "Cell isolation from secondary lymphoid organs," with the addition of a second filtration through a 70 µm cell strainer prior to tetramer staining. SBP tetramers conjugated to PE were produced as described under "SBP and α-Zein tetramer production"; I-$A^b$ $OVA_{328-337}$ tetramers conjugated to APC were obtained from the NIH Tetramer Core. Staining was performed as follows—see "Peptide-MHCII tetramer staining and extra/intracellular staining for flow cytometry" and "SBP tetramer enrichment from pooled hCom2v or SPF mice" for detailed reagent information, which was the same as described below except as noted:

*Peptide-MHCII tetramer staining:* The cell pellet was resuspended in ~175 µL per mouse pre-warmed RPMI 10% FCS 50 nM dasatinib plus TruStain FcX PLUS and incubated for 5 min. 10 µL/sample OVA-APC tetramer and 5 µL/sample SBP N- and C-PE tetramers were added. Cells were stained in a 37°C water bath for 2 h, gently mixing by tapping the side of the tube every 20 min. Cells were washed with excess flow buffer, centrifuged (5 min, 550*g*, low brake), and the supernatant was aspirated.

*MACS bead staining and MACS enrichment, live/dead staining*, *cell counting*, *extracellular staining*, *intracellular staining*, and *analysis* were performed as described under "SBP tetramer enrichment from pooled hCom2v or SPF mice" except a 200 µL total volume was used for the extracellular and intracellular staining steps.

**Measurement of colon length**

For colonic length measurements, the cecum and colon were carefully dissected free and fat was removed. Length from the cecal-colonic junction to the rectum was measured with a ruler and manually recorded.

**Acute DSS colitis**

Dextran sodium sulfate (DSS; 36-50 kDa, colitis grade; MP Biomedicals 0216011080) was dissolved in autoclaved MilliQ water (7.5 g per 500 mL for 1.5% DSS or 10 g per 500 mL for 2%), then filter sterilized and stored at 4°C until use. Prior to treatment, mice were weighed and fecal samples were obtained and frozen for metagenomic sequencing. Mice were given DSS solution or regular autoclaved drinking water (for untreated controls) *ad libitum* for 7 days, then were switched to regular water for 3 days and analyzed at day 10. Mice were monitored daily starting on experiment day 3 according to the following scale (*22*): Body weight loss (relative to pre-treatment weight): 0 = No negative change in weight; 1 = 1-5% loss of body weight; 2 = 5-10% loss of body weight; 3 = 10-20% loss of body weight; 4 = >20% loss of body weight; Stool consistency: 0 = Normal; 1 = Loose consistency; 2 = Watery; 3 = Slimy diarrhea; 4 = Severe diarrhea; Blood presence in stool: 0 = No blood; 2 = Red feces; 4 = Visible bleeding. The sum of these three scores for each mouse on each day was reported as the composite colitis score. hCom2v-colonized mice were handled with sterile gloves and instruments in a biosafety cabinet throughout these experiments.

**_Citrobacter rodentium_ infection**

*C. rodentium* DBS100 was obtained from the ATCC (51459) and *C. rodentium*-OVA (*23*) (DBS100 derivative) was obtained from A. Malik. Both strains were propagated in LB broth (37°C in a shaking incubator) or on MacConkey agar plates (37°C in a plate incubator; Sigma M7408-500G) and validated by whole-genome sequencing (Plasmidsaurus). *C. rodentium*-OVA was used for all infections in this study. The presence of the OVA transgene was confirmed by whole-genome sequencing; however, we detected neither stimulation of an OT-II hybridoma *in vitro* by co-culture assay nor OVA tetramer$^+$ cells after infection *in vivo*. Our analysis therefore focused on SBP tetramer$^+$ and tetramer$^-$ bulk CD4$^+$ T cells. To determine CFU per mL of culture for mouse infection experiments, a 5 mL starter culture was inoculated with a single colony and incubated overnight. The next day, 2 mL was diluted into 200 mL fresh LB and incubated at 37°C in a shaking incubator. Samples were collected at 2, 3, 4, 5, 6, and 7 h, serially diluted in PBS, and 100 µL of each dilution was plated on a MacConkey agar plate. The next day, plates with ~20-200 colonies were counted and CFU/mL calculated using the equation CFU/mL = CFU counted x dilution factor x 1/volume plated (mL). A standard curve of $OD_{600}$ vs CFU/mL was plotted, combining data from the two *C. rodentium* strains (both strains showed indistinguishable growth kinetics *in vitro*), and used to calculate the infectious dose for subsequent *in vivo* experiments.

For infection of hCom2v-colonized mice, *C. rodentium*-OVA was streaked on MacConkey agar and a single colony was inoculated into a 5 mL LB starter culture and incubated overnight. The next day, 1 mL of the overnight culture was diluted into 100 mL fresh LB broth, incubated at 37°C in a shaking incubator for ~4 h, and the $OD_{600}$ was measured and compared to the previously generated standard curve. An appropriate culture volume was centrifuged (15 min, ~4,000*g*), the supernatant was aspirated, and the pellet was resuspended in sterile PBS at $1 \times 10^{10}$ CFU/mL. Mice were inoculated with $2 \times 10^9$ CFU (200 µL) by oral gavage in a biosafety cabinet using sterile gloves and reagents.

To determine CFU per g of feces in infected mice (or uninfected controls), feces were collected in sterile tubes, weighed, and 50 µL sterile PBS was added per 10 mg feces (giving a 20% w/v solution). Samples were homogenized by vortexing vigorously (tubes were taped horizontally on the vortex and vortexed for 1 min). Large debris was pelleted by centrifugation (1 min, 400*g*). The supernatant was serially diluted in sterile PBS, and 100 µL of each dilution was plated on a MacConkey agar plate and incubated overnight at 37°C. The next day, CFU on plates for dilutions with ~20-200 colonies were counted and CFU per gram of feces was calculated using the equation CFU/g of feces = CFU counted x dilution factor x 1/volume plated (mL) x 5 (to convert from the 20% w/v suspension). Separate samples from day 0 (pre-infection) or day 13 mice were processed for metagenomic sequencing as described under "Metagenomic sequencing and analysis."

Infected hCom2v-colonized mice or uninfected controls were analyzed at day 13 post-infection according to the methods described under "Measurement of colon length," "Intestinal lamina propria (LP) lymphocyte isolation," and "Peptide-MHCII tetramer staining and extra/intracellular staining for flow cytometry."

**Metagenomic sequencing and analysis**

Small intestinal or colonic contents or fecal pellets obtained from mice were collected into sterile tubes and frozen, then processed for metagenomic sequencing. For small intestinal samples, genomic DNA was extracted using the QIAamp DNA Host-Free Microbiome kit (Qiagen) to remove mouse DNA from the sample. For colonic contents and fecal pellets, genomic DNA was extracted using the DNeasy PowerSoil Pro kit (Qiagen). DNA was quantified using the Qubit 4 or Quant-iT PicoGreen dsDNA Assay kit (Thermo Fisher). When possible, a minimum of 15 ng of DNA was used to construct metagenomic sequencing libraries. Some small intestinal samples yielded < 15 ng; in these cases, the library input was as close to 15 ng as possible. Libraries were constructed using the Illumina DNA Prep kit, with half-volumes used at each step to minimize cost. Following PCR, libraries were purified using a 0.8x bead cleanup and were quantified again using Qubit. Equal masses of each metagenomics library were pooled, except when small intestinal samples were mass limited. A dual-sided AMPure XP (Beckman Coulter) bead cleanup was performed on the pooled material to remove contaminants and to size-select for fragments between 300-1500 bp. The final library pool was quality-checked for size distribution using the Tapestation 2200 (Agilent Technologies). Sequencing was performed on the NovaSeq X or NextSeq500 (Illumina) using a 2 x 150 bp read configuration, targeting 20-30 million reads per sample. Raw reads were processed using NinjaMap (*24*) to determine the relative abundance of each strain in hCom2v. For *C. rodentium* infection experiments, the DBS100 strain genome was included in a database with the hCom2v strains and processed using the same NinjaMap pipeline. To determine community composition in samples from SPF-Tac or SPF-Jax mice, samples were prepared and sequenced as described above and analyzed using MetaPhlAn v4.1.0.

*Detection of SBP homologs in metagenomes from SPF mice:* Here, following read QC and host contamination removal, metagenomes were assembled using MEGAHIT v1.2.9 (*25*). Following binning and classification using MetaBAT2 (*26*) and GTDB-Tk (*27*) v2.6.1, respectively, we identified SBP homologs using tblastn with the *Tyzzerella nexilis* SBP protein sequence. Sequences of proteins with >70% amino acid identity were retrieved and aligned to N- and C-terminal epitope peptides using Geneious Prime. The phylogenetic tree in Supplementary Fig. 14d was generated using phyloT v2 (https://phylot.biobyte.de/).

**hCom2v F1 and GF scRNAseq and scTCRseq**

Lymphocytes were isolated from SI LP, LI LP, PP, and spleen of four hCom2v-colonized F1 mice or SI LP, LI LP, and PP of four GF mice as described above under "Intestinal lamina propria (LP) lymphocyte isolation" and "Cell isolation from secondary lymphoid organs," pooling the four mice within each tissue and cohort. Cells were resuspended in PBS 2% FCS containing TruStain FcX PLUS (anti-mouse CD16/32 S17011E; BioLegend 156604; 1:400), incubated 10 min on ice, then a 2X master mix containing the following antibody-fluorophore conjugates was added and incubated for 20 min on ice: TCRβ-APC (H57-597; BioLegend 109212; 1:200), CD4-PE (GK1.5; BioLegend 100408; 1:200), and CD45-FITC (I3/2.3; BioLegend 147710; 1:200). Cells were washed, centrifuged (5 min, 400*g*), supernatant was aspirated, and cells were resuspended in PBS 2% FCS. Just prior to sorting, DAPI (BioLegend 422801) was added to a final concentration of 0.1 μg/mL and the sample was mixed by vortexing gently. A 1.5 mL sterile collection tube was pre-coated by adding 1 mL PBS 20% FCS and then removing the liquid, then 20 μL was added to collect sorted cells. ~5 x $10^4$ $CD4^+$ $TCR\beta^+$ $CD45^+$ $DAPI^-$ cells were sorted from each sample into individual tubes using a 100 μm tip on a BD FACSAria II cell sorter. After sorting, cells were concentrated by centrifuging (10 min, 550*g*) and removing most of the supernatant by pipetting. Cells were resuspended in the remaining volume, and post-sort purity was verified and cells counted using counting beads (Spherotech). ~1 x $10^4$ cells per sample were loaded onto a chip (10x Genomics) and processed using the 10x Genomics Immune Profiling platform. Library preparation was carried out according to the manufacturer's instructions. Sequencing data were aligned using Cell Ranger (10x Genomics).

Data were processed primarily using the packages Seurat v5 (*28*) and scRepertoire (*29*, *30*) in R. Cells with > 3,250 or < 750 genes per cell or > 3% mitochondrial reads were discarded. Cells with more than one *TRA* or *TRB* chain were removed using the removeMulti argument in scRepertoire. scTCRseq data were appended to metadata using the scRepertoire combineExpression function and then TCR genes were removed from scRNAseq gene expression analysis to eliminate effects of clonal expansions on downstream cell clustering. scRNAseq data were normalized via the Seurat SCTransform function. Principal component analysis (PCA) was performed using the Seurat RunPCA function with 75 principal components (selected based on elbow plot visualization). Integration of the 7 samples into a single Seurat object was performed using the Seurat IntegrateLayers function with the RPCA method.

Clustering was performed with the Seurat FindNeighbors, FindClusters, and RunUMAP functions with 75 dimensions and resolution 0.5. Seurat, scCustomize (*31*), and scRepertoire were used to visualize gene expression and TCR clonotypes, and clusters were manually annotated based on marker gene expression patterns. An Excel spreadsheet was generated listing all unique clones within each cluster from each tissue and mouse group, sorted as described under “TCR screen design.” 5’ and 3’ Golden Gate extensions and a flag indicating the presence of an internal SapI cut site were added for all TCRs.

**DSS-treated or untreated hCom2v scRNAseq and scTCRseq**

Lymphocytes were isolated from the LI LP or mLN of seven DSS-treated hCom2v-colonized mice or four untreated controls as described above under “Intestinal lamina propria (LP) lymphocyte isolation” and “Cell isolation from secondary lymphoid organs.” Cells were stained for flow cytometry using the same procedures and antibodies described under “hCom2v F1 and GF scRNAseq and scTCRseq” except CD45-FITC was replaced with CD45.2-FITC (104; BioLegend; 109806; 1:200) to avoid competition with the anti-CD45 clone present in the BioLegend TotalSeq C oligo-tagged hashing antibody cocktail, and each tissue from each mouse was stained additionally with 0.75 µg of one of Hashtag antibodies 1-10 (BioLegend 155861, 155863, 155865, 155867, 155869, 155871, 155873, 155875, 155877, or 155879; centrifuged 10 min at 14,000*g* before adding to cells). Cells were washed three times with 1.2 mL PBS 2% FCS, fully resuspending the pellet and then centrifuging (5 min, 400*g*) and aspirating the supernatant in between; following these washes, mice were pooled within a tissue for each treatment group. No $CD45^+$ cells were obtained from the mLN of one of the four untreated mice; this mouse was therefore not represented in the pooled untreated mLN sample and was excluded from analyses. Cells were sorted and counted as described under “hCom2v F1 and GF scRNAseq and scTCRseq”, then loaded onto a 10x Genomics chip targeting a recovery of $2 \times 10^4$ cells per sample. Libraries were prepared according to the manufacturer’s instructions for the 10x Genomics Immune Profiling platform, and sequencing data were aligned using Cell Ranger (10x Genomics).

Data were processed similarly to that described under “hCom2v F1 and GF scRNAseq and scTCRseq”. Hashing oligo data were normalized using the Seurat NormalizeData command and demultiplexed using the HTODemux function with positive.quantile = 0.99; only singlets were retained. Cells with < 1,000 or > 4,000 genes per cell, > 3.9% mitochondrial reads, or 0% mitochondrial reads (likely representing technical artifacts) were discarded. Cells with more than one *TRA* or *TRB* chain were removed using the removeMulti argument in scRepertoire, and scTCRseq data were combined with scRNAseq data using the scRepertoire combineExpression function. scRNAseq data were normalized via the Seurat SCTransform function, TCR genes were removed from gene expression analysis using the scRepertoire quietTCRgenes function, and PCA was performed with 75 principal components. The four samples were integrated using the Seurat IntegrateLayers function with the RPCA method.

Clustering was performed using the Seurat FindNeighbors, FindClusters, and RunUMAP commands with 75 dimensions and a resolution of 0.55. Clusters were manually annotated based on expression of marker genes. For pseudobulk gene expression analysis, the Seurat AggregateExpression function was applied and then differentially expressed markers across treatment conditions identified using the FindMarkers function with the DESeq2 test, using individual mice as replicates.

**Statistical analysis**

Statistical tests were performed as indicated in figure legends or the manuscript text using GraphPad Prism (v11), R (v4.4.1), or Microsoft Excel. Details of statistical tests, the value of $n$, and the number of independent experiments are provided in the figures and corresponding figure legends.

**Use of artificial intelligence (AI)-assisted technologies**

During preparation of this manuscript, AI-assisted tools (OpenAI ChatGPT, Anthropic Claude, and Google Gemini) were used for language and clarity editing of author-written text and for assistance with analysis code, including debugging and, in some cases, drafting of code that was then reviewed, tested, and validated by the authors. The authors take full responsibility for all AI-assisted output and content.

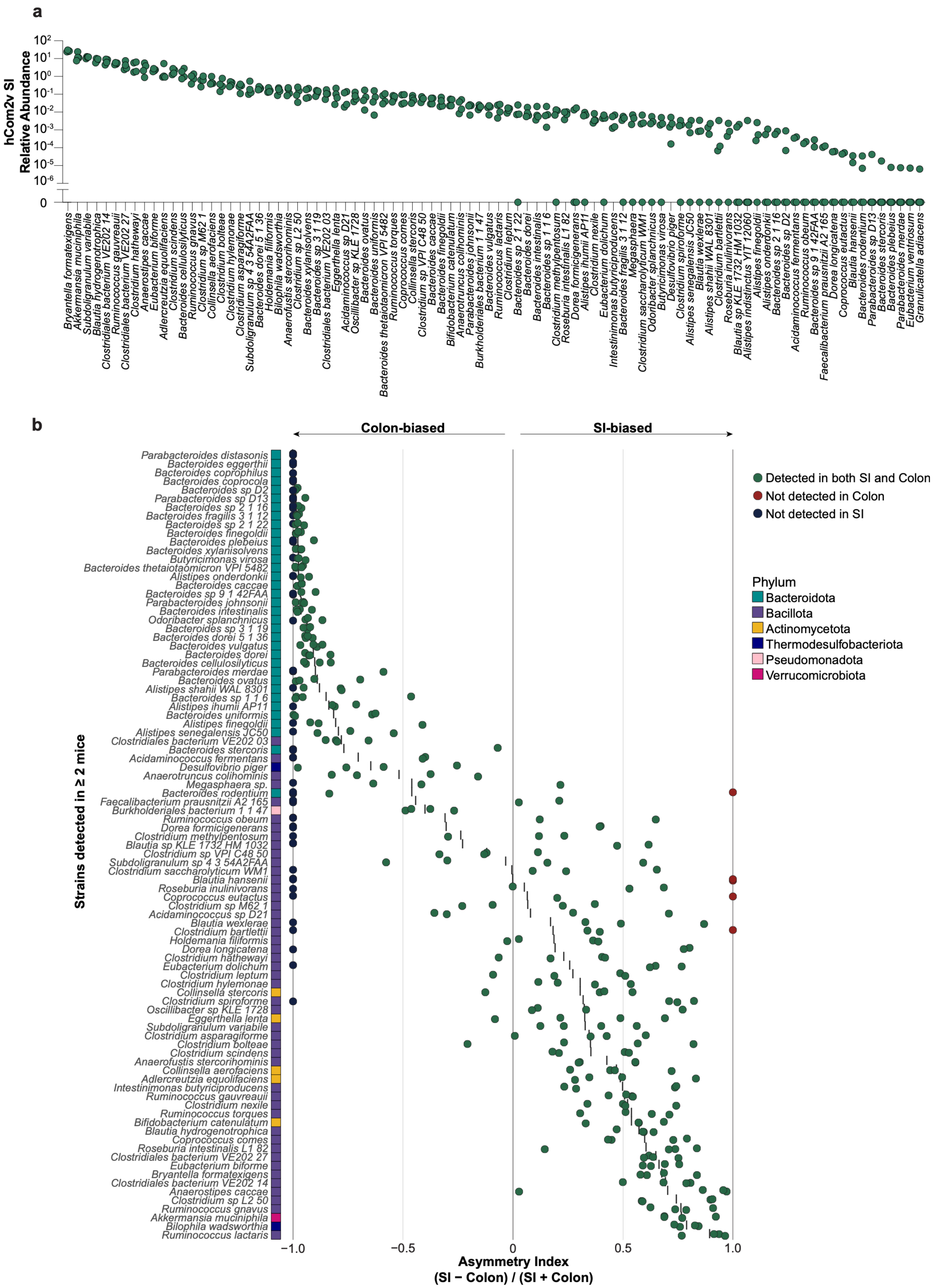

a
hCom2v SI Relative Abundance
b
Colon-biased
SI-biased
Strains detected in ≥ 2 mice
Asymmetry Index
(SI − Colon) / (SI + Colon)
Detected in both SI and Colon
Not detected in Colon
Not detected in SI
Phylum
Bacteroidota
Bacillota
Actinomycetota
Thermodesulfobacteriota
Pseudomonadota
Verrucomicrobiota

**Supplementary Figure 1: SI community composition and regional distribution of strains in hCom2v F1 mice.** (**a**) Relative abundance of individual strains in the SI of hCom2v F1 mice by metagenomic sequencing. Each dot represents an individual mouse (*n* = 4), and only strains detected in the SI or colon of at least one mouse are shown. (**b**) SI or colon asymmetry index, calculated as (SI – colon) / (SI + colon), for individual strains in hCom2v F1 mice. Each dot indicates an individual mouse, and only strains detected in ≥ 2 mice are shown. Dot colors indicate strains that were only detected in the colon (blue), only in the SI (red), or in both compartments (green). Colors adjacent to each strain name indicate the phylum.

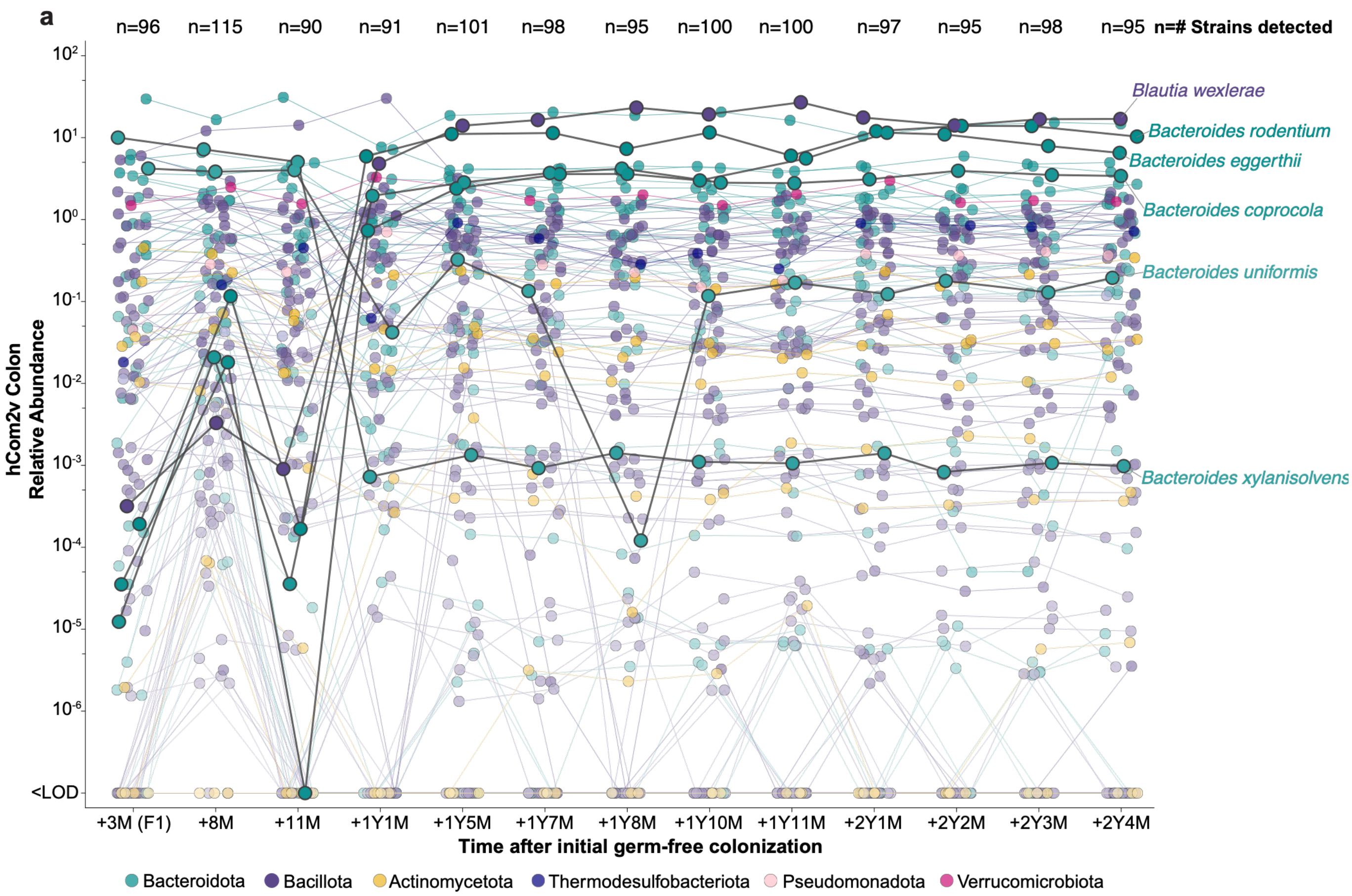


**Supplementary Figure 2: hCom2v composition over time in the breeding colony.** (**a**) Relative abundance of individual hCom2v strains in colonic or fecal samples as determined by metagenomic sequencing. Each dot represents an individual strain, colored according to phylum; lines connect the same strain across time points, and selected strains with notable changes in abundance over time are highlighted. Each time point comprises 1-5 cages of mice from the breeding colony, with no cage sampled at more than one time point; strain abundances were averaged across individual mice within a time point. Time from the date of the original GF colonization in months (M) and years (Y) is shown; the number of strains detected in at least one mouse at the indicated time point is displayed above the plot. Some data shown here also appear in Fig. 1a and Supplementary Figs. 12g, 15b-c (d0 samples), and 20b (unimmunized samples).

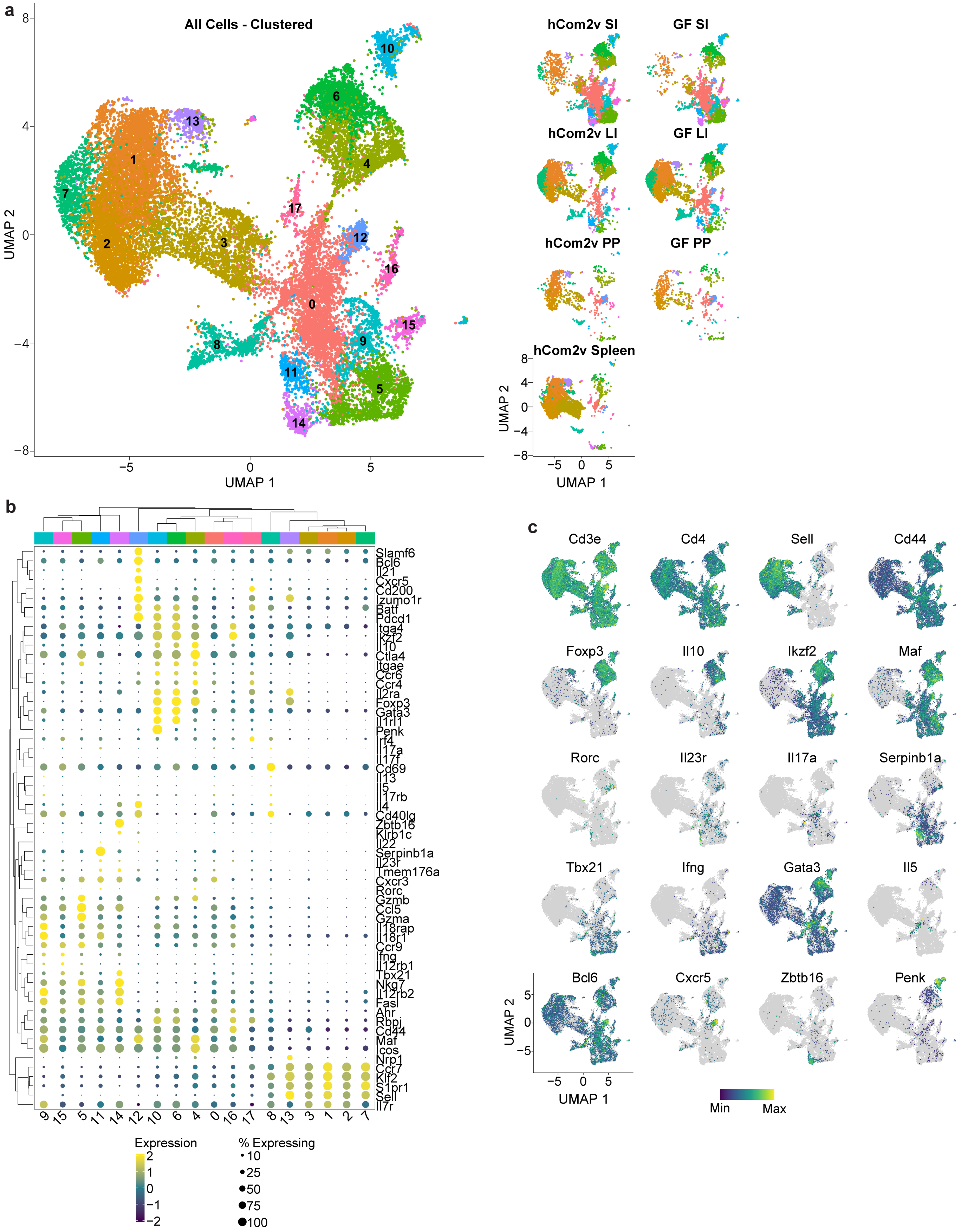
a
All Cells - Clustered
UMAP 2
UMAP 1
8
4
0
−4
−8
−5
0
5
10
6
13
1
4
7
17
2
3
12
16
0
15
8
9
11
5
14
hCom2v SI
GF SI
hCom2v LI
GF LI
hCom2v PP
GF PP
hCom2v Spleen
b
Slamf6
Bcl6
Il21
Cxcr5
Cd200
Izumo1r
Batf
Pdcd1
Itga4
Ikzf2
Il10
Ctla4
Itgae
Ccr6
Ccr4
Il2ra
Foxp3
Gata3
Il1rl1
Penk
Irf4
Il17a
Il17f
Cd69
Il13
Il5
Il17rb
Il4
Cd40lg
Zbtb16
Klrb1c
Il22
Serpinb1a
Il23r
Tmem176a
Cxcr3
Rorc
Gzmb
Ccl5
Gzma
Il18rap
Il18r1
Ccr9
Ifng
Il12rb1
Tbx21
Nkg7
Il12rb2
Fasl
Ahr
Rbpj
Cd44
Maf
Icos
Nrp1
Ccr7
Klf2
S1pr1
Sell
Il7r
9 15 5 11 14 12 10 6 4 0 16 17 8 13 3 1 2 7
Expression
2
1
0
−1
−2
% Expressing
10
25
50
75
100
c
Cd3e
Cd4
Sell
Cd44
Foxp3
Il10
Ikzf2
Maf
Rorc
Il23r
Il17a
Serpinb1a
Tbx21
Ifng
Gata3
Il5
Bcl6
Cxcr5
Zbtb16
Penk
Min
Max

**Supplementary Figure 3: Analysis of $CD4^+$ T cell clusters in hCom2v F1 and GF mice by scRNAseq.** (**a**) UMAP plot depicting unannotated $CD4^+$ T cell clusters identified by scRNAseq across all tissues combined (*left*) or subset by tissue and colonization status (hCom2v or GF; *right*); four mice were pooled per group. See Supplementary Fig. 4 for annotated clusters. (**b**) Clustered dot plot and (**c**) feature plots displaying expression patterns of selected marker genes used for cluster annotation. See Methods for clustering parameters.

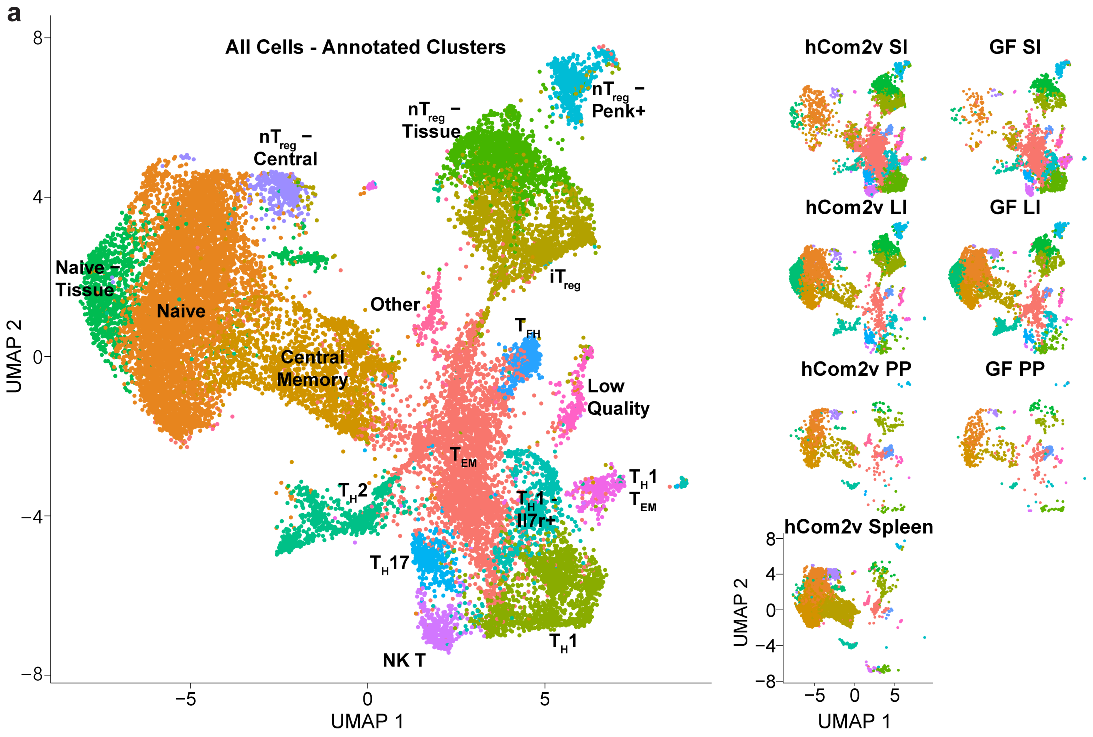


**Supplementary Figure 4: Identification of CD4⁺ T cell populations in hCom2v F1 and GF mice.** (**a**) UMAP plot depicting annotated CD4⁺ T cell clusters identified by scRNAseq across all tissues combined (*left*) or subset by tissue and colonization status (hCom2v or GF; *right*); four mice were pooled per group. See Supplementary Fig. 3 for unannotated clusters and the marker genes used for annotation.

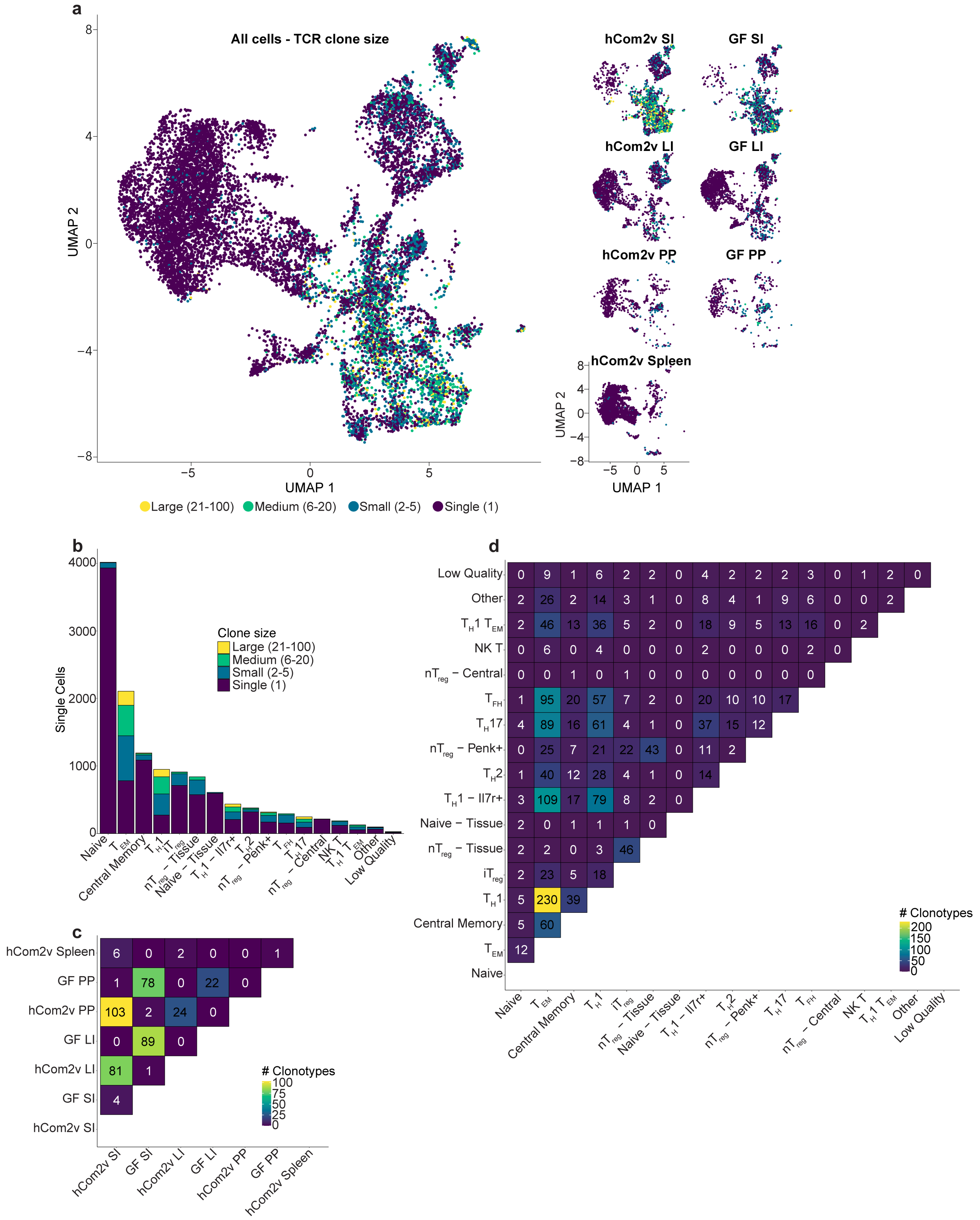

a
All cells - TCR clone size
UMAP 2
UMAP 1
Large (21-100)
Medium (6-20)
Small (2-5)
Single (1)
hCom2v SI
GF SI
hCom2v LI
GF LI
hCom2v PP
GF PP
hCom2v Spleen
b
Single Cells
Clone size
Large (21-100)
Medium (6-20)
Small (2-5)
Single (1)
Naive
T EM
Central Memory
T H 1
iT reg
nT reg – Tissue
Naive – Tissue
T H 1 – Il7r+
T H 2
nT reg – Penk+
T FH
T H 17
nT reg – Central
NK T
T H 1 T EM
Other
Low Quality
c
hCom2v Spleen 6 0 2 0 0 1
GF PP 1 78 0 22 0
hCom2v PP 103 2 24 0
GF LI 0 89 0
hCom2v LI 81 1
GF SI 4
hCom2v SI
hCom2v SI
GF SI
hCom2v LI
GF LI
hCom2v PP
GF PP
hCom2v Spleen
### Clonotypes
d
Low Quality 0 9 1 6 2 2 0 4 2 2 2 3 0 1 2 0
Other 2 26 2 14 3 1 0 8 4 1 9 6 0 0 2
T H 1 T EM 2 46 13 36 5 2 0 18 9 5 13 16 0 2
NK T 0 6 0 4 0 0 0 2 0 0 0 2 0
nT reg – Central 0 0 1 0 1 0 0 0 0 0 0 0
T FH 1 95 20 57 7 2 0 20 10 10 17
T H 17 4 89 16 61 4 1 0 37 15 12
nT reg – Penk+ 0 25 7 21 22 43 0 11 2
T H 2 1 40 12 28 4 1 0 14
T H 1 – Il7r+ 3 109 17 79 8 2 0
Naive – Tissue 2 0 1 1 1 0
nT reg – Tissue 2 2 0 3 46
iT reg 2 23 5 18
T H 1 5 230 39
Central Memory 5 60
T EM 12
Naive
### Clonotypes

**Supplementary Figure 5: TCR clonal architecture in hCom2v F1 and GF mice.** (**a**) UMAP plot with dots colored by TCR clone size (as determined by scTCRseq) across all tissues combined (*left*) or subset by tissue and colonization status (hCom2v or GF; *right*); four mice were pooled per group. (**b**) Number of TCR clonotypes with the indicated clone sizes in each cluster. (**c**) TCR clonal overlap between tissues and (**d**) between clusters; fill color indicates the number of shared clonotypes. See Supplementary Fig. 4 for annotated scRNAseq clusters.

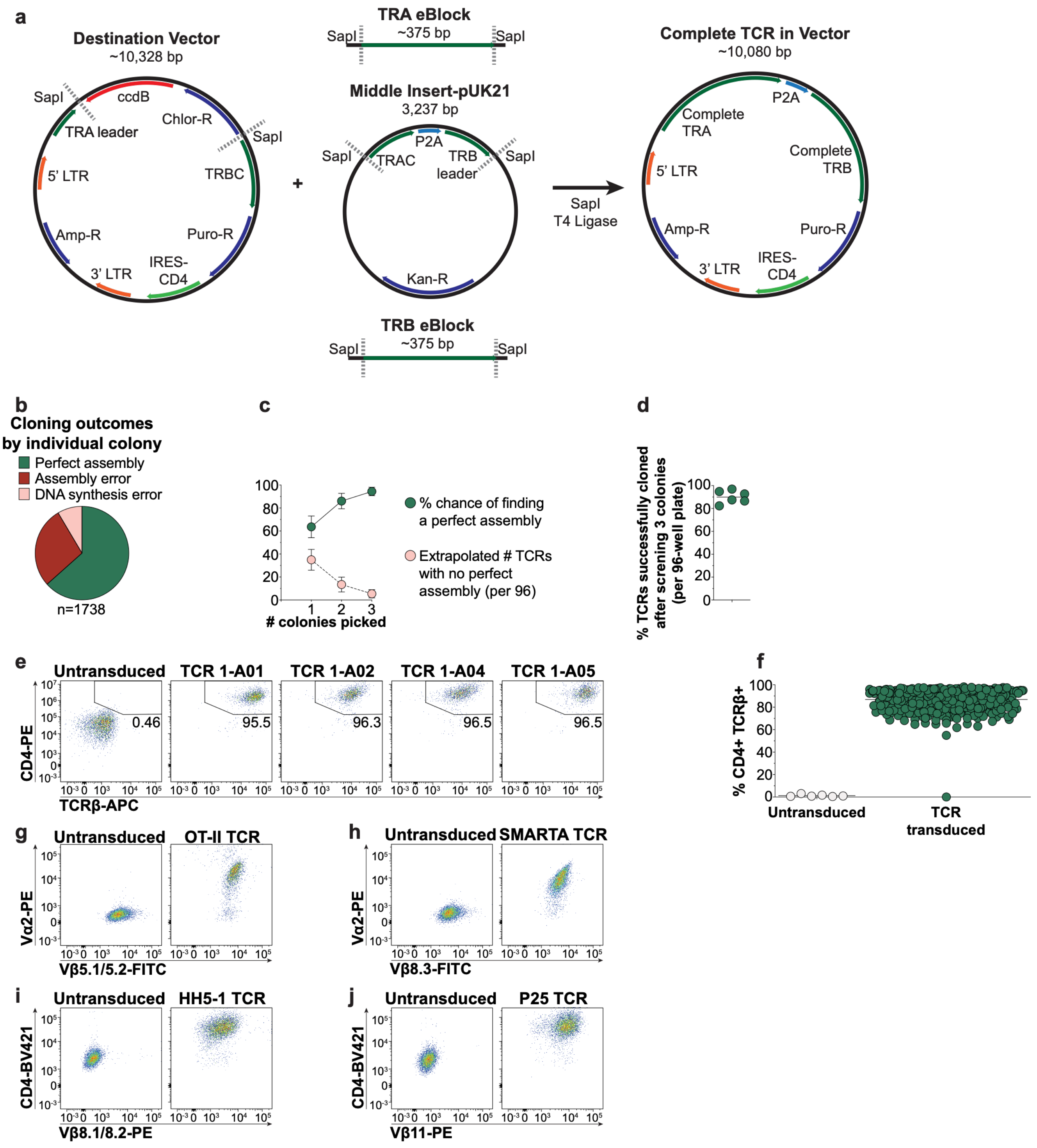


**Supplementary Figure 6: Cloning and expression of TCRs at high throughput.** (**a**) Schematic depicting the Golden Gate assembly strategy for TCR cloning. Synthesized *TRA* and *TRB* eBlocks with SapI overhangs are mixed with Destination Vector and Middle Insert Vector plasmids. Upon thermal cycling in the presence of SapI and T4 ligase, the complete TCR is ligated into a single retroviral expression vector. Notable DNA elements in each component are labeled; see Methods for additional details. (**b**) Cloning outcomes summarized across 1,738 colonies screened by miniprep and whole-plasmid sequencing. Plasmid sequences were aligned to the intended TCR variable regions and vector elements and manually scored as perfect assembly, assembly errors (incorrect ligation of the intended fragments), or DNA synthesis errors (mutations within the synthesized variable region fragments); fill color indicates the fraction of each category observed. (**c**) Percentage chance of finding a perfect assembly (green) and extrapolated number of TCRs per 96-well plate with no perfect assembly (pink), both by number of colonies screened. The percentage of colonies with perfect assemblies was tallied for each of six 96-well batches of TCRs and used to calculate the percentage chance of finding a perfect assembly using the formula $[1-(1-p)^n] \times 100$, where $p$ is the fraction of colonies with perfect assemblies and $n$ is the number of colonies screened. The extrapolated number of TCRs with no perfect assembly per 96-well plate was calculated using the formula $96 \times (1-c/100)$, where $c$ is the percentage chance from the preceding formula. Dots indicate the mean and error bars standard deviation. (**d**) Overall percentage of TCRs successfully cloned per 96-well plate after screening three colonies per TCR; each dot represents an individual 96-well batch of TCRs. (**e**) Representative flow cytometry and (**f**) summary plot depicting surface CD4 and TCRβ expression in the untransduced and unselected parental 58αβ$^{-/-}$ T cell hybridoma line or in TCR-expressing lines following retroviral transduction and puromycin selection. Each dot represents an individual cell line, and horizontal lines indicate the mean; data compiled from six independent experiments. (**g-j**) Flow cytometry plots depicting surface staining of T cell hybridoma lines expressing the OT-II, SMARTA, HH5-1, or P25 control TCRs or the parental 58αβ$^{-/-}$ T cell hybridoma line with the indicated Vα- and Vβ-specific antibodies or CD4. Data are from a single experiment.

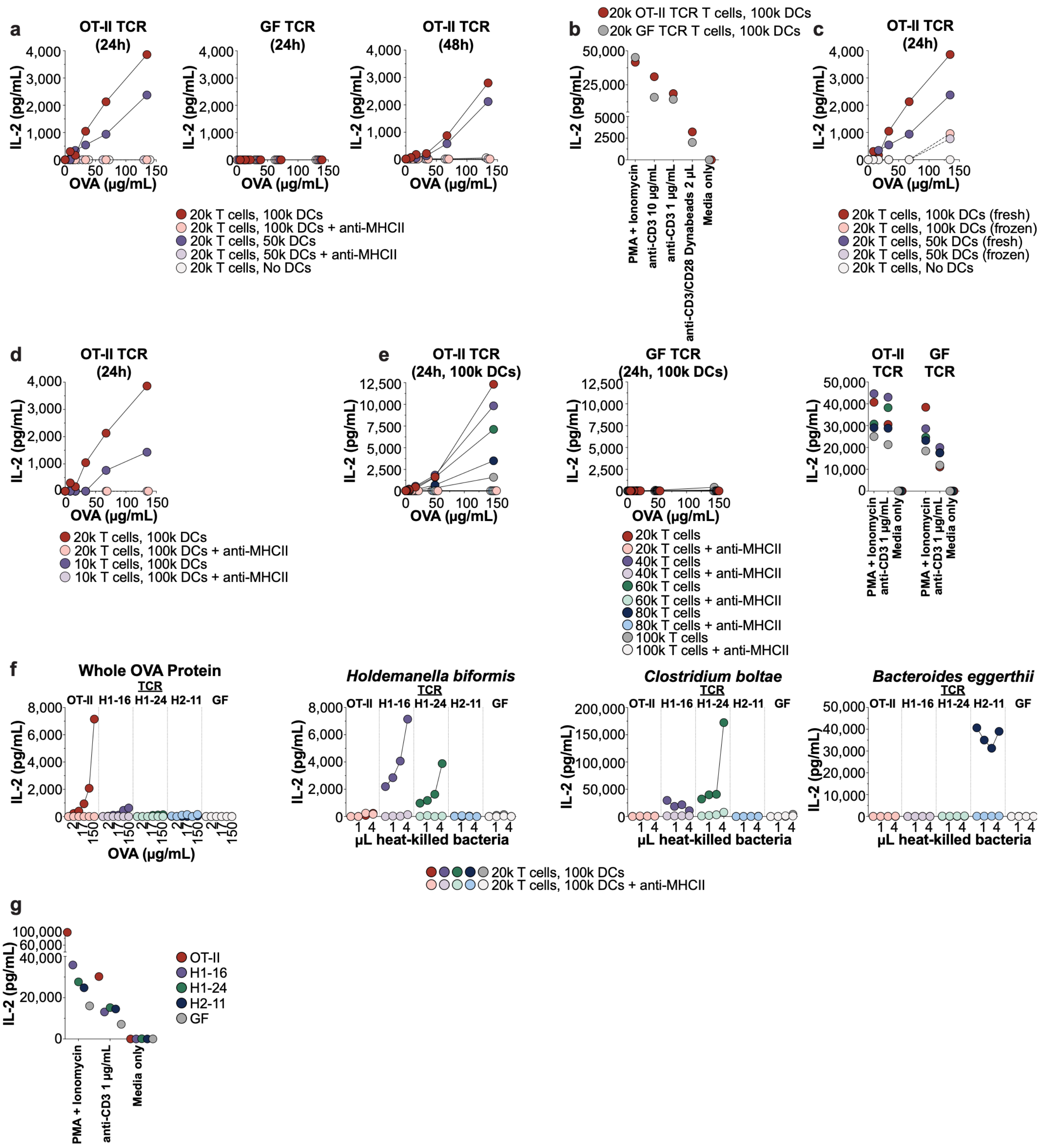


**Supplementary Figure 7: Optimization of a 384-well co-culture assay for TCR specificity.** (**a**) Titration of DC cell numbers with 2 x $10^4$ cells expressing the OT-II TCR or a non-reactive control TCR from a GF mouse (GF) in the presence of the indicated concentrations of whole OVA with or without an anti-MHCII blocking antibody and co-cultured for 24 h (*left*) or 48 h (*right*). (**b**) Response of the OT-II and GF TCRs to positive control stimuli (PMA/ionomycin, soluble anti-CD3ε at the indicated concentrations, or anti-CD3/CD28 Dynabeads) or medium alone. (**c**) Titration of DC cell numbers with a fixed 2 x $10^4$ OT-II or GF TCR-expressing cells in the presence of the indicated concentrations of whole OVA. DCs were either prepared fresh or cryopreserved (frozen). (**d**) Signal obtained with 1 x $10^4$ vs 2 x $10^4$ OT-II or GF TCR-expressing cells in the presence of 1 x $10^5$ DCs and the indicated concentrations of whole OVA, with or without an anti-MHCII blocking antibody. Data shown in a-d are from a single experiment and some data points are reproduced across panels to facilitate comparison. (**e**) Titration of increasing numbers of OT-II or GF TCR-expressing cells with a fixed 1 x $10^5$ DCs and the indicated concentrations of whole OVA, with or without an anti-MHCII blocking antibody (*left*) and response to positive or negative control stimuli (*right*). Data are from a single experiment. (**f**) Co-culture of 2 x $10^4$ of the indicated TCR-expressing cells and 1 x $10^5$ DCs against the indicated concentration of whole OVA or the indicated volume (in µL) of normalized autoclaved bacterial strains, with or without an anti-MHCII blocking antibody. (**g**) Response of the same TCRs to positive or negative control stimuli. Data in f-g are from a single experiment. Co-cultures were incubated for ~24 h except where indicated, and IL-2 concentrations in all panels were measured by AlphaLISA.

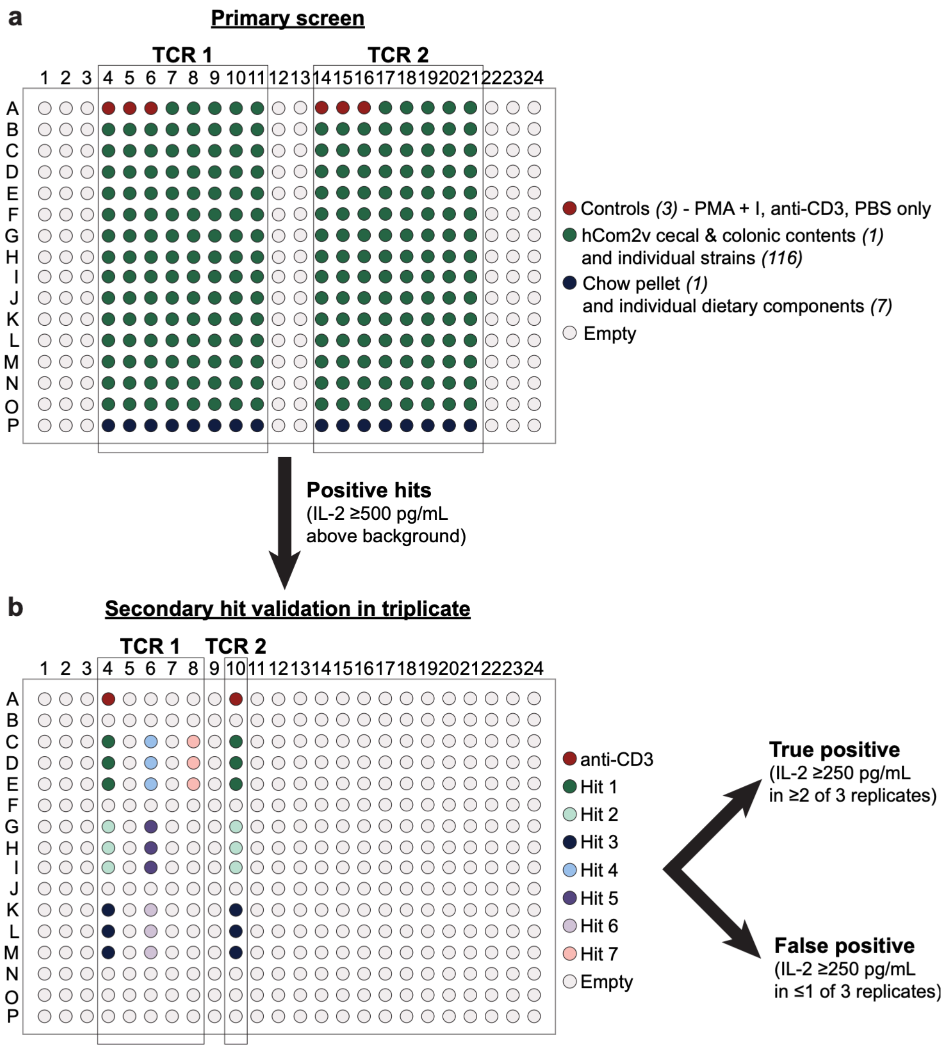


**Supplementary Figure 8: Plate layouts for the primary screen and secondary hit validation.** (**a**) Plate layout for the primary screen. Each TCR was assayed against 128 stimuli by co-culture: three controls (PMA/ionomycin, anti-CD3ε, or PBS-only), pooled hCom2v cecal and colonic contents, the 116 individual bacterial strains in hCom2v, and the chow pellet and seven proteinaceous subcomponents. Two TCRs were screened per 384-well plate as indicated. (**b**) Positive hits in the primary screen (IL-2 ≥ 500 pg/mL) were re-tested in an independent secondary validation assay. Each hit was re-tested in triplicate in plate layouts designed to minimize spillover from adjacent wells; example layouts for two TCRs are shown. Stimuli showing IL-2 ≥ 250 pg/mL in ≥ 2 wells were considered true positives. See Methods for additional details regarding the design and execution of the screen.

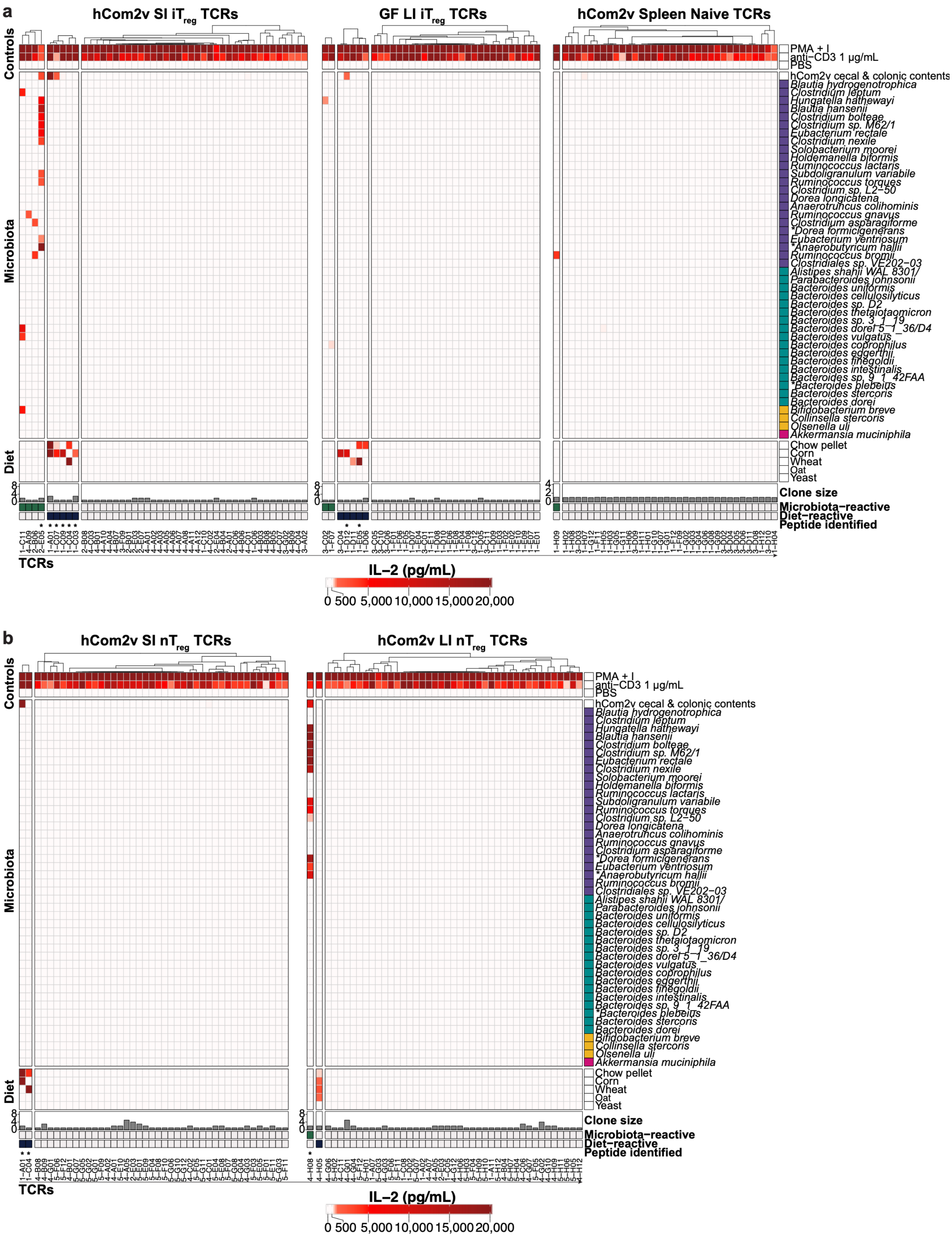

a
hCom2v SI iTreg TCRs
GF LI iTreg TCRs
hCom2v Spleen Naive TCRs
Controls
Microbiota
Diet
PMA + I
anti−CD3 1 μg/mL
PBS
hCom2v cecal & colonic contents
Blautia hydrogenotrophica
Clostridium leptum
Hungatella hathewayi
Blautia hansenii
Clostridium bolteae
Clostridium sp. M62/1
Eubacterium rectale
Clostridium nexile
Solobacterium moorei
Holdemanella biformis
Ruminococcus lactaris
Subdoligranulum variabile
Ruminococcus torques
Clostridium sp. L2−50
Dorea longicatena
Anaerotruncus colihominis
Ruminococcus gnavus
Clostridium asparagiforme
*Dorea formicigenerans
Eubacterium ventriosum
*Anaerobutyricum hallii
Ruminococcus bromii
Clostridiales sp. VE202−03
Alistipes shahii WAL 8301/
Parabacteroides johnsonii
Bacteroides uniformis
Bacteroides cellulosilyticus
Bacteroides sp. D2
Bacteroides thetaiotaomicron
Bacteroides sp. 3_1_19
Bacteroides dorei 5_1_36/D4
Bacteroides vulgatus
Bacteroides coprophilus
Bacteroides eggerthii
Bacteroides finegoldii
Bacteroides intestinalis
Bacteroides sp. 9_1_42FAA
*Bacteroides plebeius
Bacteroides stercoris
Bacteroides dorei
Bifidobacterium breve
Collinsella stercoris
Olsenella uli
Akkermansia muciniphila
Chow pellet
Corn
Wheat
Oat
Yeast
Clone size
Microbiota−reactive
Diet−reactive
Peptide identified
TCRs
IL−2 (pg/mL)
0 500 5,000 10,000 15,000 20,000
b
hCom2v SI nTreg TCRs
hCom2v LI nTreg TCRs
Controls
Microbiota
Diet
PMA + I
anti−CD3 1 μg/mL
PBS
hCom2v cecal & colonic contents
Blautia hydrogenotrophica
Clostridium leptum
Hungatella hathewayi
Blautia hansenii
Clostridium bolteae
Clostridium sp. M62/1
Eubacterium rectale
Clostridium nexile
Solobacterium moorei
Holdemanella biformis
Ruminococcus lactaris
Subdoligranulum variabile
Ruminococcus torques
Clostridium sp. L2−50
Dorea longicatena
Anaerotruncus colihominis
Ruminococcus gnavus
Clostridium asparagiforme
*Dorea formicigenerans
Eubacterium ventriosum
*Anaerobutyricum hallii
Ruminococcus bromii
Clostridiales sp. VE202−03
Alistipes shahii WAL 8301/
Parabacteroides johnsonii
Bacteroides uniformis
Bacteroides cellulosilyticus
Bacteroides sp. D2
Bacteroides thetaiotaomicron
Bacteroides sp. 3_1_19
Bacteroides dorei 5_1_36/D4
Bacteroides vulgatus
Bacteroides coprophilus
Bacteroides eggerthii
Bacteroides finegoldii
Bacteroides intestinalis
Bacteroides sp. 9_1_42FAA
*Bacteroides plebeius
Bacteroides stercoris
Bacteroides dorei
Bifidobacterium breve
Collinsella stercoris
Olsenella uli
Akkermansia muciniphila
Chow pellet
Corn
Wheat
Oat
Yeast
Clone size
Microbiota−reactive
Diet−reactive
Peptide identified
TCRs
IL−2 (pg/mL)
0 500 5,000 10,000 15,000 20,000

**Supplementary Figure 9: TCR reactivity heatmaps for additional iT$_{reg}$, naïve, and nT$_{reg}$ populations.** (**a**) Heatmaps depicting reactivity patterns of TCRs from SI iT$_{regs}$ from hCom2v-colonized mice, LI iT$_{regs}$ from GF mice, and splenic naïve T cells from hCom2v-colonized mice. (**b**) Heatmaps depicting reactivity patterns of TCRs from SI and LI nT$_{regs}$ from hCom2v-colonized mice. In a and b, each column represents a unique TCR with its identifier listed on the x-axis; each row represents an individual stimulus. Dendrograms depict hierarchical clustering of columns by Spearman correlation distance with Ward's D2 method. Cell color indicates IL-2 concentration in pg/mL measured by AlphaLISA following co-culture assay. Each TCR was screened against 128 stimuli, and control stimuli plus those against which at least one TCR had a validated reactivity in any panel are shown. Bottom annotations indicate the clone size for each TCR within its cluster, whether TCRs were scored as microbiota-reactive (green), diet-reactive (blue), or non-reactive (gray), and asterisks indicate TCRs for which a peptide epitope was identified. Rows with an asterisk contain some missing data points. See Supplementary Table 2 for the complete dataset, Fig. 2b for reactivity summaries for each TCR panel, and Methods for the complete screening workflow.

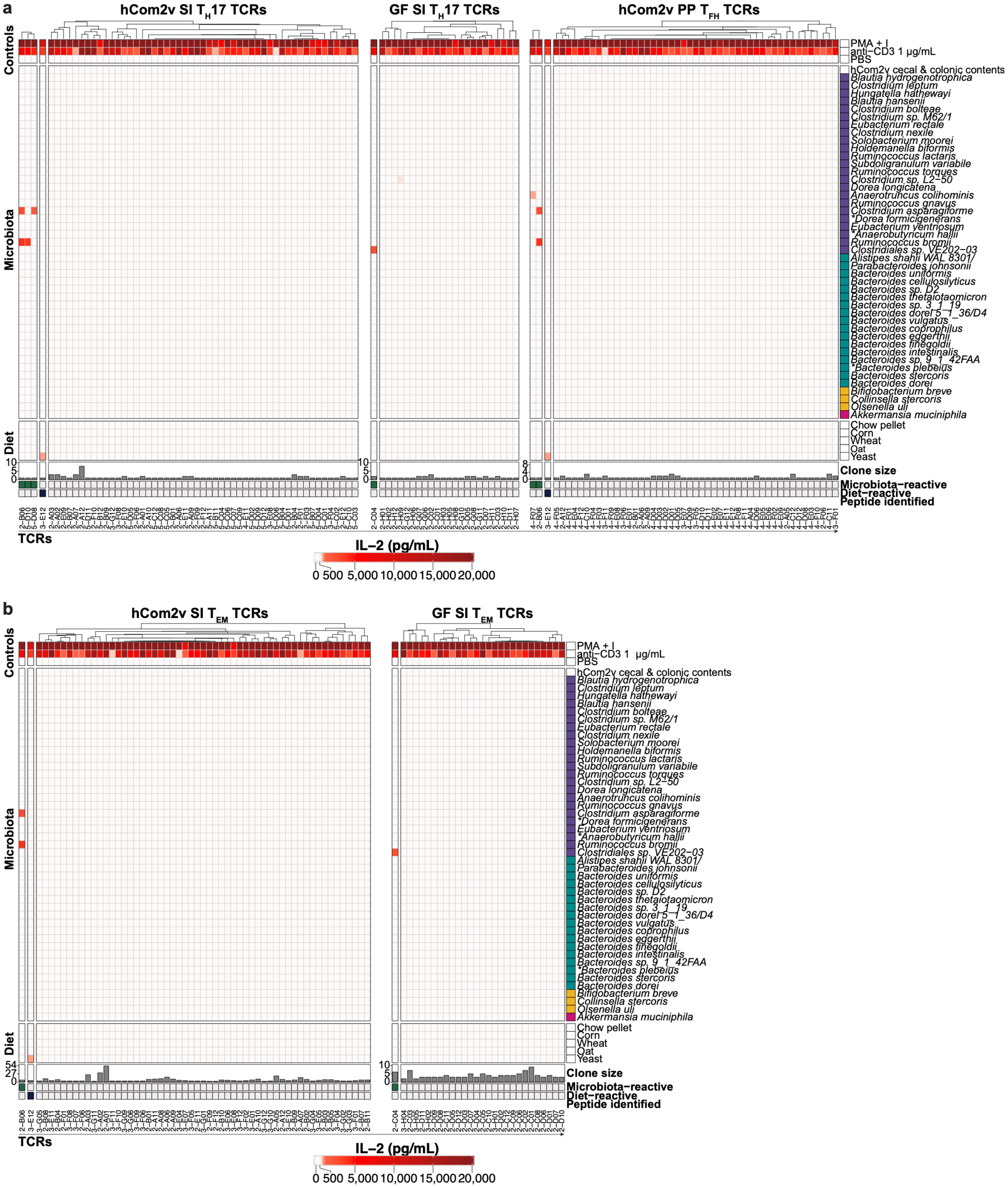


**Supplementary Figure 10: TCR reactivity heatmaps for $T_H17$, $T_{FH}$, and $T_{EM}$ populations.** (a) Heatmaps depicting reactivity patterns of TCRs from SI $T_H17$ cells from hCom2v-colonized and GF mice, and from PP $T_{FH}$ from hCom2v-colonized mice. (**b**) Heatmaps depicting reactivity patterns of TCRs from SI $T_{EM}$ cells from hCom2v-colonized and GF mice. In a and b, each column represents a unique TCR with its identifier listed on the x-axis; each row represents an individual stimulus. Dendrograms depict hierarchical clustering of columns by Spearman correlation distance with Ward's D2 method. Cell color indicates IL-2 concentration in pg/mL measured by AlphaLISA following co-culture assay. Each TCR was screened against 128 stimuli, and control stimuli plus those against which at least one TCR had a validated reactivity in any panel are shown. Bottom annotations indicate the clone size for each TCR within its cluster, whether TCRs were scored as microbiota-reactive (green), diet-reactive (blue), or non-reactive (gray), and asterisks indicate TCRs for which a peptide epitope was identified. Rows with an asterisk contain some missing data points. See Supplementary Table 2 for the complete dataset, Fig. 2b for reactivity summaries for each TCR panel, and Methods for the complete screening workflow.

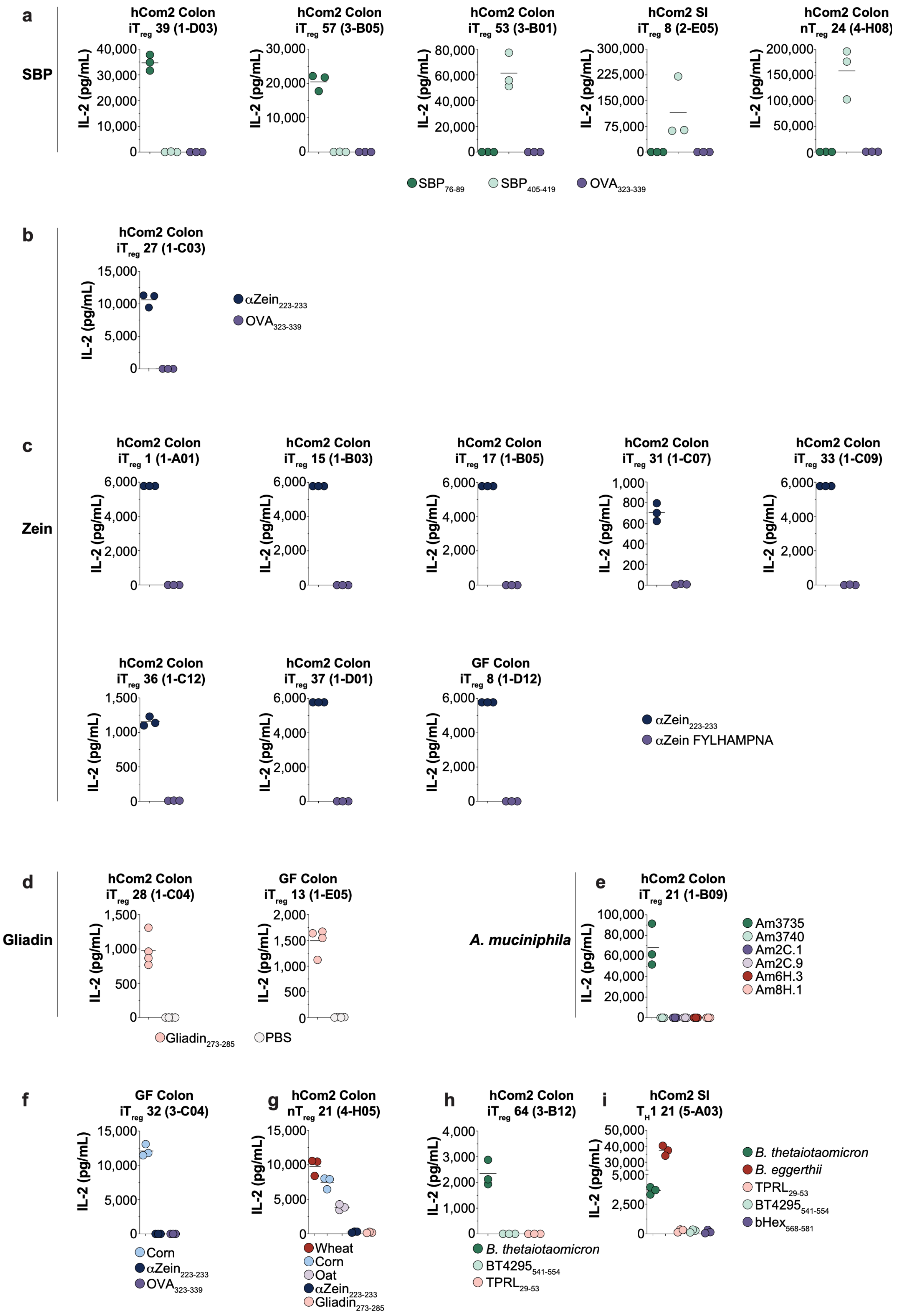

a
SBP
hCom2 Colon iTreg 39 (1-D03)
hCom2 Colon iTreg 57 (3-B05)
hCom2 Colon iTreg 53 (3-B01)
hCom2 SI iTreg 8 (2-E05)
hCom2 Colon nTreg 24 (4-H08)
IL-2 (pg/mL)
SBP76-89
SBP405-419
OVA323-339
b
hCom2 Colon iTreg 27 (1-C03)
αZein223-233
OVA323-339
c
Zein
hCom2 Colon iTreg 1 (1-A01)
hCom2 Colon iTreg 15 (1-B03)
hCom2 Colon iTreg 17 (1-B05)
hCom2 Colon iTreg 31 (1-C07)
hCom2 Colon iTreg 33 (1-C09)
hCom2 Colon iTreg 36 (1-C12)
hCom2 Colon iTreg 37 (1-D01)
GF Colon iTreg 8 (1-D12)
αZein223-233
αZein FYLHAMPNA
d
Gliadin
hCom2 Colon iTreg 28 (1-C04)
GF Colon iTreg 13 (1-E05)
Gliadin273-285
PBS
e
A. muciniphila
hCom2 Colon iTreg 21 (1-B09)
Am3735
Am3740
Am2C.1
Am2C.9
Am6H.3
Am8H.1
f
GF Colon iTreg 32 (3-C04)
Corn
αZein223-233
OVA323-339
g
hCom2 Colon nTreg 21 (4-H05)
Wheat
Corn
Oat
αZein223-233
Gliadin273-285
h
hCom2 Colon iTreg 64 (3-B12)
B. thetaiotaomicron
BT4295541-554
TPRL29-53
i
hCom2 SI TH1 21 (5-A03)
B. thetaiotaomicron
B. eggerthii
TPRL29-53
BT4295541-554
bHex568-581

**Supplementary Figure 11: Peptide epitope mapping for microbiota- and diet-reactive TCRs.** (**a**) Reactivity of the indicated TCRs against SBP N- or C-terminal peptide epitopes or OVA peptide as a control. (**b-c**) Reactivity of the indicated TCRs against the corn α-Zein C-terminal epitope, with OVA or a second α-Zein peptide as controls. (**d**) Reactivity of the indicated TCRs against a peptide epitope from gliadin or a PBS-only control. (**e**) Reactivity of the indicated TCR against a panel of peptide epitopes from *A. muciniphila*. (**f**) Reactivity of a corn-reactive TCR against the indicated epitopes from corn α-Zein or OVA. (**g**) Reactivity of a corn-, wheat-, and oat-reactive TCR against the indicated epitopes from corn α-Zein or wheat gliadin. (**h-i**) Reactivity of the indicated Bacteroidota-reactive TCRs against peptide epitopes from Bacteroidota spp. No reactive epitope was identified in f-i. For all panels, each dot represents one of 3-4 replicate wells run in parallel in a single experiment and horizontal lines indicate the mean. Panels a, b, and e-i display co-culture assays run in 384-well format with IL-2 measured by AlphaLISA; panels c-d show co-culture assays run in 96-well format with IL-2 measured by ELISA. In TCR identifiers, “Colon” denotes the LI LP. See Methods for peptide sequences and Supplementary Table 2 for the complete dataset.

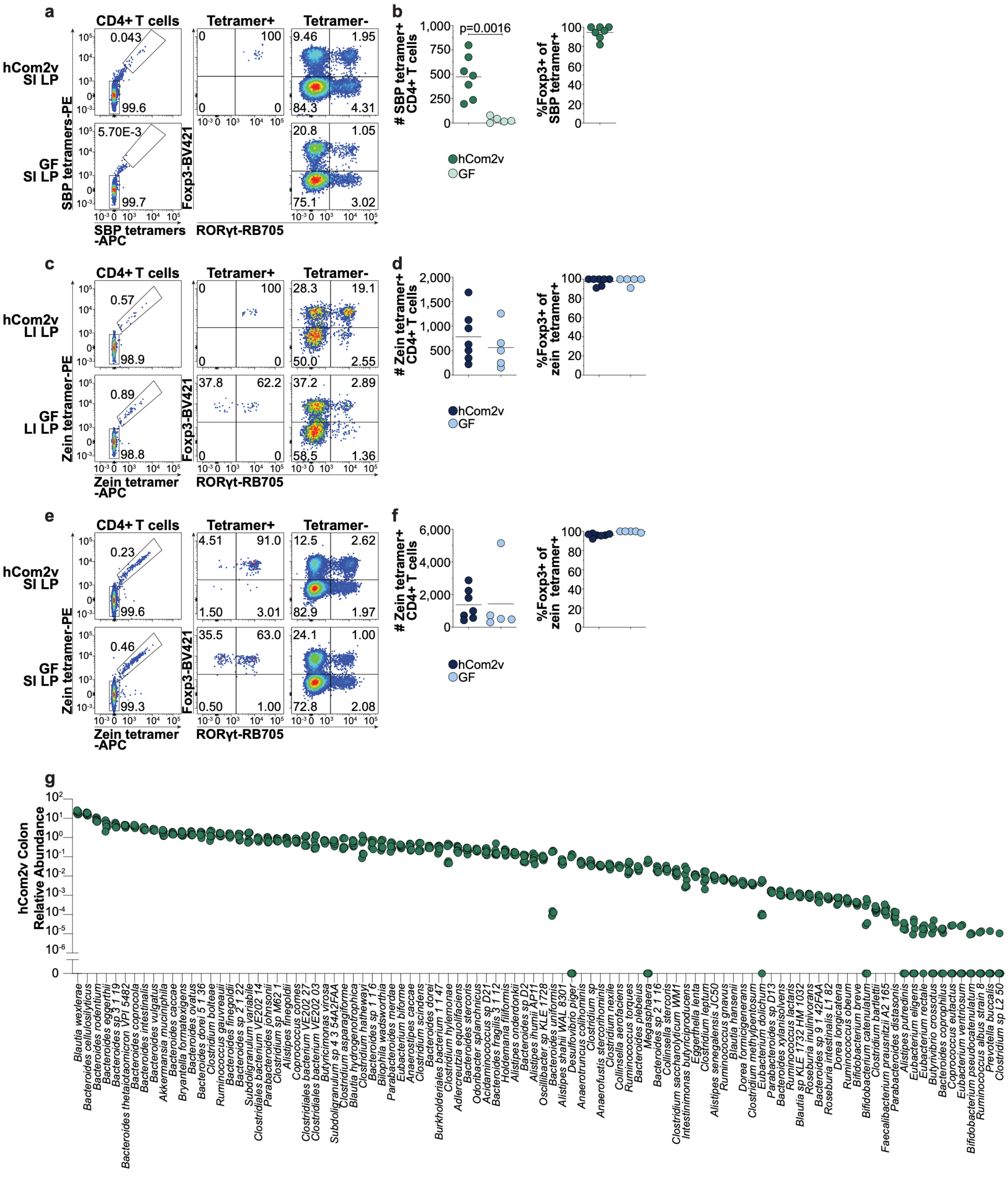


**Supplementary Figure 12: Additional SBP and α-Zein tetramer staining in hCom2v-colonized and GF mice.** (**a**) Representative flow cytometry and (**b**) summary plots of SI LP SBP tetramer$^{+}$ CD4$^{+}$ T cell number and phenotype in hCom2v-colonized or GF mice. (**c**) Representative flow cytometry and (**d**)

summary plots of LI LP α-Zein tetramer$^{+}$ CD4$^{+}$ T cell number and phenotype in hCom2v-colonized or GF mice. (**e**) Representative flow cytometry and (**f**) summary plots of SI LP α-Zein tetramer$^{+}$ CD4$^{+}$ T cell number and phenotype in hCom2v-colonized or GF mice. In panels b, d, and f, each dot represents an individual mouse, and horizontal lines indicate the mean; data compiled from two independent experiments and P values calculated by unpaired Welch's t-test. (**g**) Relative abundance of individual strains in the colon of the hCom2v-colonized mice used for tetramer staining, determined by metagenomic sequencing. Each dot represents an individual mouse ($n$ = 7), and only strains detected in at least one mouse are shown.

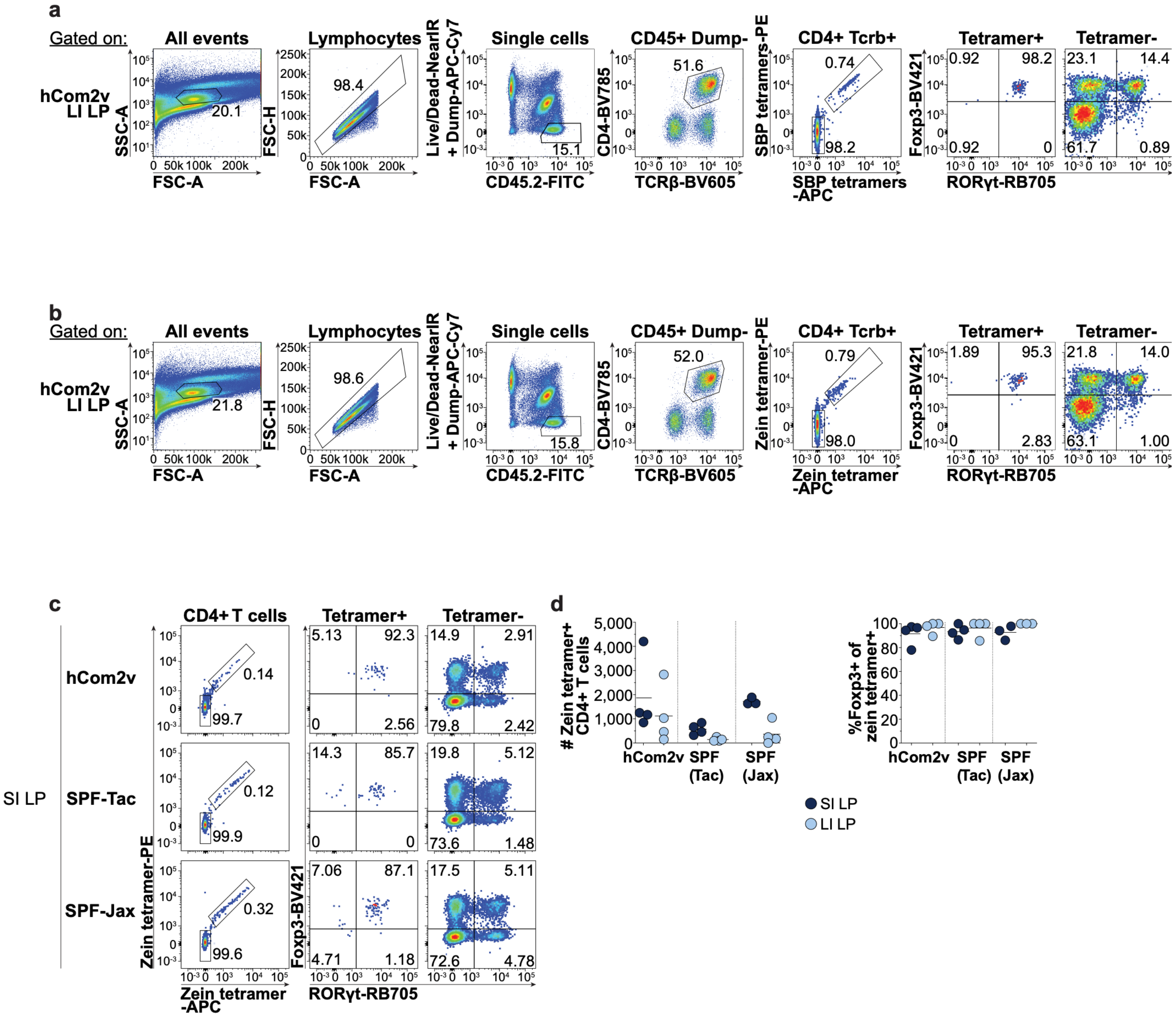


**Supplementary Figure 13: SBP and α-Zein tetramer gating strategy and detection of α-Zein-specific iT$_{regs}$ in SPF mice.** Representative flow cytometry plots depicting the gating strategy used for (**a**) SBP and (**b**) α-Zein tetramer staining throughout the manuscript. (**c**) Representative flow cytometry and (**d**) summary plots of SI LP and LI LP α-Zein tetramer$^+$ CD4$^+$ T cell number and phenotype in hCom2v-colonized, SPF-Tac, or SPF-Jax mice. In d, one SPF-Jax animal was excluded from the SI LP analysis due to failed lymphocyte recovery (negligible CD45$^+$ events). In summary plots, each dot represents an individual mouse and horizontal lines indicate the mean; data compiled from two independent experiments.

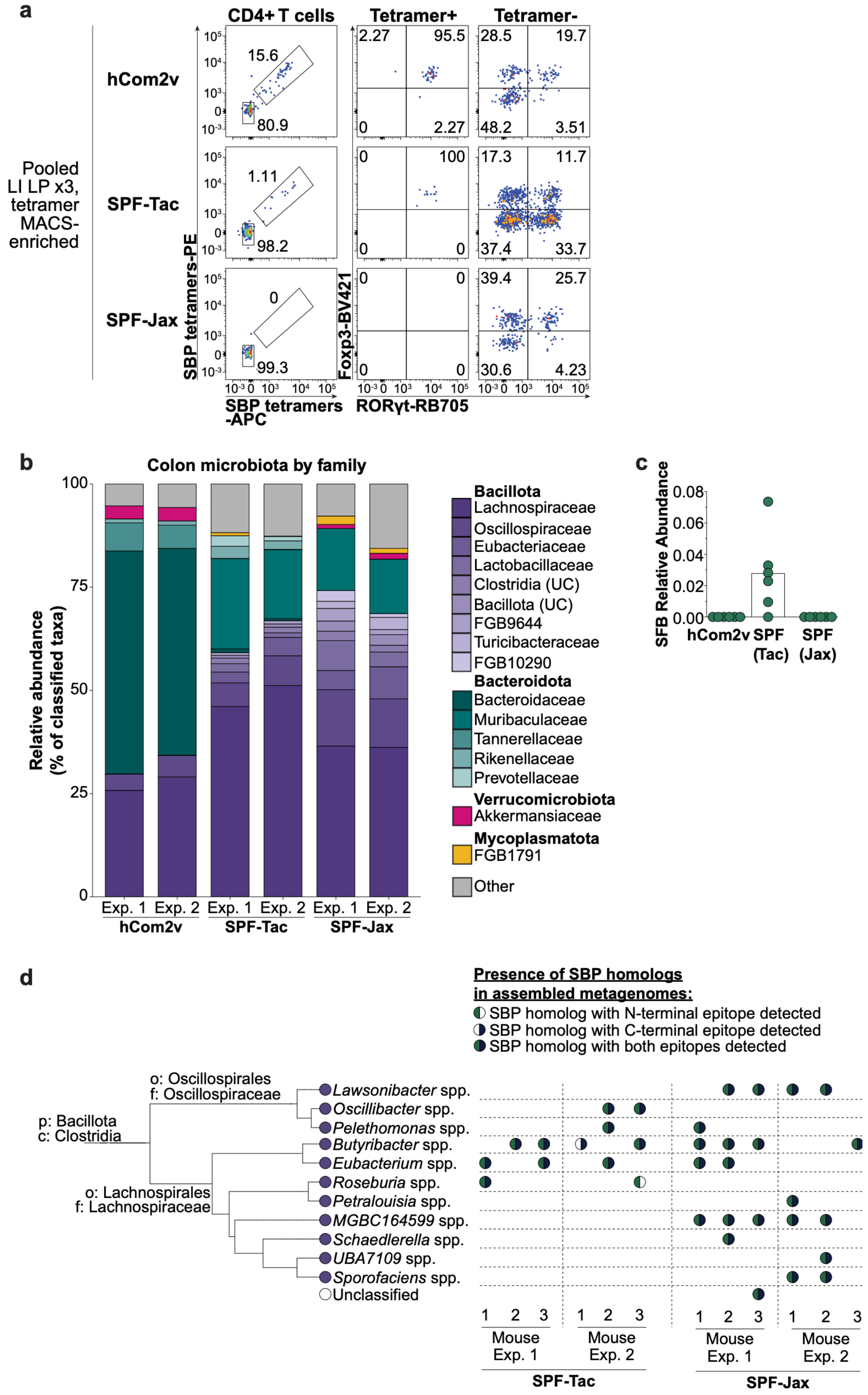


**Supplementary Figure 14: SBP-specific iT$_{regs}$ and microbiota-encoded SBP homologs in SPF mice.** (**a**) Representative flow cytometry plots of SBP tetramer$^{+}$ CD4$^{+}$ T cell number and phenotype from each group of three mice following tetramer enrichment. Data are from one of two independent experiments; a second experiment and summary plots for both appear in Fig. 3j-k. (**b**) Colonic microbiota composition across hCom2v, SPF-Tac, and SPF-Jax mice as determined by metagenomic sequencing. Each family is shown as a distinct shade of the color assigned to its phylum. Bars indicate the mean abundance for each family across the three mice used in each of two independent experiments. (**c**) Relative abundance of SFB in hCom2v, SPF-Tac, or SPF-Jax mice. Each dot represents an individual mouse; data compiled from two independent experiments. (**d**) Identification of SBP homologs in assembled metagenomes from SPF-Tac and SPF-Jax mice. Homologs were identified by tblastn; taxonomy was retrieved for the strain encoding the homolog and the amino acid sequence was aligned to the known SBP N- and C-terminal peptide epitopes. Circles and fill color indicate the presence of SBP homologs encoding the N- and/or C-terminal epitopes in the indicated strain across the three mice used in each of two independent experiments. The tree on the left depicts approximate phylogenetic relationships among identified strains encoding SBP homologs.

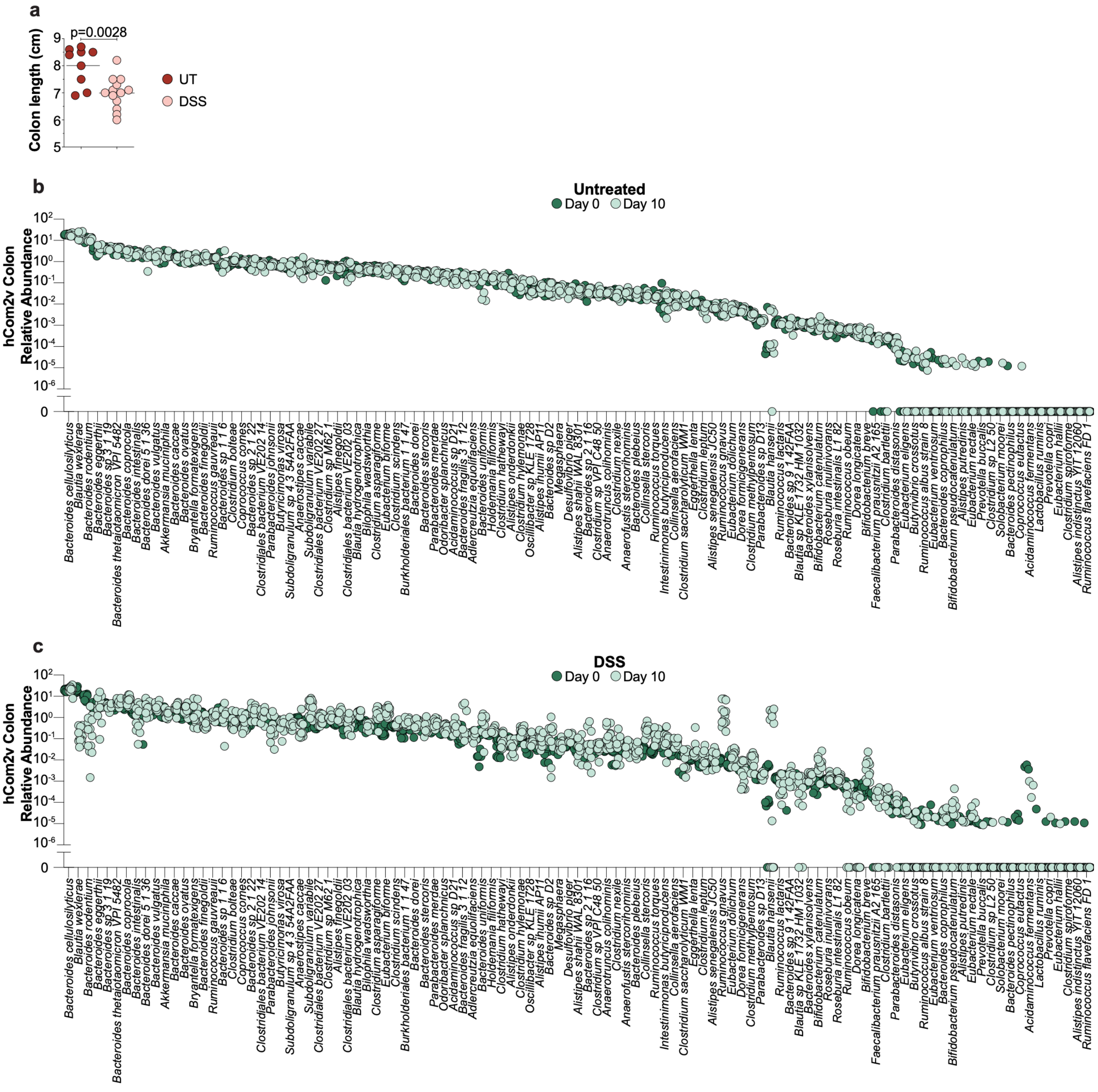


**Supplementary Figure 15: Colon shortening and hCom2v composition during acute DSS colitis.** (**a**) Colon length of untreated (UT) or DSS-treated hCom2v-colonized mice on experiment day 10. Each dot represents an individual mouse and horizontal lines indicate the mean; data compiled from three independent experiments. P value calculated by unpaired Welch's t-test. (**b**) Relative abundance of hCom2v strains in day 0 fecal pellets (pre-treatment) or day 10 colonic contents in untreated controls or (**c**) DSS-treated hCom2v-colonized mice, determined by metagenomic sequencing. Data compiled from three independent experiments. Each dot represents an individual mouse (*n* = 7-9 UT and 12-13 DSS; not all mice produced a pellet at day 0), and only strains detected in at least one mouse are shown.

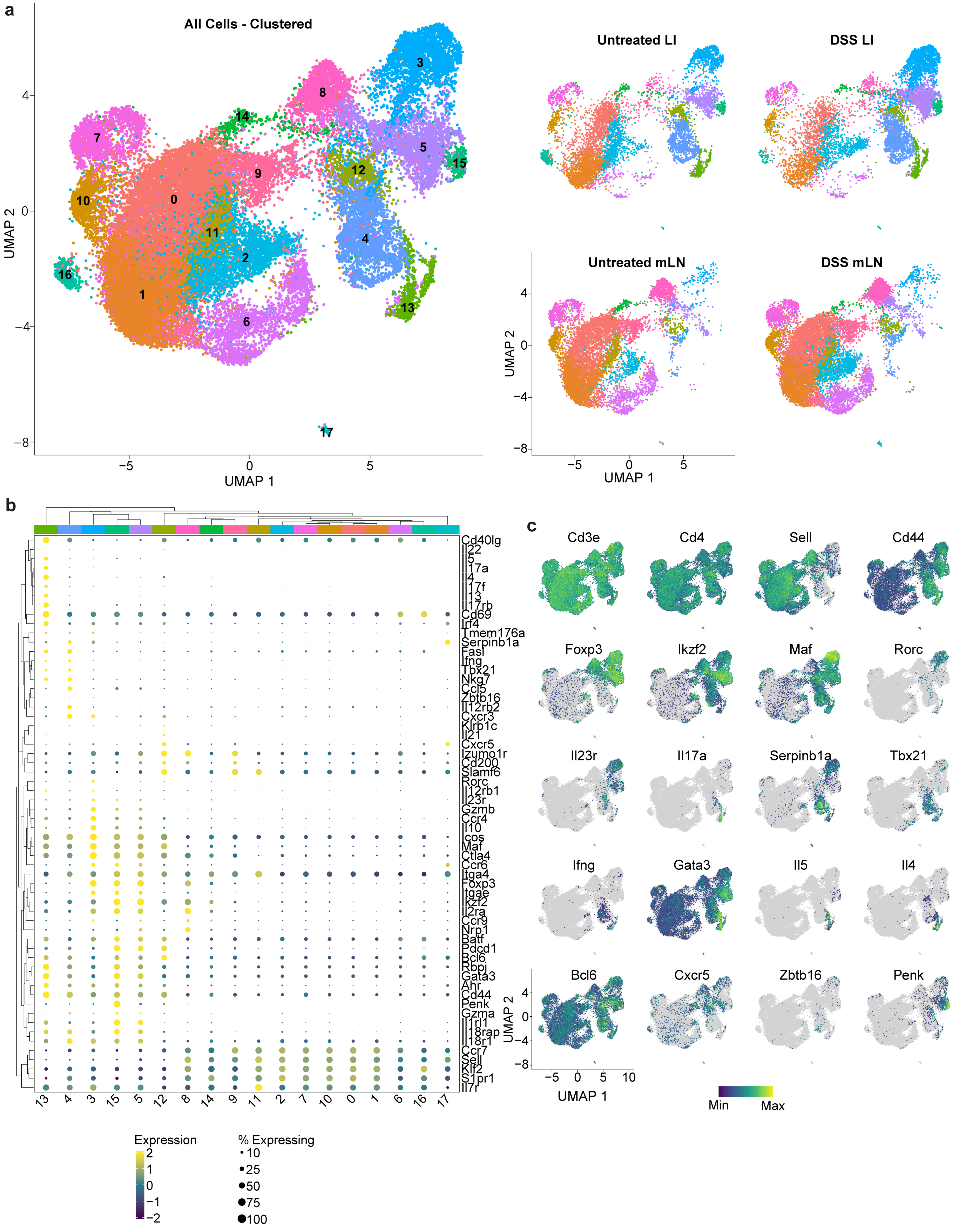
a
All Cells - Clustered
UMAP 1
UMAP 2
Untreated LI
DSS LI
Untreated mLN
DSS mLN
b
Cd40lg
Il22
Il5
Il17a
Il4
Il17f
Il13
Il17rb
Cd69
Irf4
Tmem176a
Serpinb1a
Fasl
Ifng
Tbx21
Nkg7
Ccl5
Zbtb16
Il12rb2
Cxcr3
Klrb1c
Il21
Cxcr5
Izumo1r
Cd200
Slamf6
Rorc
Il12rb1
Il23r
Gzmb
Ccr4
Il10
Icos
Maf
Ctla4
Ccr6
Itga4
Foxp3
Itgae
Ikzf2
Il2ra
Ccr9
Nrp1
Batf
Pdcd1
Bcl6
Rbpj
Gata3
Ahr
Cd44
Penk
Gzma
Il1rl1
Il18rap
Il18r1
Ccr7
Sell
Klf2
S1pr1
Il7r
Expression
% Expressing
c
Cd3e
Cd4
Sell
Cd44
Foxp3
Ikzf2
Maf
Rorc
Il23r
Il17a
Serpinb1a
Tbx21
Ifng
Gata3
Il5
Il4
Bcl6
Cxcr5
Zbtb16
Penk
Min
Max

**Supplementary Figure 16: Analysis of $CD4^+$ T cell clusters in DSS-treated hCom2v-colonized mice by scRNAseq.** (**a**) UMAP plot depicting unannotated $CD4^+$ T cell clusters identified by scRNAseq across all tissues combined (*left*) or subset by tissue and treatment group (untreated or DSS; *right*); cells from four untreated and seven DSS-treated mice were pooled. See Supplementary Fig. 17 for annotated clusters. (**b**) Clustered dot plot and (**c**) feature plots displaying expression patterns of selected marker genes used for cluster annotation. See Methods for multiplexing strategy and clustering parameters.

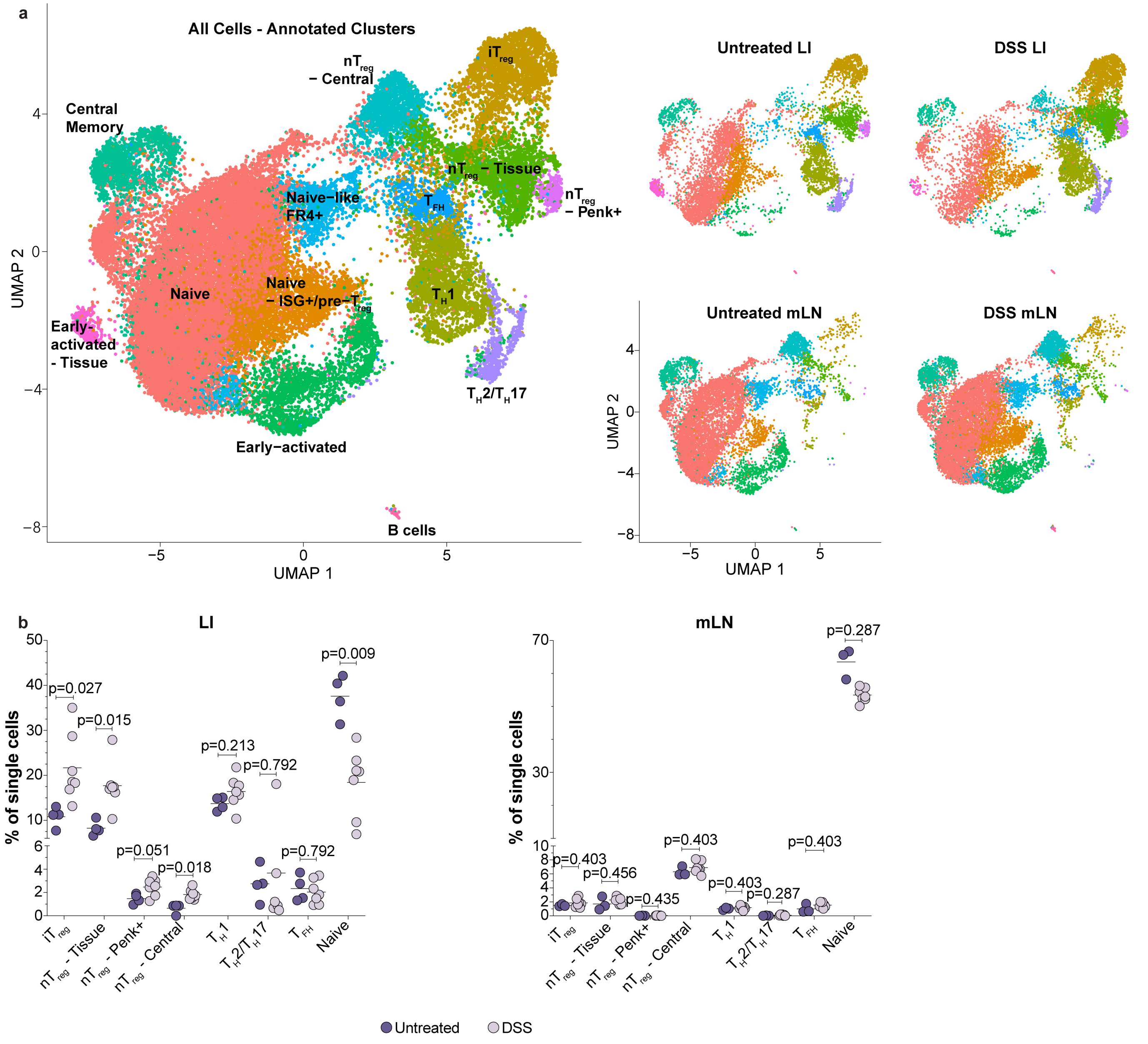


**Supplementary Figure 17: DSS treatment leads to expansion of multiple LI LP $T_{reg}$ subsets. (a)** UMAP plot depicting annotated CD4$^{+}$ T cell clusters identified by scRNAseq across all tissues combined (*left*) or subset by tissue and treatment group (untreated or DSS; *right*); cells from four untreated and seven DSS-treated mice were pooled. See Supplementary Fig. 16 for unannotated clusters and the marker genes used for annotation. (**b**) Percentage of cells in the indicated clusters of the LI LP (*left*) or mLN (*right*) of untreated or DSS-treated mice. Percentages were calculated per mouse after demultiplexing. Each dot represents an individual mouse and horizontal lines indicate the mean; one mouse is absent from the untreated mLN group due to negligible cell recovery. Data are from a single experiment. P values calculated by unpaired Welch's t-test and adjusted for multiple comparisons with the Benjamini-Hochberg method.

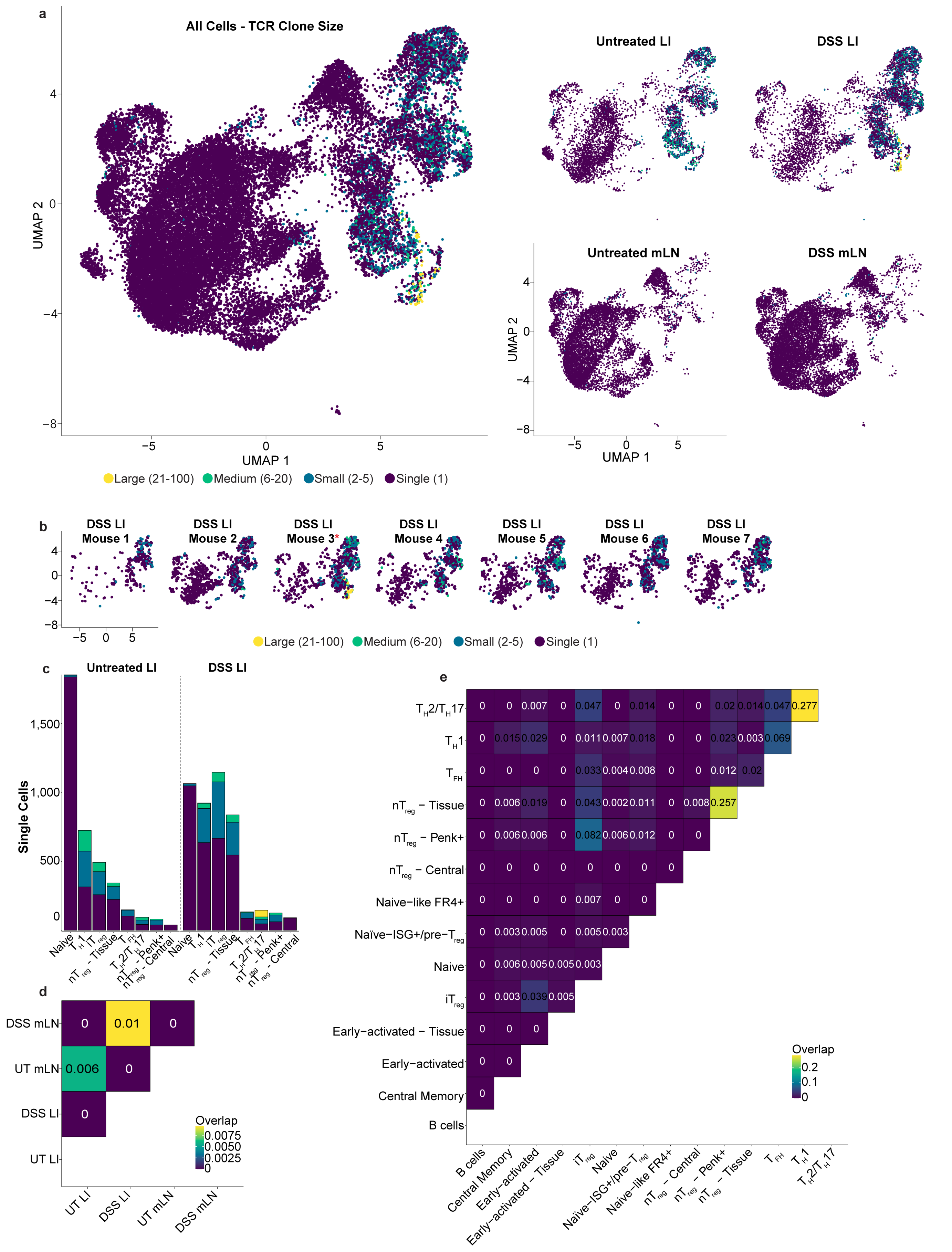
a
All Cells - TCR Clone Size
Untreated LI
DSS LI
Untreated mLN
DSS mLN
UMAP 1
UMAP 2
Large (21-100)
Medium (6-20)
Small (2-5)
Single (1)
b
DSS LI Mouse 1
DSS LI Mouse 2
DSS LI Mouse 3*
DSS LI Mouse 4
DSS LI Mouse 5
DSS LI Mouse 6
DSS LI Mouse 7
c
Untreated LI
DSS LI
Single Cells
d
DSS mLN
UT mLN
DSS LI
UT LI
Overlap
e
Overlap
B cells
Central Memory
Early−activated
Early−activated − Tissue
Naive
Naïve−ISG+/pre−Treg
Naive−like FR4+
nTreg − Central
nTreg − Penk+
nTreg − Tissue
TFH
TH1
TH2/TH17

**Supplementary Figure 18: TCR clonal architecture in DSS-treated hCom2v-colonized mice.** (**a**) UMAP plot with dots colored by TCR clone size (as determined by scTCRseq) across all tissues combined (*left*) or subset by tissue and treatment group (untreated or DSS; *right*); cells from four untreated and seven DSS-treated mice were pooled. (**b**) UMAP plots depicting TCR clone size in each DSS-treated mouse. The large clonal expansions in the $T_H2/T_H17$ cluster in panel a derive predominantly from a single mouse (Mouse 3, red asterisk), which was retained in all analyses. (**c**) Number of TCR clonotypes with the indicated clone sizes in each cluster. (**d**) TCR clonal overlap between tissues and (**e**) between clusters in the LI LP; fill color indicates the clonotype overlap coefficient. See Supplementary Fig. 17a for annotated scRNAseq clusters.

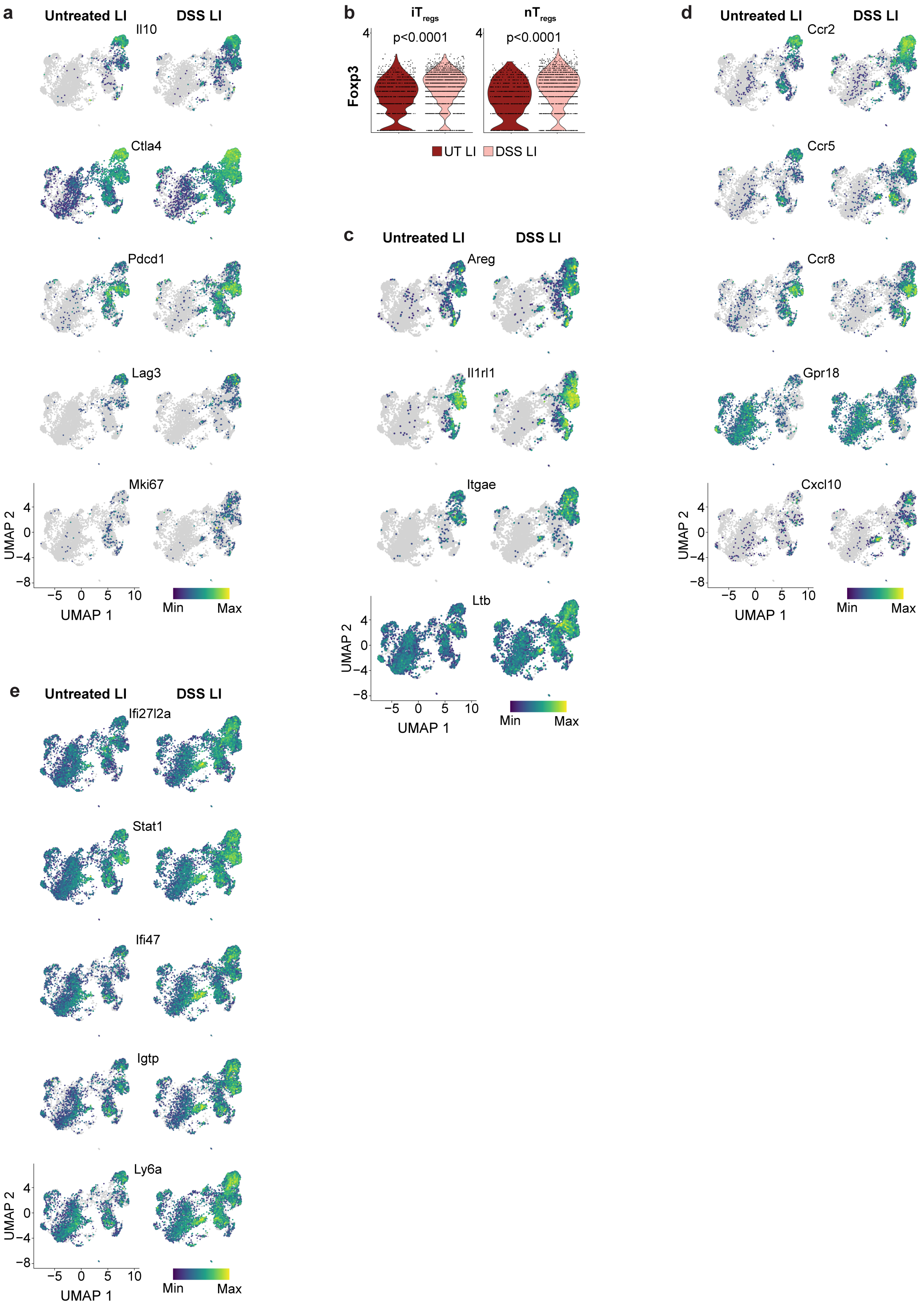
a
Untreated LI
DSS LI
Il10
Ctla4
Pdcd1
Lag3
Mki67
UMAP 2
UMAP 1
Min
Max
b
iTregs
nTregs
p<0.0001
p<0.0001
Foxp3
UT LI
DSS LI
c
Untreated LI
DSS LI
Areg
Il1rl1
Itgae
Ltb
d
Untreated LI
DSS LI
Ccr2
Ccr5
Ccr8
Gpr18
Cxcl10
e
Untreated LI
DSS LI
Ifi27l2a
Stat1
Ifi47
Igtp
Ly6a

**Supplementary Figure 19: Gene expression in DSS-treated hCom2v-colonized mice.** (**a**) Feature plots displaying expression patterns of selected genes associated with immunosuppressive function or proliferation. (**b**) Violin plots displaying *Foxp3* gene expression levels in the LI LP $iT_{reg}$ (*left*) or $nT_{reg}$ (*right*) clusters of untreated or DSS-treated mice; adjusted P values calculated by pseudobulk analysis using individual mice as replicates after demultiplexing (see Methods). (**c**) Feature plots displaying expression patterns of selected genes associated with tissue repair, (**d**) chemotaxis, or (**e**) an interferon-stimulated gene signature. All plots depict gene expression in LI LP cells pooled from four untreated and seven DSS-treated mice; data are from a single experiment.

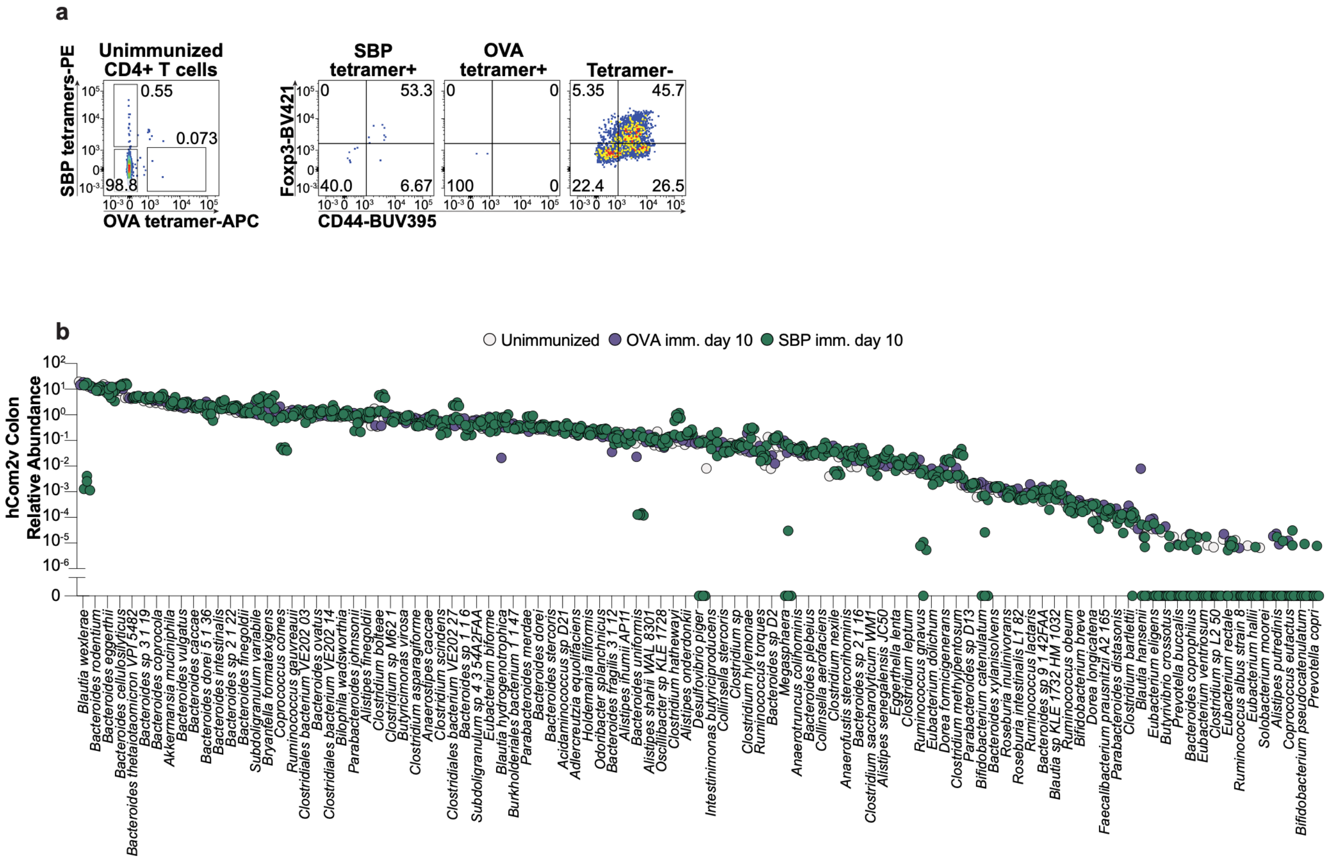


**Supplementary Figure 20: Tetramer staining in unimmunized controls and hCom2v community composition in immunized mice.** (**a**) Representative flow cytometry plots depicting SBP and OVA tetramer staining following tetramer enrichment from pooled spleen and LNs from unimmunized hCom2v-colonized mice (*left*) and Foxp3 and CD44 staining of tetramer$^{+}$ or tetramer$^{-}$ populations (*right*); see Fig. 4f for summary plots and replicate information. (**b**) Relative abundance of hCom2v strains in colonic contents from SBP- or OVA-immunized or unimmunized mice on day 10, determined by metagenomic sequencing. Data compiled from two independent experiments. Each dot represents an individual mouse ($n$ = 3 unimmunized, 3 OVA imm., and 7 SBP imm.), and only strains detected in at least one mouse are shown.

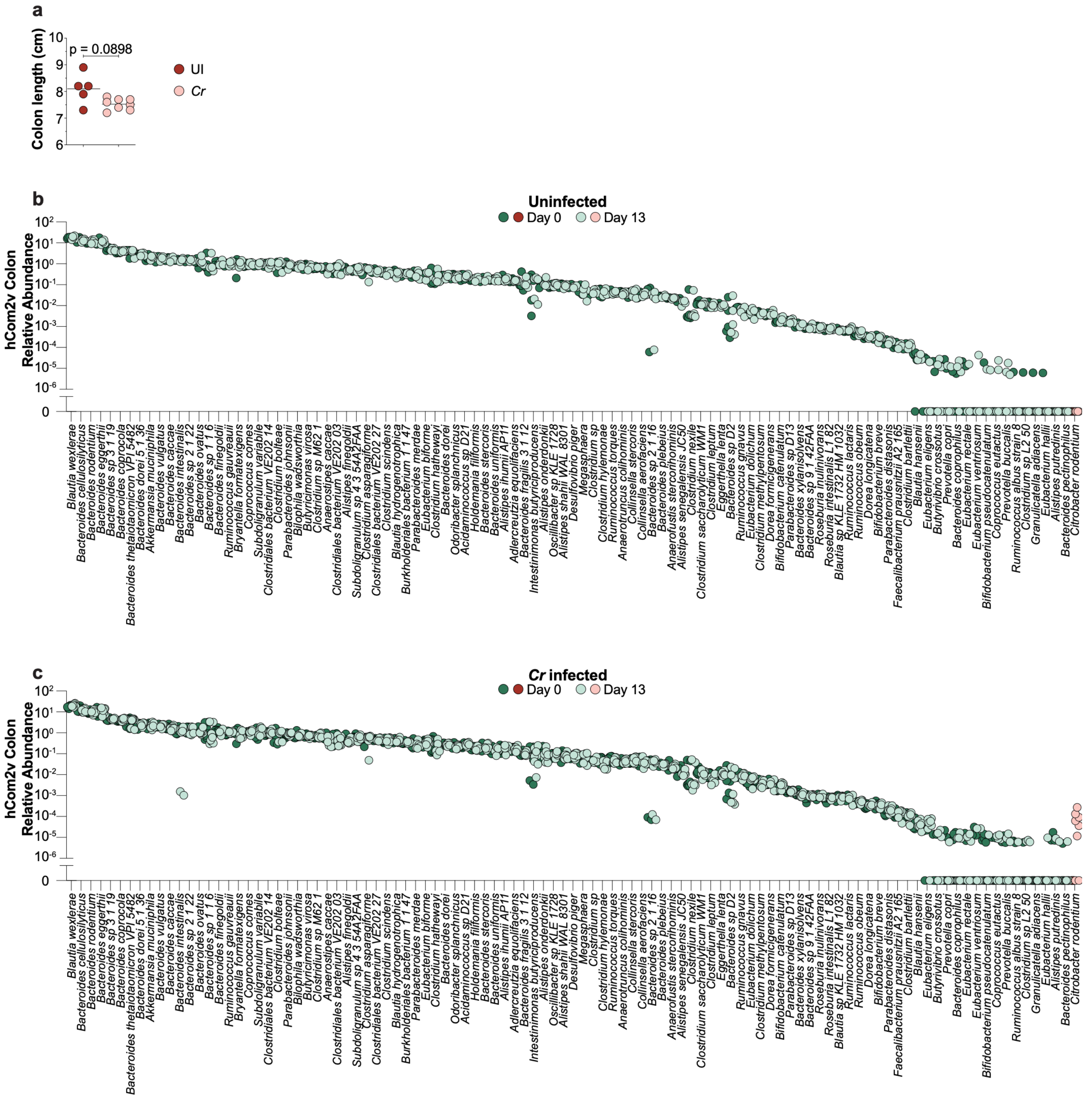


**Supplementary Figure 21: Colon length and hCom2v composition during *C. rodentium* infection.** (**a**) Colon length of uninfected (UI) or *C. rodentium*-infected (*Cr*) hCom2v-colonized mice on experiment day 13. Each dot represents an individual mouse and horizontal lines indicate the mean; data compiled from two independent experiments. P value calculated by unpaired Welch's t-test. (**b**) Relative abundance of hCom2v strains and *C. rodentium* in day 0 fecal pellets (pre-infection) or day 13 colonic contents in uninfected controls or (**c**) *C. rodentium*-infected hCom2v-colonized mice, determined by metagenomic sequencing. Data compiled from two independent experiments. Each dot represents an individual mouse ($n$ = 5 UI and 8 *Cr*), and only strains detected in at least one mouse are shown.

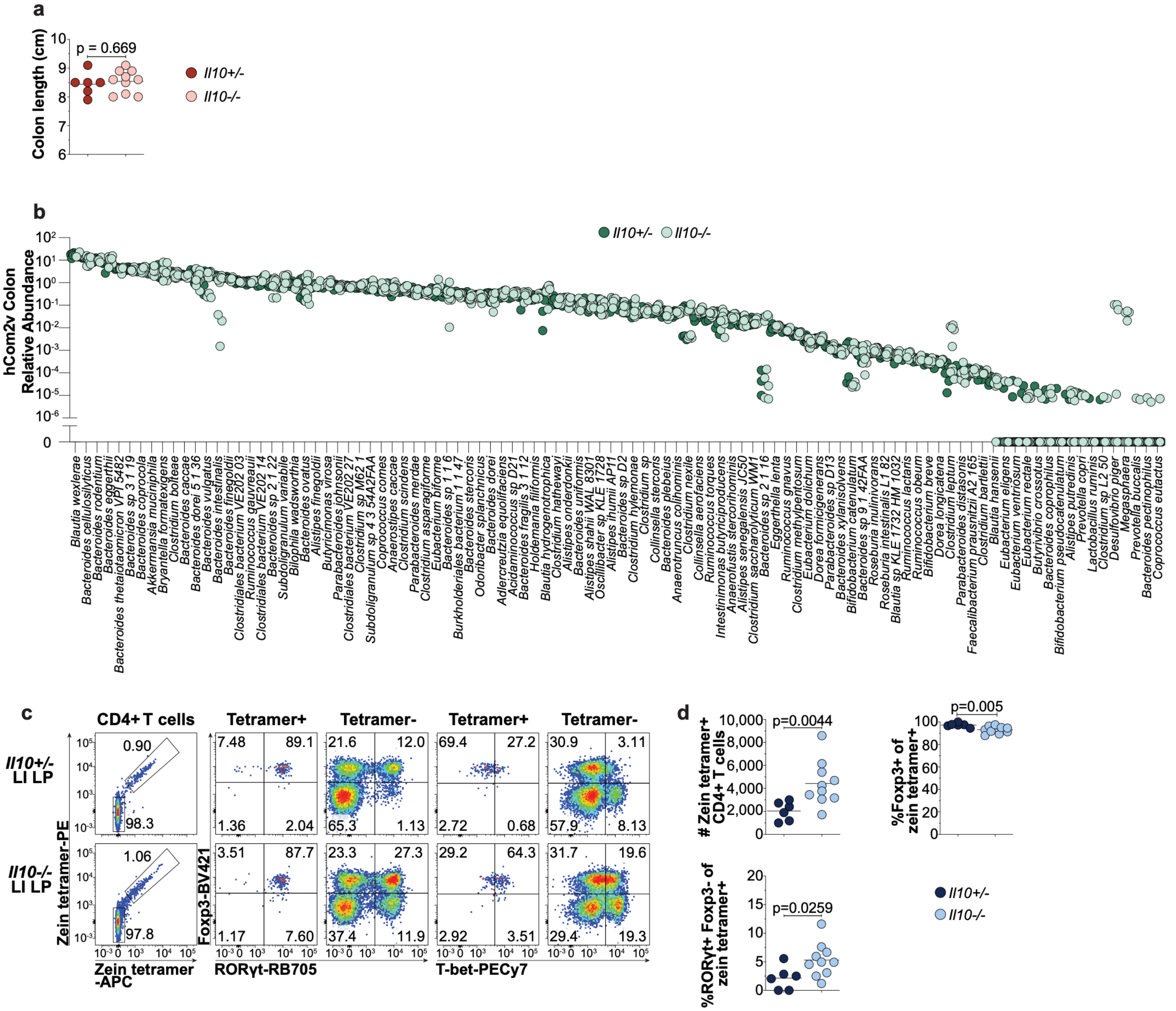


**Supplementary Figure 22: Colon length, hCom2v composition, and α-Zein-specific responses in *Il10*$^{-/-}$ mice.** (**a**) Colon length of *Il10*$^{-/-}$ or *Il10*$^{+/-}$ hCom2v-colonized mice. Each dot represents an individual mouse and horizontal lines indicate the mean; data compiled from two independent experiments. P value calculated by unpaired Welch's t-test. (**b**) Relative abundance of hCom2v strains in the *Il10*$^{-/-}$ or *Il10*$^{+/-}$ mice analyzed in c-d and Fig. 4k-l at ~12 weeks of age, determined by metagenomic sequencing. Data compiled from two independent experiments. Each dot represents an individual mouse ($n$ = 6 *Il10*$^{+/-}$ and 10 *Il10*$^{-/-}$), and only strains detected in at least one mouse are shown. (**c**) Representative flow cytometry and (**d**) summary plots of α-Zein tetramer$^{+}$ CD4$^{+}$ T cell number and phenotype in *Il10*$^{-/-}$ or *Il10*$^{+/-}$ mice; data compiled from two independent experiments. In summary plots, each dot represents an individual mouse and horizontal lines indicate the mean; P values calculated by unpaired Welch's t-test.

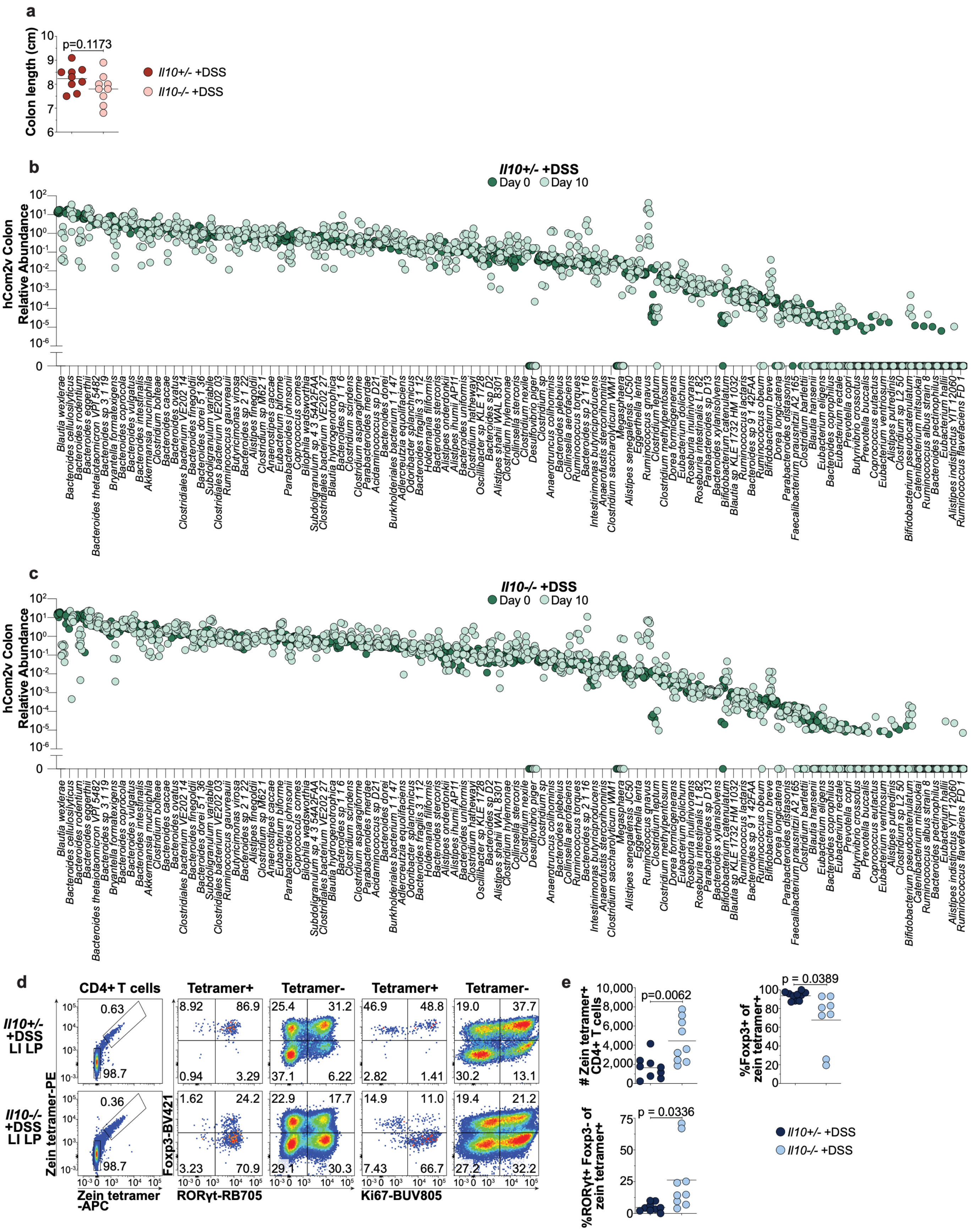
a
Colon length (cm)
p=0.1173
Il10+/- +DSS
Il10-/- +DSS
b
Il10+/- +DSS
Day 0
Day 10
hCom2v Colon Relative Abundance
c
Il10-/- +DSS
d
CD4+ T cells
Tetramer+
Tetramer-
Il10+/- +DSS LI LP
Il10-/- +DSS LI LP
Zein tetramer-PE
Zein tetramer -APC
Foxp3-BV421
RORγt-RB705
Ki67-BUV805
e
### Zein tetramer+ CD4+ T cells
p=0.0062
%Foxp3+ of zein tetramer+
p = 0.0389
%RORγt+ Foxp3- of zein tetramer+
p = 0.0336

**Supplementary Figure 23: Colon length, hCom2v composition, and α-Zein-specific responses in *Il10$^{-/-}$* mice during acute DSS colitis.** (**a**) Colon length of DSS-treated *Il10$^{-/-}$* or *Il10$^{+/-}$* hCom2v-colonized mice on experiment day 10. Each dot represents an individual mouse and horizontal lines indicate the mean; data compiled from two independent experiments. P value calculated by unpaired Welch's t-test. (**b**) Relative abundance of hCom2v strains in DSS-treated *Il10$^{+/-}$* or (**c**) *Il10$^{-/-}$* mice in day 0 fecal pellets (pre-treatment) or day 10 colonic contents, determined by metagenomic sequencing. Data compiled from two independent experiments. Each dot represents an individual mouse ($n$ = 9 *Il10$^{+/-}$* +DSS and 9 *Il10$^{-/-}$* +DSS), and only strains detected in at least one mouse are shown. (**d**) Representative flow cytometry and (**e**) summary plots of α-Zein tetramer$^{+}$ CD4$^{+}$ T cell number and phenotype in DSS-treated *Il10$^{-/-}$* or *Il10$^{+/-}$* mice; data compiled from two independent experiments. In summary plots, each dot represents an individual mouse and horizontal lines indicate the mean; P values calculated by unpaired Welch's t-test.

**Supplementary Table 1: Golden Gate cloning and TCR sequences with QC outcomes.** Sheet 1 ("Sequences for Golden Gate") lists whole-plasmid sequences of the Destination and Middle Insert Vectors used for Golden Gate assembly and 5' and 3' attachment sequences for cloning *TRA* or *TRB* variable regions. Sheet 2 ("TCR sequences and QC") lists all TCRs selected for this study with their sequences, the synthesized fragments used for Golden Gate assembly (which carry silent mutations eliminating internal SapI sites where present), the whole-plasmid sequence of each cloned TCR expression vector, and QC outcomes (see Methods for TCR selection and QC criteria). In TCR names, "Colon" denotes the LI LP. TCR numbering is non-consecutive, and absent numbers correspond to clones not analyzed in this study.

**Supplementary Table 2: Complete TCR reactivity data.** Reactivity data for all TCRs against the 128 stimuli in the primary screen. Values indicate IL-2 (in pg/mL) as measured by AlphaLISA, and values ≥ 500 pg/mL for non-control stimuli were confirmed in a secondary hit validation assay (see Methods). In TCR names, "Colon" denotes the LI LP. Some TCRs were shared between populations; these were assayed on the first occurrence, with the same data reported in each population for which they met selection criteria, with the assayed TCR number, plate, and well identifying the clone used (see Methods and Supplementary Table 1). Each sheet lists reactivity data for all TCRs included in the indicated population, together with the assigned peptide epitope where identified. "Population summaries" lists the number of TCRs in each panel scored in each reactivity category.

**Supplementary Table 3: hCom2v strains and growth media.** Sheet 1 ("hCom2v strains and media") lists the 116 strains in hCom2v with TY number identifiers and preferred liquid or solid growth media. The remaining sheets give recipes for liquid growth media made in-house. Modified PYG, Ethanoligenes, and Bifidobacterium media recipes were from the DSMZ.
